# The *Cryptococcus neoformans* STRIPAK complex controls genome stability, sexual development, and virulence

**DOI:** 10.1101/2024.08.06.606879

**Authors:** Patricia P. Peterson, Jin-Tae Choi, Ci Fu, Leah E. Cowen, Sheng Sun, Yong-Sun Bahn, Joseph Heitman

## Abstract

The eukaryotic serine/threonine protein phosphatase PP2A is a heterotrimeric enzyme composed of a scaffold A subunit, a regulatory B subunit, and a catalytic C subunit. Of the four known B subunits, the B’’’ subunit (known as striatin) interacts with the multi-protein striatin-interacting phosphatase and kinase (STRIPAK) complex. Orthologs of STRIPAK components were identified in *Cryptococcus neoformans*, namely PP2AA/Tpd3, PP2AC/Pph22, PP2AB/Far8, STRIP/Far11, SLMAP/Far9, and Mob3. Structural modeling, protein domain analysis, and detected protein-protein interactions suggest *C. neoformans* STRIPAK is assembled similarly to the human and fungal orthologs. Here, STRIPAK components Pph22, Far8, and Mob3 were functionally characterized. Whole-genome sequencing revealed that mutations in STRIPAK complex subunits lead to increased segmental and chromosomal aneuploidy, suggesting STRIPAK functions in maintaining genome stability. We demonstrate that *PPH22* is a haploinsufficient gene: heterozygous *PPH22/pph22*Δ mutant diploid strains exhibit defects in hyphal growth and sporulation and have a significant fitness disadvantage when grown in competition against a wild-type diploid. Deletion mutants *pph22*Δ*, far8*Δ, and *mob3*Δ exhibit defects in mating and sexual differentiation, including impaired hyphae, basidia, and basidiospore production. Loss of either *PPH22* or *FAR8* in a haploid background leads to growth defects at 30⁰C, severely reduced growth at elevated temperature, abnormal cell morphology, and impaired virulence. Additionally, *pph22*Δ strains frequently accumulate suppressor mutations that result in overexpression of another putative PP2A catalytic subunit, *PPG1.* The *pph22*Δ and *far8*Δ mutants are also unable to grow in the presence of the calcineurin inhibitors cyclosporine A or FK506, and thus these mutations are synthetically lethal with loss of calcineurin activity. Conversely, *mob3*Δ mutants display increased thermotolerance, capsule production, and melanization, and are hypervirulent in a murine infection model. Taken together, these findings reveal that the *C. neoformans* STRIPAK complex plays an important role in genome stability, vegetative growth, sexual development, and virulence in this prominent human fungal pathogen.

**Author summary:** This study focused on a highly conserved protein signaling complex known as STRIPAK, which is important for various developmental processes in fungi and humans. By investigating the functions of this complex in *Cryptococcus neoformans*, it was discovered to play crucial roles in maintaining genome stability, sexual development, and pathogenesis. In particular, mutations in the genes encoding two subunits of the STRIPAK complex were found to lead to significant defects, including abnormal growth and cell morphology, compromised stress response, and impaired virulence. Interestingly, mutation of a third STRIPAK complex subunit resulted in hypervirulence, characterized by increased thermotolerance, enhanced production of melanin pigment and polysaccharide capsule, and reduced survival in infected animals. Our findings reveal that the STRIPAK complex is an important regulator of growth and virulence of *C. neoformans*, highlighting its potential as a target for therapies aimed at combating fungal infections. This work furthers understanding of how the STRIPAK complex functions in *Cryptococcus*, and also in other organisms including humans, where related protein phosphatase complexes govern key cellular processes.

## Introduction

Eukaryotic organisms utilize dynamic signaling networks to respond and adapt to changes in their internal and external environments. Sensing of environmental stimuli triggers a cascade of downstream events, eliciting a coordinated cellular response tightly orchestrated by interconnected signal transduction pathways. The function and activity of signaling components within these pathways are modulated through post-translational modifications, including protein phosphorylation, which is governed by the balanced actions of kinases and phosphatases. Phosphorylation and dephosphorylation of target proteins by kinases and phosphatases are coordinated by finely-tuned mechanisms to ensure signaling pathways are turned on and off as needed, maintaining cellular homeostasis. Among eukaryotic phosphatases, protein phosphatase 2A (PP2A) plays a pivotal role in governing numerous cellular processes, including cell cycle progression, proliferation, apoptosis, metabolism, and stress responses [1].

The PP2A regulatory B subunit striatin is a key component of the striatin-interacting phosphatase and kinase (STRIPAK) complex [2–5]. First identified in mammals, striatin proteins act as scaffolds to assemble the other STRIPAK subunits, forming a large, multifunctional signaling complex. In addition to the PP2A holoenzyme, STRIPAK is comprised of a striatin-interacting protein (STRIP), monopolar spindle-one-binder related protein (Mob3/Phocein), sarcolemmal membrane-associated protein (SLMAP), small coiled-coil protein (SIKE), cerebral cavernous malformation protein (CCM3), and a germinal center kinase (GCKIII) [3]. Orthologs of the mammalian STRIPAK complex have been identified in many fungi, including *Sordaria macrospora, Neurospora crassa, Saccharomyces cerevisiae, Schizosaccharomyces pombe, Aspergillus nidulans, Fusarium graminearum,* and several other ascomycete species [6–24].

In mammals and fungi, the STRIPAK complex connects signal transduction pathways to regulate numerous aspects of cell growth and development, such as TORC2 signaling and actin cytoskeleton remodeling in *S. cerevisiae,* and the Hippo pathway and regulation of tissue growth in humans [14, 25, 26]. In fungi, STRIPAK plays a critical role in morphogenesis and sexual development, including control of cell fusion, hyphal elongation, fruiting body formation, fertility, nuclear division, and sporulation [27, 28]. The *S. cerevisiae* STRIPAK counterpart, the Far complex, is implicated in pheromone-induced cell cycle arrest during mating, acts as an antagonist to TORC2 at the endoplasmic reticulum (ER), and inhibits mitophagy at the mitochondrial membrane [14, 29–32]. In *N. crassa*, homologs of STRIP and Striatin act on two MAP kinase pathways, downstream of the cell wall integrity (CWI) and pheromone response pathways, modulating fungal self-signaling and developmental morphogenesis [8]. The STRIPAK complex also governs virulence in plant fungal pathogens including *Magnaporthe oryzae* and several *Fusarium* species [12, 13, 23, 33, 34].

Although the STRIPAK complex has been characterized in pathogenic and non-pathogenic ascomycetes, it has not yet been elucidated in a basidiomycete or notable human fungal pathogen. The basidiomycetous yeast and opportunistic pathogen *Cryptococcus neoformans* is a clinically relevant and genetically tractable model for studying the molecular mechanisms underlying fungal pathogenesis in humans. *C. neoformans* infections occur following inhalation of spores or desiccated yeast cells from the environment and are more prevalent in immunocompromised hosts. Systemic cryptococcosis can lead to lethal meningoencephalitis, accounting for approximately 20% of HIV/AIDS-related deaths annually, thus representing a significant burden of global fungal diseases [35–37]. *C. neoformans* possesses virulence traits necessary to cause disease, including melanization, extracellular polysaccharide capsule production, resistance to oxidative stress, and thermotolerance [38–40]. Understanding the regulatory pathways underlying *C. neoformans* virulence is critical for developing targeted therapies against cryptococcosis.

Due to the key roles of STRIPAK in controlling cell growth, developmental processes, and pathogenicity in other fungi, this study aimed to functionally characterize the cellular functions of the *C. neoformans* STRIPAK complex. We identified protein orthologs for six STRIPAK complex subunits, three of which, Pph22 (PP2A catalytic subunit), Far8 (regulatory B¢¢¢ subunit), and Mob3, were characterized. Genome sequencing of *pph22*Δ, *far8*Δ, and *mob3*Δ deletion mutants revealed frequent segmental and whole chromosomal aneuploidy, suggesting a role in maintaining genome stability. We demonstrate that *PPH22* is not an essential gene, though the loss of one functional copy leads to haploinsufficiency in a heterozygous diploid strain, and the *pph22*Δ mutation incurs a significant fitness cost in a haploid. Similar to findings in other fungi, *C. neoformans* STRIPAK is important for mating, hyphal formation, and sporulation. Both Pph22 and Far8 are crucial for high-temperature growth, stress response, and virulence. Surprisingly, *mob3*Δ mutants exhibited increased thermotolerance, melanization, and capsule production, resulting in hypervirulence in a murine model of *C. neoformans* infection. Taken together, these studies shed light on the organization and function of the STRIPAK complex in *C. neoformans* and provide a foundation for future identification of its cellular targets to understand the mechanisms mediating its various roles.

## Results

### Identification of *Cryptococcus neoformans* STRIPAK complex components

To identify the *C. neoformans* STRIPAK complex, BLAST searches (blastp) were performed with known *Homo sapiens* and *S. cerevisiae* STRIPAK components and the *C. neoformans* H99 genome (Fig 1A). Significant alignments were found for the protein phosphatase 2A catalytic C subunit (CNAG_02177/Pph22), the scaffold A subunit (CNAG_07914/Tpd3), the regulatory B¢¢¢ subunit (CNAG_00073/Far8), the striatin-interacting protein (CNAG_00008/Far11), the tail-anchor domain protein (CNAG_04838/Far9), and the striatin-associated protein (CNAG_04629/Mob3). No ortholog of the coiled-coil domain protein (Far3/7 in *S. cerevisiae,* SIKE in *H. sapiens,* Sci1 in *S. macrospora,* SipB in *A. nidulans,* and Csc4 in *S. pombe*) was identified, possibly due to its small size and low sequence similarity [16]. BLAST analysis of the STRIPAK-associated kinase with the sequences of the three mammalian kinases Stk24, Stk25, and Stk26 [3], which are part of the GCKIII family of kinases, produced three significant alignments in *C. neoformans* (CNAG_03290, CNAG_00405, and CNAG_05274). Reciprocal BLAST analyses confirmed an orthologous relationship between the proteins. With the exception of *PPH22* [41], none of the *C. neoformans* STRIPAK complex components were annotated on FungiDB (https://fungidb.org/fungidb) and thus are referred to here based on *S. cerevisiae* or *H. sapiens* nomenclature.

**Fig 1.**
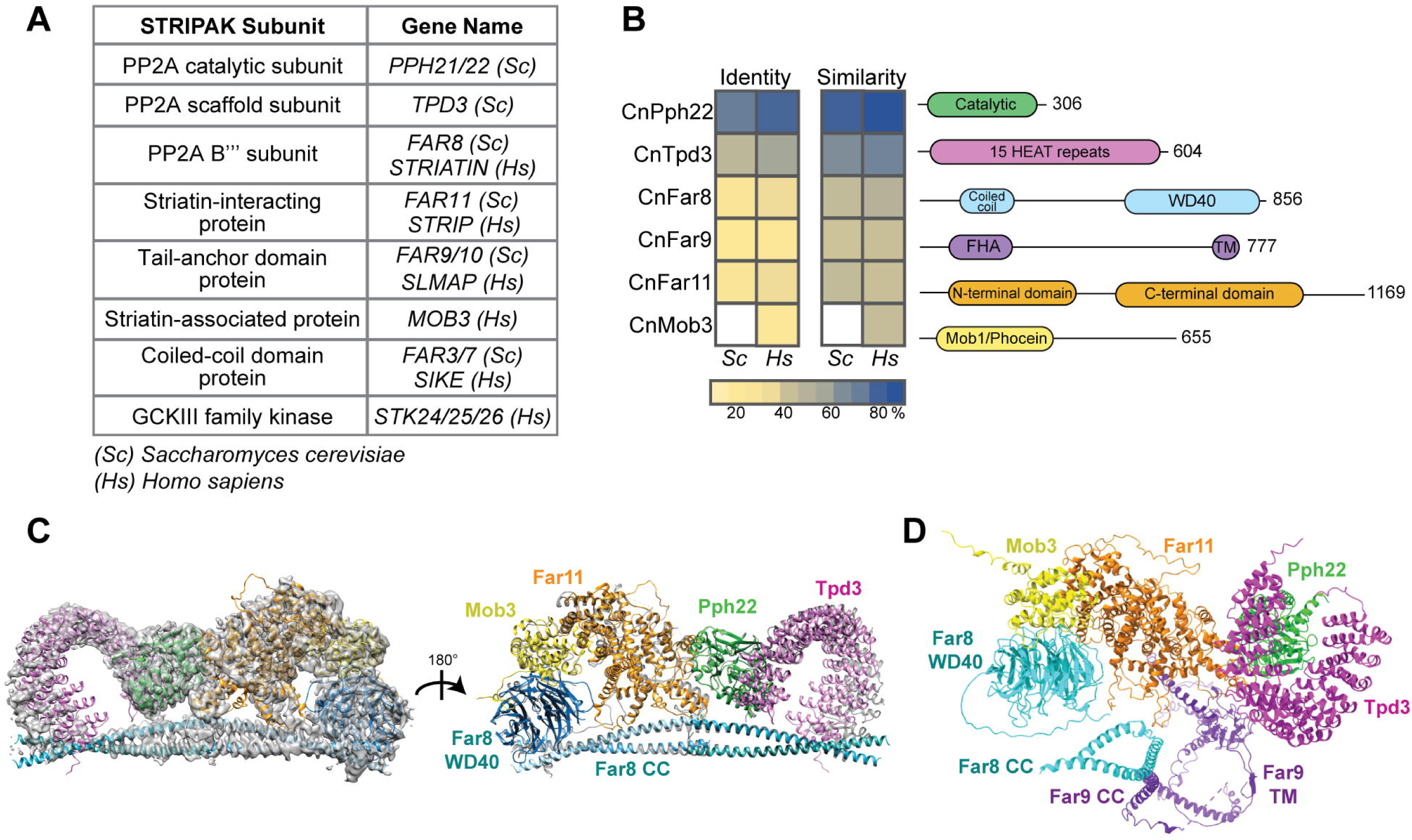
Conservation analysis of the *C. neoformans* STRIPAK complex. (A) Table listing STRIPAK complex subunits used in BLAST analysis to identify protein orthologs in *C. neoformans*. (B) Sequence alignment of STRIPAK protein orthologs in *C. neoformans, S. cerevisiae*, and *H. sapiens*. Conserved protein domains are labeled in the schematic diagram: HEAT (Huntingtin, elongation factor 3, protein phosphatase 2A, TOR1), FHA (forkhead-associated domain), TM (transmembrane domain), and Mob1 (monopolar spindle-one-binder protein). (C) The *Cryptococcus* STRIPAK complex superimposed on the electron density map and ribbon model of the human STRIPAK complex using the program ChimeraX. The human complex is shown in grey, and the corresponding *C. neoformans* proteins are labeled in color. (D) AlphaFold structure prediction of CnSTRIPAK, with dotted lines representing pseudobonds.

Multi-sequence alignments revealed the *C. neoformans* STRIPAK components share 23%-86% identity and 42%-93% similarity with the *S. cerevisiae* and *H. sapiens* orthologs, with greater homology to the human than the yeast proteins (Fig 1B). Mob3 from *C. neoformans* only aligned with the human protein, as *S. cerevisiae* lacks Mob3. The STRIPAK components also exhibit highly conserved structures and domain architecture [28] (Fig 1B). For example, CnTpd3 contains 15 tandem HEAT (huntingtin-elongation-A subunit-TOR) repeats, which mediate interactions with the B and C subunits to form the PP2A holoenzyme [1]. Eukaryotic striatin proteins share a coiled-coil domain and a WD40 repeat domain, forming a β-propeller. Human striatin interacts with the PP2A A and C subunits via the coiled-coil domain [42, 43]. The tail-anchor domain protein CnFar9 contains a conserved FHA (forkhead-associated) domain and a small, hydrophobic transmembrane domain at its C-terminus. In fungi and mammals, mutations in the tail-anchor domain of Far9/SLMAP lead to changes in their subcellular localization, and the membrane association of Far9 with ER and mitochondria is important for its functions [14, 31, 44].

Due to the evolutionarily conserved nature of the *C. neoformans* STRIPAK complex protein sequences and domains (Fig 1B), it was predicted that the complex would also have a similar three-dimensional structure and organization. Recently, the core of the human STRIPAK complex was resolved at high resolution by cryo-EM [43]. The *C. neoformans* proteins were aligned with the model of the human STRIPAK core with the program ChimeraX [45] (EMD-22650, PDB-7k36) (Fig 1C). The predicted and known protein structures are remarkably similar. The *C. neoformans* STRIPAK complex was predicted with AlphaFold2 multimer [46] (Fig 1D). This structure prediction included Far9, which has not been identified in the cryo-EM model or crystal structures of human STRIPAK [42, 43]. The predicted CnSTRIPAK complex model showed a linear arrangement of the proteins Mob3-Far11-Pph22-Tpd3, similar to the human complex. The Far8 WD40 domain lies in close proximity to Mob3, and the coiled-coil domain is at the base of the complex. The middle of the Far9 protein contains a large coiled-coil domain (approximately 135 amino acids) that interacts closely with the coiled-coil of Far8, stabilizing the core complex. Taken together, these results suggest that the key components of the *C. neoformans* STRIPAK complex are significantly conserved in their domain architecture, three-dimensional structure, and assembly. Our data also suggest the CnSTRIPAK complex is more similar to human STRIPAK than to the *S. cerevisiae* counterpart.

### Yeast two-hybrid analysis of subunit interactions of the STRIPAK complex

We next sought to detect physical interactions between STRIPAK components predicted to be in close proximity. To this end, yeast two-hybrid analysis of each CnSTRIPAK subunit was performed (Fig 2A). Plasmids encoding fusion proteins of the Gal4 DNA-binding domain and CnSTRIPAK subunits Pph22, Tpd3, Far8, Far11, Far9, and Mob3 were generated and transformed into the yeast two-hybrid reporter strain Y2HGold. Similarly, plasmid constructs encoding the Gal4 transcriptional activation domain fused to individual STRIPAK subunits were transformed into reporter strain Y187. A 6-by-6 crossing between Y2HGold and Y187 strains carrying plasmids encoding GBD (Gal4 DNA-binding domain) and GAD (Gal4 activation domain) fusion proteins was conducted and assayed for Gal4-dependent expression of the *ADE2* and *HIS3* reporter genes (Fig S1). For Pph22 and Tpd3, because the GBD fusions alone were able to activate reporter gene expression, only the GAD fusions of these two proteins were analyzed to determine their interactions with GBD-fused STRIPAK components. Positive interactions between Mob3 and Far8 were detected in both configurations, suggesting specific and robust binding between these two proteins (Fig 2A). Positive protein-protein interactions were also detected for Far9-Tpd3, Far11-Pph22, and Far11-Tpd3 (Fig 2A). This yeast two-hybrid analysis provides further support for the predicted CnSTRIPAK model (Figs 1D and 2B).

**Fig 2.**
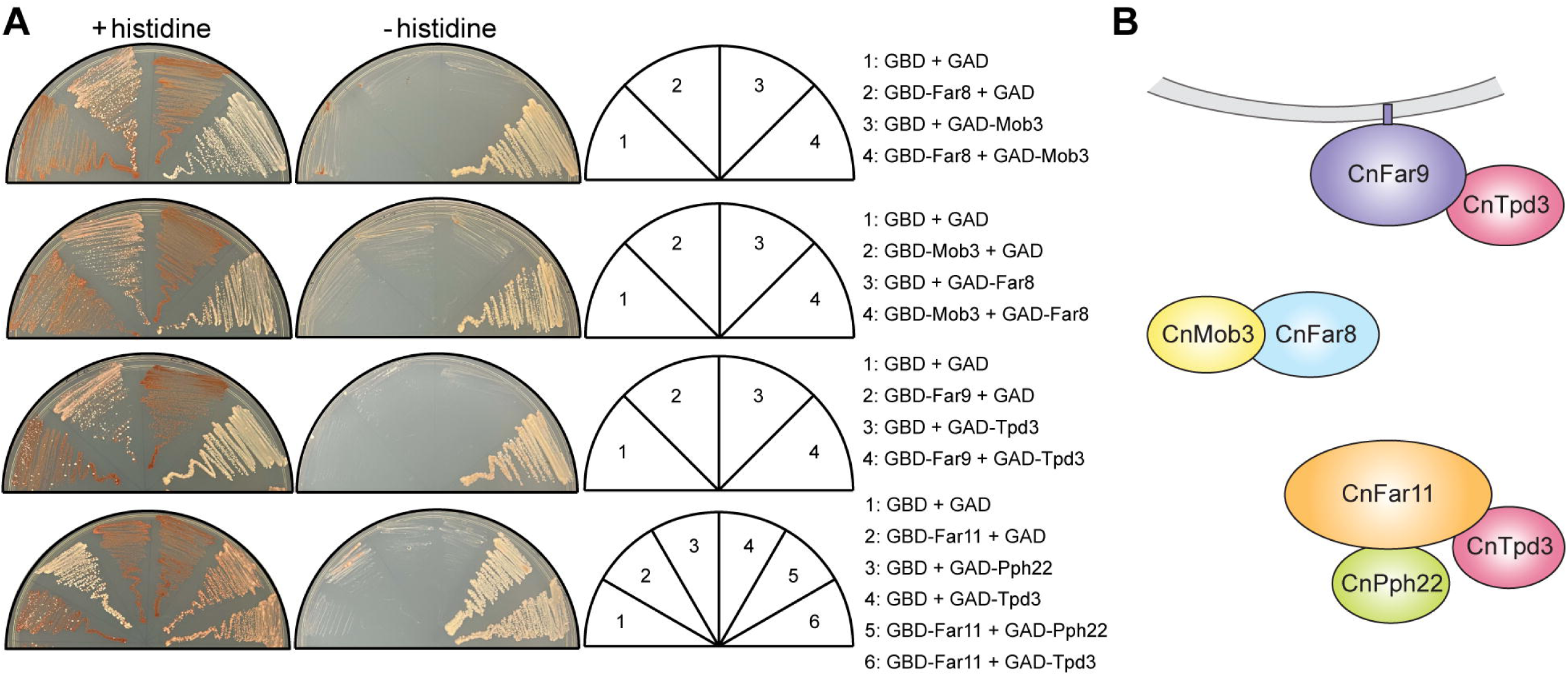
Yeast two-hybrid analysis of STRIPAK complex subunit interactions. (A) *S. cerevisiae* cells carrying plasmids encoding the Gal4 DNA-binding domain (GBD) or the Gal4 transcriptional activation domain (GAD), fused to individual STRIPAK subunits, were crossed. The resulting diploid cells were grown on synthetic dextrose medium with or without histidine. Interaction between bait and prey proteins activates Gal4, driving the expression of *ADE2* and *HIS3*, making cells less red (+histidine plates) and allowing growth on medium deficient in both histidine and adenine (-histidine plates). Results are representative of two independent experiments. (B) Illustration of positive protein-protein interactions detected in the yeast two-hybrid assay.

### Mutations in genes encoding the STRIPAK complex components lead to genome instability

To analyze the function of the *C. neoformans* STRIPAK complex, deletion mutations were generated for genes encoding STRIPAK complex subunits. We first selected the genes encoding the PP2A trimeric enzyme, Pph22, Tpd3, and Far8, for targeted deletion mutagenesis. While we were able to successfully generate *far8*Δ deletion strains, we failed to obtain *pph22*Δ and *tpd3*Δ mutant strains in H99α or KN99**a** haploid wild-type strains, after multiple attempts of both biolistic transformation and CRISPR-Cas9 gene deletion approaches, suggesting that *PPH22* and *TPD3* might be essential for cell viability. In *S. cerevisiae,* the deletion of *TPD3* causes growth defects [47], and deletions of *PPH21* and *PPH22* are synthetically lethal [48]. In *C. neoformans,* a genome-wide functional analysis of phosphatases suggested that *PPH22* is a putative essential gene [41]. Conversely, *FAR8* and other striatin subunit homologs have not been characterized as essential in other species.

We then took a different approach to test the essentiality of *PPH22* and *TPD3,* as well as the other STRIPAK component encoding genes *FAR9*, *FAR11*, and *MOB3*. Specifically, we sought to delete one of the two alleles for each gene in a wild-type diploid strain CnLC6683, which is a fusion product between the congenic strain pair KN99**a** and KN99α, and then isolate haploid deletion mutants by sporulating and dissecting meiotic progeny. Using CRISPR-Cas9 directed mutagenesis, we successfully obtained *PPH22/pph22*Δ and *MOB3/mob3*Δ heterozygous mutants in strain CnLC6683 background. We were unable to isolate heterozygous *TPD3/tpd3*Δ, *FAR9/far9*Δ, or *FAR11/far11*Δ mutant strains for this study; therefore, we concentrated our analyses on *PPH22/pph22*Δ and *MOB3/mob3*Δ. Four *PPH22/pph22*Δ strains were generated from two independent transformations, and three *MOB3/mob3*Δ strains from two independent transformations. PCR genotyping and Illumina whole-genome sequencing confirmed heterozygosity at the *PPH22* and *MOB3* loci. Analysis of read depth revealed multiple segmental and chromosomal aneuploidies throughout the genome (Fig 3A), including increased coverage for entire chromosomes, and both increased and decreased coverage in segments of chromosomes. Whole chromosome 13 trisomy was observed in *PPH22/pph22*Δ*-3, 4* and *MOB3/mob3*Δ-*1, 2, 3* strains, while partial trisomy of this chromosome was seen in *PPH22/pph22*Δ-1. Partial trisomy for additional chromosomes was observed in other cases: chr.4 in *PPH22/pph22*Δ-1; chr. 3 in *PPH22/pph22*Δ-2; chr. 6 in *PPH22/pph22*Δ-3 and *MOB3/mob3*Δ-1; and chr.9 and chr. 10 in *MOB3/mob3*Δ-1. Segmental monosomy was also observed in chr. 2 and chr. 4 in *PPH22/pph22*Δ-4; and chr. 10 in *MOB3/mob3*Δ-1, 3. *PPH22/pph22*Δ-1 also exhibited increased coverage for chr. 1 and chr. 6, although to a lesser extent.

**Fig 3.**
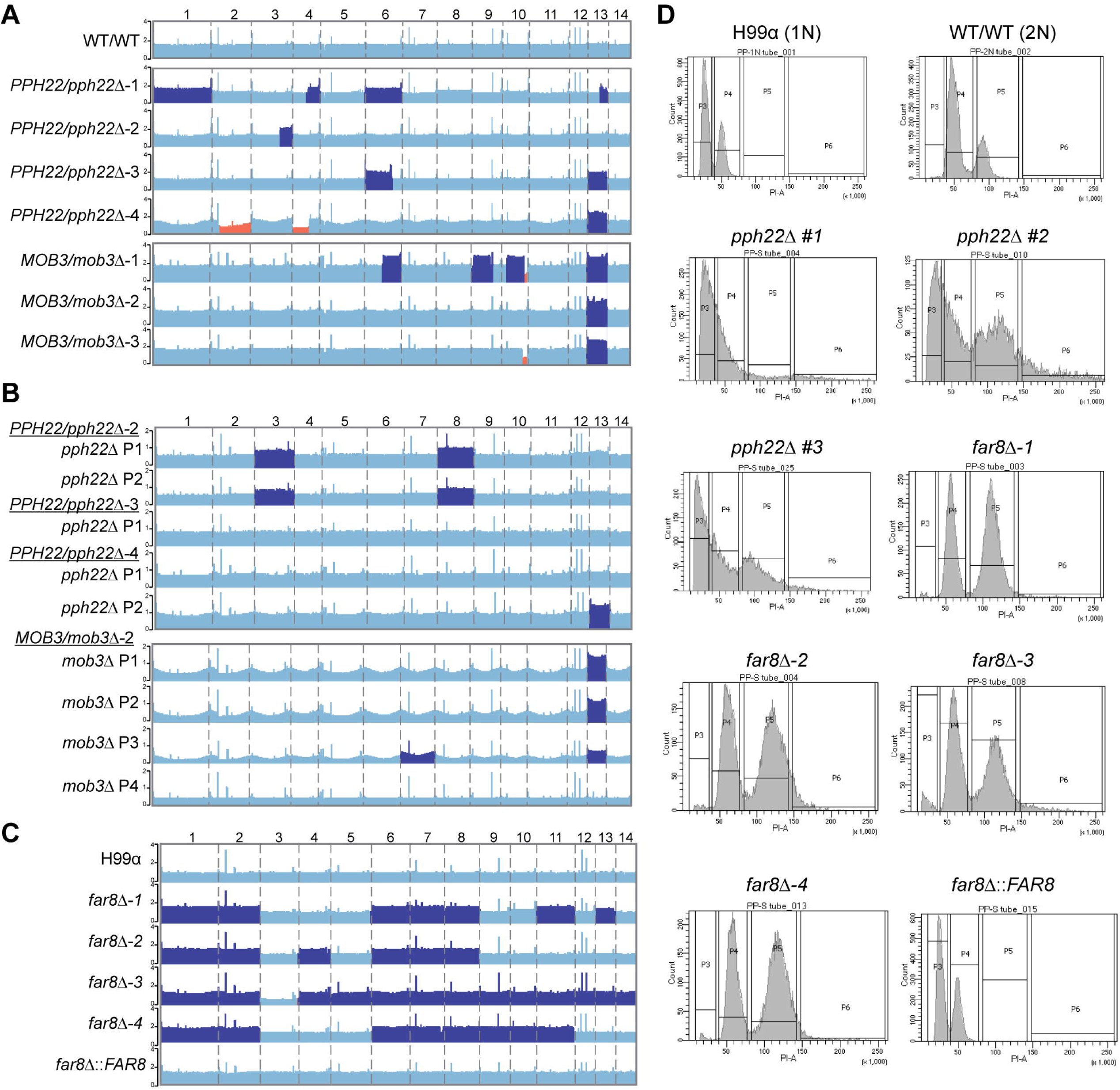
Genome instability in STRIPAK complex mutants. (A) Read depth analyses from whole-genome sequencing of the WT/WT (CnLC6683) parental diploid strain and heterozygous mutant diploid strains. The WT/WT genome is euploid, while *PPH22/pph22*Δ and *MOB3/mob3*Δ strains exhibit multiple instances of segmental and whole chromosome aneuploidy, with dark blue and orange highlighting regions/chromosomes with increased and decreased read depth, respectively. (B) Chromosome maps of *pph22*Δ and *mob3*Δ strains show aneuploidy in some, but not all, haploid progeny. Progeny can also acquire segmental or chromosomal aneuploidies absent in the parental strain. “P” stands for meiotic progeny from the indicated parent. (C) Coverage of *far8*Δ mutants generated in the H99 background reveals duplication of most chromosomes, suggesting a diploid state. Complementation of the deletion allele in the *far8*Δ::*FAR8* strain restores euploidy or results in haploidy. Regions of read coverage in the genome maps in (A-C) are shaded based on *Z* >1.96, *P* <0.05 (dark blue); or *Z* < -1.96, *P* <0.05 (orange). (D) FACS analysis of *pph22*Δ and *far8*Δ strains. Cells were stained with propidium iodide to measure DNA content via flow cytometry. Peaks represent relative DNA content during G_1_/S and G_2_/mitotic phases. H99 and WT/WT (CnLC6683) (KN99**a**/KN99α) served as 1N and 2N controls, respectively. The number sign (#) after a strain name indicates F1 meiotic progeny dissected from a heterozygous mutant diploid parental strain. Graphs are representative of two biological replicates.

We next assessed whether the genetic changes present in the heterozygous mutant diploid populations would be inherited and maintained in meiotic progeny. *PPH22/pph22*Δ and *MOB3/mob3*Δ cells were incubated on MS medium at room temperature for at least three weeks to induce self-filamentation. Basidiospores were dissected, and haploid *pph22*Δ and *mob3*Δ deletion mutant progeny were validated through PCR genotyping (Fig S2). Haploid *pph22*Δ deletion mutants were obtained from the dissection of spores from *PPH22/pph22*Δ*-2, 3*, and *4*, and *mob3*Δ mutants were similarly obtained from *MOB3/mob3*Δ*-2*. Whole-genome sequencing supported the PCR analysis, showing no reads mapping to the *PPH22* or *MOB3* genes. Therefore, *PPH22* and *MOB3* are not essential in *C. neoformans*. Single colonies from *pph22*Δ and *mob3*Δ deletion mutants were passaged at least three times prior to whole-genome sequencing to obtain a pure cell population and reduce the likelihood of sequencing a population of cells with mixed genotypes. Interestingly, whole-genome sequencing analysis showed that whole chromosome aneuploidy was present in some but not all haploid mutant progeny, and that the genomic profile of the progeny often did not mirror the genomic profile of the parent (Figs 3A and 3B). Haploid *pph22*Δ progeny from *PPH22/pph22*Δ-*2* exhibited increased coverage for chr. 3 and chr. 8, while the parental diploid strain was euploid for those chromosomes. One other haploid strain showed aneuploidy that was not present in its diploid parent: P3 dissected from *MOB3/mob3*Δ*-2* was disomic for chr. 7 (Figs 3A and 3B). The most common aneuploidy present in the heterozygous diploid progenitor strains was associated with chr. 13 (Fig 3A), which was found in P2 from *PPH22/pph22*Δ*-4* and P1-P3 from *MOB3/mob3*Δ*-2*. It has been previously reported that chromosome 13 disomy is common among clinical isolates, but it remains unclear whether unique elements on this chromosome are specifically linked to virulence [49]. No ploidy changes were detected in any of the haploid and diploid control strains from each experiment. These results suggest that the *PPH22* and *MOB3* genes play roles in maintaining genome stability.

Genetic analysis of *FAR8* revealed a related role in genome stability. The *far8*Δ mutants were successfully isolated via biolistic transformation in the congenic H99α and YL99**a** haploid strains. Deletion of *FAR8* was confirmed by PCR genotyping and Illumina whole-genome sequencing (Fig S3). Read-depth analyses indicated that the four independent *far8*Δ mutants in H99α had rampant aneuploidy throughout the genome, suggesting that they were diploid instead of haploid (Fig 3C). Similar results were obtained from whole-genome sequencing of YL99**a** *far8*Δ strains (Fig S4). Among seven independently obtained *far8*Δ mutant strains, all were aneuploid for at least six out of 14 chromosomes, and the only chromosome that was never found to be duplicated was chr. 3. The parental H99α and YL99**a** isolates were haploid, confirming the aneuploidy occurred after deletion of *FAR8.* Complementation of the *far8*Δ mutation in the H99α background strain *far8*Δ-*1* yielded a euploid population, suggesting reintroduction of *FAR8* prevents formation of aneuploidy.

Fluorescence-activated cell sorting (FACS) analysis was conducted to determine the DNA content in the STRIPAK mutants (Figs 3D and S4). A wild-type haploid shows a major 1C peak and a minor 2C peak, while a wild-type diploid has a major 2C peak and a minor 4C peak. The *pph22*Δ mutants displayed either one single merged, widened peak, or additional widened peaks at 4C or larger, indicating that there is a heterogeneous population of cells with different DNA content with modest to severe aneuploidy. Another possibility is that *pph22*Δ mutants form clusters or have defects in cytokinesis, as described later in Fig 4C. FACS analysis of *far8*Δ mutants showed two major peaks at 2C and 4C, suggesting that these strains are largely diploid. FACS data from the *far8*Δ::*FAR8* complemented strain had a similar peak profile to the 1N control strain H99α, indicating that it is haploid. The *mob3*Δ mutants were also analyzed by FACS and exhibited only two peaks at 1C and 2C, indicating they are haploid (Fig S4).

**Fig 4.**
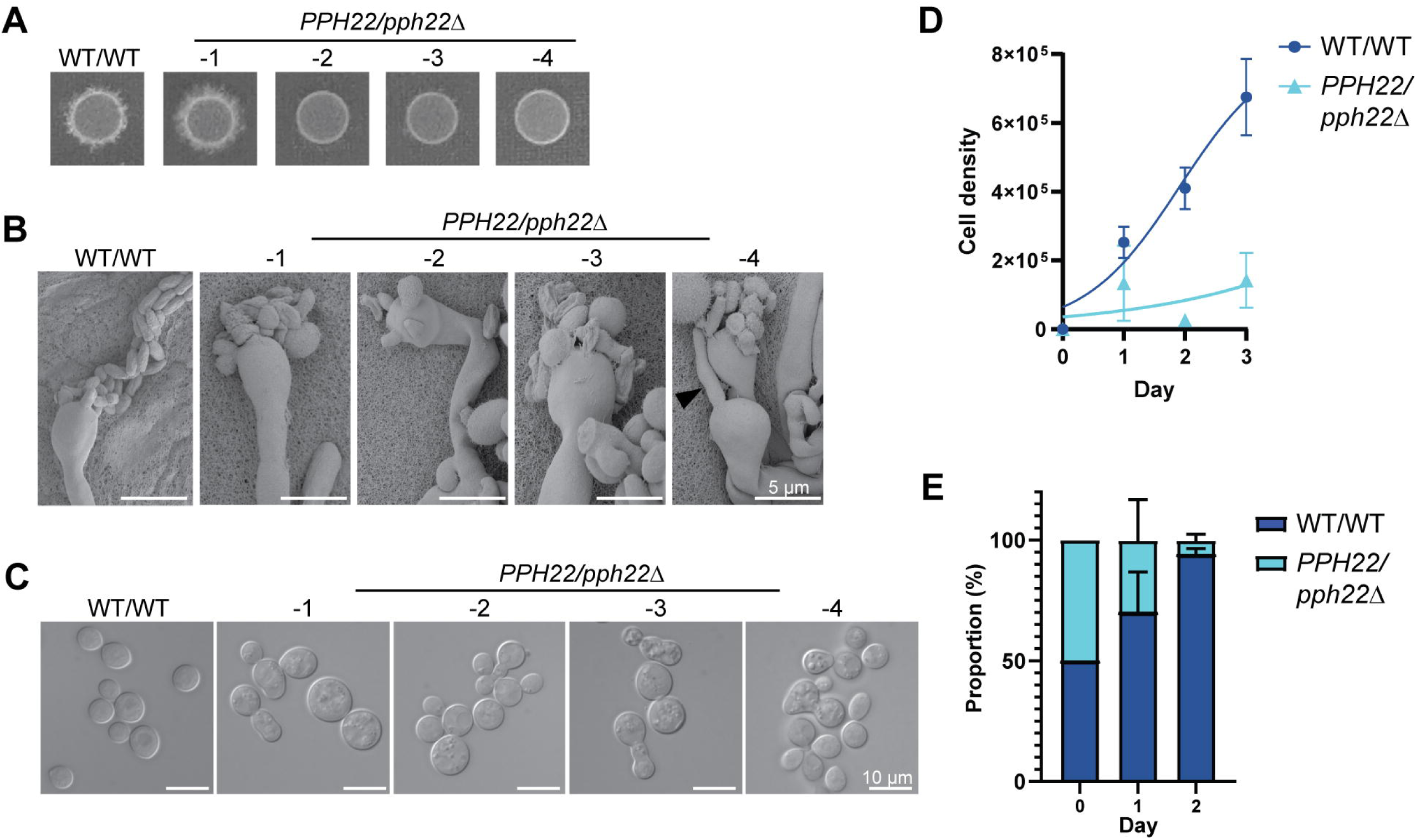
Deletion of *PPH22* in a diploid background leads to haploinsufficiency. (A) Self-filamentation of WT/WT (CnLC6683) and *PPH22/pph22*Δ diploid strains on MS medium. Strains were grown on MS and incubated at room temperature (24°C) in the dark for 3 to 4 weeks before images were taken. Each spot represents an independent experiment. A representative image from one out of four independent experiments for each strain is shown. (B) Scanning electron microscopy (SEM) analysis of basidia and basidiospores from WT/WT and *PPH22/pph22*Δ strains. Samples were prepared following incubation on MS media for four weeks [shown in (A)]. The black arrowhead in image *PPH22/pph22*Δ-4 shows a hyphal filament being produced from a basidia head. Scale bars represent 5 μm. (C) Differential interference contrast (DIC) microscopy images of WT/WT and *PPH22/pph22*Δ cells grown to mid-logarithmic phase in YPD liquid media at 30°C. *PPH22/pph22*Δ cells showed elongated cells, increased cell size, incomplete budding, and abnormal cell clusters. Cultures were grown in biological replicates for analysis. Scale bars represent 10 μm. (D) Growth rates of WT/WT and *PPH22/pph22*Δ strains grown in YPD liquid culture at 30°C. Cell density was quantified as cells/mL, and logarithmic growth was modeled using nonlinear regression. (E) Competition assay of WT/WT versus *PPH22/pph22*Δ. The growth of strains in the coculture is expressed as a percentage of the total number of cells at the indicated time point. The data correspond to the mean of four biological replicates ± standard deviation.

Taken together, whole-genome sequencing and FACS analysis showed that *pph22*Δ, *far8*Δ, and *mob3*Δ mutations cause genome instability, resulting in aneuploidy caused by whole chromosome and segmental duplication/deletion. Our data also suggest that the deletion of *FAR8* might have led to genome endoreplication, which was followed by rampant chromosomal losses resulting in aneuploidy, suggesting a role in cell cycle control. These results demonstrate that the *C. neoformans* STRIPAK complex is important for maintaining genome stability.

### *PPH22* is a haploinsufficient gene in diploid *C. neoformans*

During self-filamentation, we observed that *PPH22/pph22*Δ strains showed delayed production of hyphae, basidia, and basidiospores compared to the wild-type control. To analyze this further, equal amounts of WT/WT (CnLC6683) and *PPH22/pph22*Δ diploid cells were plated on MS media and incubated at room temperature. After 1 to 2 weeks, the wild-type diploid strain produced abundant hyphae and basidia, and basidia with spore chains were also visible. However, no hyphae were seen in the *PPH22/pph22*Δ strains at that time. After 3 to 4 weeks of incubation, the patch of *PPH22/pph22*Δ-*1* had robust filamentation (Fig 4A), and hyphae, basidia, and spore chains were observed at high magnification. However, patches of *PPH22/pph22*Δ-*2, 3, 4* cells exhibited only minimal hyphae, basidia, and spores. Hyphae, basidia, and basidiospores of the self-filamenting diploid strains were observed by scanning electron microscopy (SEM) (Fig 4B). The basidia in *PPH22/pph22*Δ strains exhibited abnormal and fewer spore formation, with spores produced in irregular clusters instead of chains. The morphology of the spores was also atypical, with significant variation in shape and size. There were also instances of ectopic hyphal growth, as seen in *PPH22/pph22*Δ*-4*, where basidia appeared to produce a new hyphal filament instead of spores (Fig 4B).

The defects in sexual development observed in *PPH22/pph22*Δ strains led us to hypothesize that *PPH22* might be haploinsufficient. When WT/WT and *PPH22/pph22*Δ strains were grown to mid-logarithmic phase in YPD liquid media at 30°C and cell morphology was observed using differential interference contrast (DIC) microscopy, *PPH22/pph22*Δ cells were typically larger and formed abnormal clusters compared to wild-type diploid cells (Fig 4C). Some *PPH22/pph22*Δ cells were elongated and showed incomplete budding. The *PPH22/pph22*Δ-*1-4* strains also exhibited severe growth defects compared to the WT/WT strain in liquid culture (Fig 4D). In competition assays to compare the fitness of the WT/WT and *PPH22/pph22*Δ strains, the *pph22*Δ deletion allele in the heterozygous diploid strain conferred a significant competitive disadvantage compared to the wild type (Fig 4E). To determine if this disadvantage was due to reduced viability of *PPH22/pph22*Δ strains, *PPH22/pph22*Δ and WT/WT cells that had been grown in YPD overnight cultures for the competition assays were serially diluted and plated onto YPD medium at 30°C (Fig S5). There was no observable difference in growth between the wild-type and heterozygous mutant diploid strains, indicating cells were similarly viable at the time of competition. The significant reduction in the *PPH22/pph22*Δ population at the end of the competition assay likely results from its inherent growth defects, leading to reduced fitness compared to the wild type. Thus, the loss of one *PPH22* allele in a diploid leads to defects in sexual development, cell morphology, and competitive growth, indicating *PPH22* is a haploinsufficient gene. This may be attributable to loss of one copy of *PPH22*, aneuploidy arising due to reduction in *PPH22* level, or both.

### Haploid *pph22*Δ mutants exhibit severe growth defects and frequently accumulate suppressor mutations

To further characterize the functions of *PPH22*, haploid *pph22*Δ mutants were obtained for phenotypic analysis. Self-filamenting *PPH22/pph22*Δ-*1-4* strains produced sufficient basidiospores for dissection after 4 to 6 weeks of incubation on MS plates. *pph22*Δ mutant colonies were much smaller than wild-type colonies on the dissection plate (Fig 5A). The severe growth defects of *pph22*Δ mutants likely explain why *PPH22* was initially thought to be essential. *pph22*Δ mutants formed tan colonies on YPD plates incubated at 30°C. After prolonged incubation, faster-growing white colonies appeared, possibly as a result of suppressor mutations (referred to as *pph22*Δ *suppressor* or *pph22*Δ *sup* mutants) (Fig 5B). While *pph22*Δ strains grew poorly compared to wild type at 24°C and 30°C, produced tan colonies, and could not grow at 33°C, the *pph22*Δ *sup* strains exhibited near wild-type growth at 24°C, 30°C, and 33°C, and produced whiter colonies (Fig 5C).

**Fig 5.**
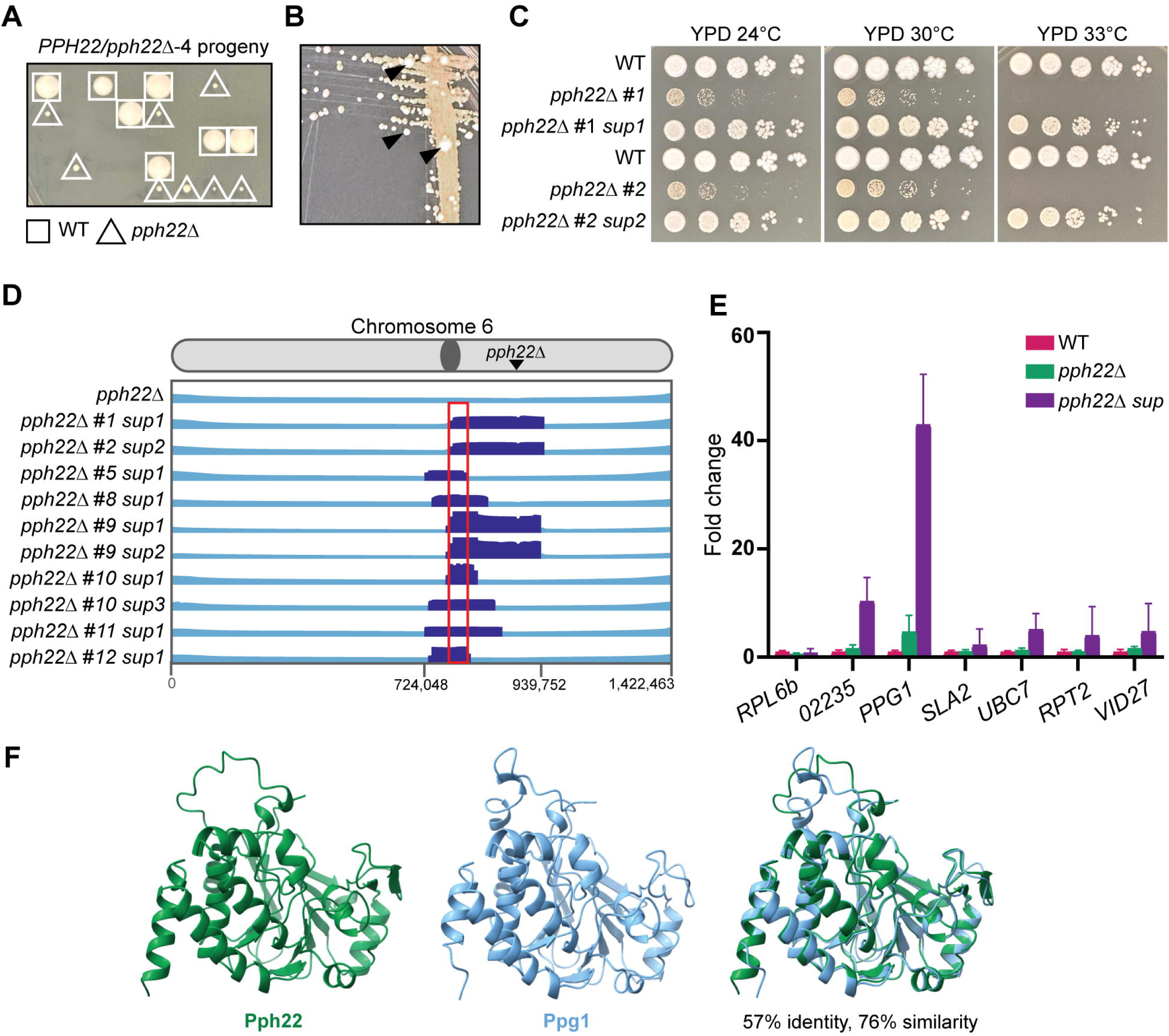
*pph22*Δ mutants frequently accumulate suppressor mutations. (A) Representative image showing the growth of spores dissected from the *PPH22/pph22*Δ*-4* strain on YPD medium. Wild-type spores are marked with a square, and *pph22*Δ deletion mutant spores with a triangle. The image was taken after five days of incubation at 30°C. (B) *pph22*Δ strains exhibit very slow growth and form tan-colored colonies. After prolonged incubation, large white colonies appear (indicated by black arrowheads), resulting from suppressor mutations. The image shows a YPD plate incubated at 30°C for one week. (C) Wild-type (KN99), isogenic *pph22*Δ, and *pph22*Δ *suppressor* (*pph22*Δ *sup*) strains were serially diluted and spotted onto YPD medium at 24°C, 30°C, and 33°C. Images were taken after 5 days of incubation. (D) *pph22*Δ *sup* strains show shared aneuploidy on chromosome 6. The diagram displays read coverage from whole-genome sequencing, with chromosomal coordinates at the bottom. The region of increased coverage shared by all 10 *pph22*Δ *sup* strains is indicated by a red box, which spans 7 genes. The read depth in this region is at least 3 times higher than the average depth of the rest of chromosome 6. (E) Fold change in expression of the 7 genes with a complete ORF included in the ∼15 kb boxed region from (D), determined from RNA-seq. Fold changes in gene expression in *pph22*Δ (PP53, PP55, PP56) and *pph22*Δ *sup* (PP80, PP83, PP84) samples relative to wild type (WT, KN99**a**), calculated by normalizing each gene’s FKPM to the housekeeping gene *ACT1.* Error bars represent the standard deviations calculated from biological triplicates. The gene ID numbers, from left to right in the graph, are CNAG_02234, CNAG_02235, CNAG_02236, CNAG_02237, CNAG_02238, CNAG_02239, and CNAG_02240. (F) AlphaFold structure prediction of Pph22 and Ppg1 and a superimposition of the two proteins. Root mean square deviation (RMSD) values of superimposed structures are 0.507 Å (between 285 pruned atom pairs) and 1.644 Å (across all 302 atom pairs).

Ten *pph22*Δ *suppressor* strains were subjected to whole-genome sequencing to identify the causative suppressor mutation(s). Variant calling failed to identify any significant single-nucleotide polymorphisms (SNPs) or insertion/deletion mutations. However, all 10 *pph22*Δ *suppressor* mutant strains exhibited segmental aneuploidy on chr. 6 in a shared overlapping region of ∼200 kbp (Fig 5D) and were euploid for the remainder of their genomes. Read depth in this region ranged from 2X to 11X coverage compared to the mean coverage of the remainder of the chromosome. The *PPH22* gene lies in this region, but as expected, no reads were found mapping to the *PPH22* locus. All 10 isolates shared an approximately 15 kbp region adjacent to the centromere, encompassing 7 genes (Figs 5D and 5E). One notable candidate for the *pph22*Δ *suppressor* gene is *PPG1,* which encodes a serine/threonine type 2A-like protein phosphatase catalytic subunit involved in the cell wall integrity pathway [50]. Studies from *S. cerevisiae* have shown that Ppg1, and not Pph22, interacts with Far11 to regulate assembly of the Far complex [51, 52], which may have pleiotropic roles beyond those of the STRIPAK complex. Expression data collected from RNA sequencing of wild type (KN99**a**), *pph22*Δ, and *pph22*Δ *suppressor* strains was used to determine the FKPM (fragments per kilobase per million mapped fragments) of these 7 candidate genes, along with the housekeeping gene *ACT1 (*Fig 5E). Normalized expression data is plotted in the graph as fold change in expression compared to wild type. In *pph22*Δ *suppressors, PPG1* was overexpressed 43-fold compared to wild type. *SLA2, UBC7, RPT2,* and *VID27* showed modest increases in expression (2- to 5-fold) and the hypothetical protein CNAG_02235 increased in expression by 10-fold over wild type. The detected increase in expression of CNAG_02235 may be due to sequence overlap with 4 exons of the *PPG1* gene. Interestingly, *PPG1* was also moderately overexpressed in *pph22*Δ mutants (4.7-fold increase), suggesting that upregulation of *PPG1* may be a compensatory mechanism to mitigate the loss of *PPH22.* This finding suggests that Ppg1 may share functional similarities with Pph22 and could also interact to form a complex with PP2A. Indeed, while the protein sequence alignment showed that Ppg1 shares 57% identity and 76% similarity with Pph22, their predicted 3D structures appeared to be virtually indistinguishable (Fig 5F). Additionally, AlphaFold predicted that Ppg1 could replace Pph22 as the catalytic subunit within the PP2A heterotrimeric complex with Tpd3 and Far8 (Fig S6). Taken together, our data suggest that significantly increased expression of *PPG1* may partially compensate for the loss of *PPH22*, supporting the hypothesis that Ppg1 may functionally substitute for Pph22.

### STRIPAK is important for mating and sexual development in *C. neoformans*

To investigate roles for the *C. neoformans* STRIPAK complex in sexual development, we analyzed the effects of mutations in STRIPAK components on mating and the sexual cycle using *PPH22/pph22*Δ and *MOB3/mob3*Δ heterozygous diploid strains compared to WT/WT (Fig 6A). As observed previously, *PPH22/pph22*Δ exhibited severe self-filamentation defects. Similarly, *MOB3/mob3*Δ displayed defects in sexual development, showing a smaller area of self-filamentation and shortened hyphae compared to the wild type. Although *MOB3/mob3*Δ was able to produce basidia and basidiospores, this occurred only near the edge of the cell patch, indicating that *MOB3* is also important for sexual development in a self-filamenting diploid.

**Fig 6.**
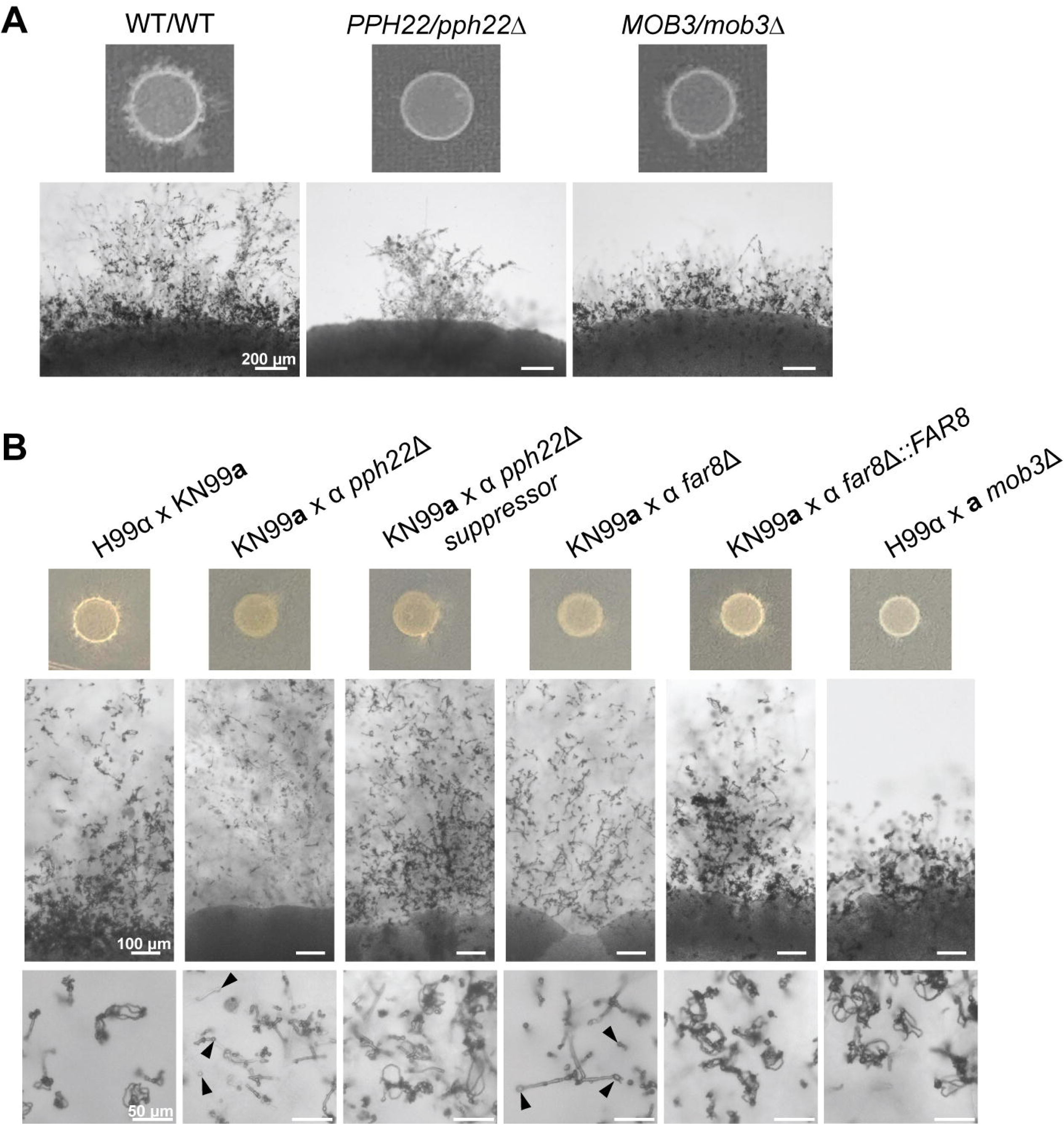
Deletion mutations in *PPH22, FAR8,* and *MOB3* lead to defects in sexual differentiation. (A) *PPH22/pph22*Δ and *MOB3/mob3*Δ strains exhibit defects in self-filamentation compared to WT/WT (CnLC6683). Images were taken after four weeks of incubation on MS plates at room temperature. Scale bars represent 200 μm. (B) Mating efficiency of STRIPAK complex mutants. *pph22*Δ, *pph22*Δ *sup, far8*Δ, *mob3*Δ, and *far8*Δ::*FAR8* cells were co-cultured with isogenic wild-type cells of the opposite mating type (H99α or KN99**a**) on MS plates. An H99α x KN99**a** cross served as a control. Black arrows indicate basidia that did not produce spores in KN99**a** x α *pph22*Δ and KN99**a** x α *far8*Δ crosses. Scale bars represent 100 μm and 50 μm in the middle and bottom panels, respectively.

We next examined the role of the STRIPAK complex during bisexual mating. Mutant cells (*pph22*Δ, *pph22*Δ *suppressor, mob3*Δ, and *far8*Δ) and the complemented *far8*Δ::*FAR8* strain were mixed with wild-type cells of the opposite mating type, spotted onto MS media, and incubated at room temperature (Fig 6B). The H99α x KN99**a** cross served as a wild-type control. Initial crosses involving *pph22*Δ and *far8*Δ strains did not show any signs of mating, even after 8 weeks. This issue could be due to the growth defects of these strains on MS media, leading to them being outcompeted by the wild-type partner. Additional crosses were conducted utilizing two strategies to promote mating: (1) mixing mutant and wild-type cells in a 10:1 ratio, and (2) growing mutant cells alone for two to three days on MS media before plating wild-type cells on top. These strategies enabled mating in *pph22*Δ x WT and *far8*Δ x WT crosses. *pph22*Δ, *pph22*Δ *suppressor,* and *far8*Δ crosses with WT could form large branches of hyphae extending from the mating patch, similar to the wild-type control. The *pph22*Δ x WT cross could produce hyphae, but mostly bald basidia heads with almost no spore formation. This phenotype was partially rescued in the *pph22*Δ *suppressor* mutants, which produced basidia with basidiospores after mating, albeit to a lesser extent than the wild-type control.

Mating of *far8*Δ x WT strains is an unusual case, as the *far8*Δ mutant contains a largely diploid genome. This results in a diploid-by-haploid cross followed by triploid meiosis. Similar to *pph22*Δ x WT crosses, we observed mostly bald basidia heads with no spores in *far8*Δ x WT crosses, though some basidia with spore chains were found at low frequency. Crossing WT with the *far8*Δ::*FAR8* complemented strain successfully restored mating and sporulation efficiency to the wild-type level. The *mob3*Δ and wild-type cells formed stunted hyphae and a smaller area of hyphal growth, resembling the self-filamentation of the *MOB3/mob3*Δ heterozygous diploid strain, but produced abundant basidia and basidiospore chains. We did not observe mating during bilateral crosses between STRIPAK component deletion mutants (Fig S7). This suggests that there are additive interactions between the defects caused by each individual deletion of either *PPH22*, *FAR8*, or *MOB3*. In summary, the STRIPAK complex plays a critical role in various aspects of sexual development in *C. neoformans*.

### Phenotypes of *pph22*Δ and *far8*Δ mutants in response to nutrients, temperature, and stress

Next, the role of STRIPAK in vegetative growth and stress response was analyzed. *pph22*Δ and *far8*Δ mutant strains, along with *pph22*Δ *suppressors*, *far8*Δ::*FAR8,* and isogenic wild-type control strains were serially diluted and spotted onto media under different nutrient, temperature, and stress conditions (Figs 7A and 7B). *pph22*Δ mutants exhibited severe growth defects on YPD, which was exacerbated by supplementation with 1 M sorbitol (Fig 7A). This finding could explain why we were unable to generate a haploid *pph22*Δ deletion mutant via biolistic transformation, during which sorbitol medium was used to provide osmotic support for cells to recover from puncture by DNA-coated gold particles [53].

**Fig 7.**
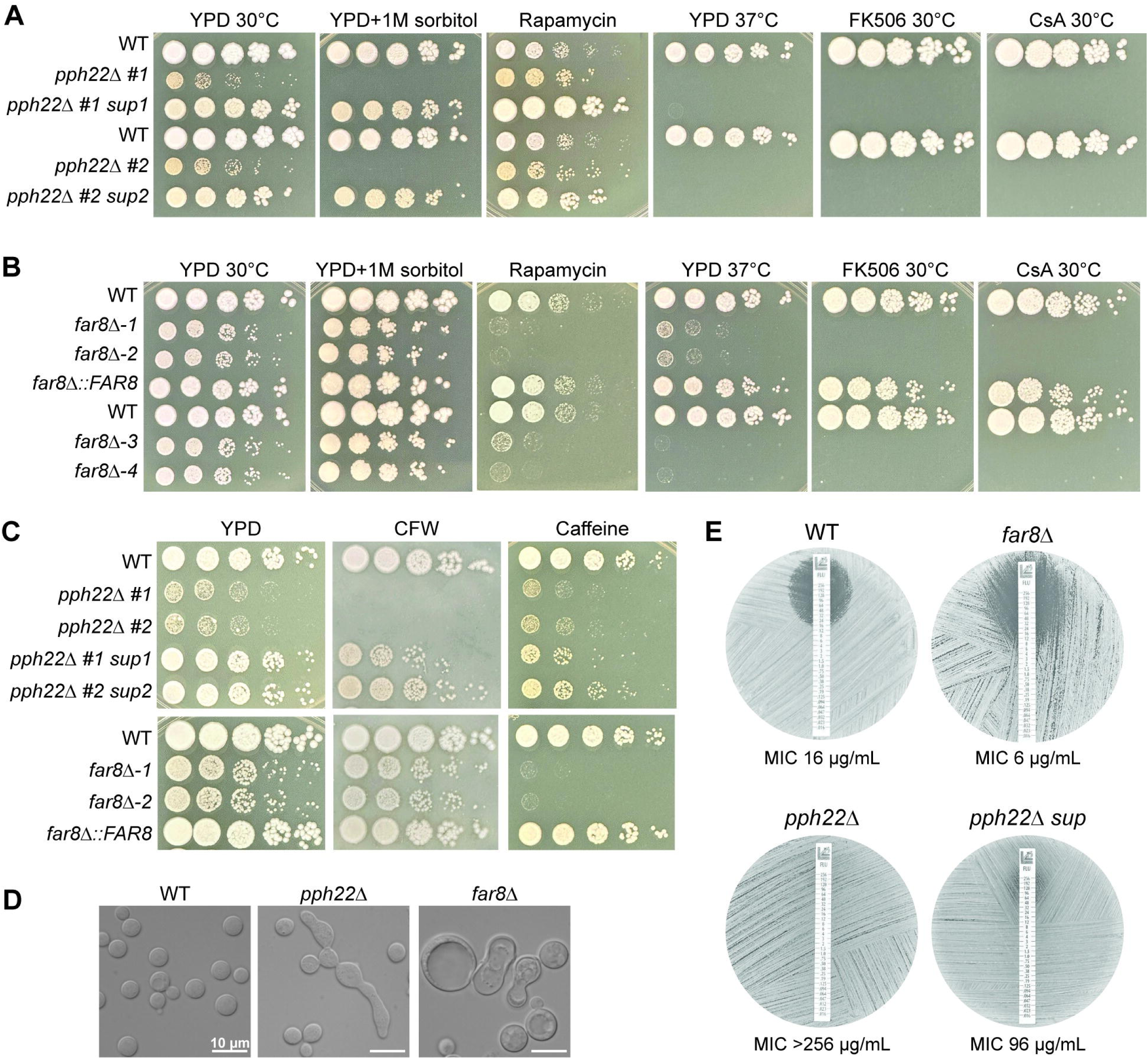
Phenotypic analyses of *pph22*Δ and *far8*Δ mutants. (A) WT (KN99**a**) and isogenic *pph22*Δ and *pph22*Δ *sup* strains were serially diluted and plated on YPD, YPD+1 M sorbitol, YPD+100 ng/mL rapamycin, YPD+1 μg/mL FK506, and YPD+100 μg/mL cyclosporine A (CsA) at 30°C, as well as YPD at 37°C. Images were taken between 2 and 5 days of incubation. This set is from the same experiment as Fig 5C. (B) WT (H99) and isogenic *far8*Δ-*1, far8*Δ*-2,* and *far8*Δ::*FAR8*, along with WT (YL99**a**) and isogenic *far8*Δ-*3* and *far8*Δ*-4* strains, were serially diluted and plated on the indicated media and temperature conditions. (C) WT, *pph22*Δ, *pph22*Δ *sup, far8*Δ, and *far8*Δ::*FAR8* cells were serially diluted and spotted onto YPD supplemented with 3 mg/mL calcofluor white (CFW) or 0.5 mg/mL caffeine. Plates were incubated at 30°C and images were taken after 3 days. (D) DIC microscopy images depicting cell morphology of WT (H99), *pph22*Δ, and *far8*Δ strains grown to mid-logarithmic phase in synthetic complete (SC) media at 30°C. Scale bars represent 10 μm. (E) Fluconazole Etest to analyze drug susceptibility in WT (H99), *far8*Δ, *pph22*Δ, and *pph22*Δ *sup* strains. Cells were grown in an overnight culture in YPD to saturation, and then spread onto YPD plates before adding FLC Etest strips. Plates were incubated at 30°C and images were taken after 48 hours. Drug sensitivities are representative of two biological replicates.

The *pph22*Δ mutants were then tested on YPD with rapamycin, which inhibits TORC1 and mimics nutrient starvation (Fig 7A). Interestingly, *pph22*Δ grew slightly better than the wild type, while the *pph22*Δ *#1 sup1* and *pph22*Δ *#2 sup2* strains exhibited robust growth on this medium. This suggests that *pph22*Δ leads to rapamycin tolerance, and that the suppressor mutations of *pph22*Δ, potentially via overexpression of the phosphatase *PPG1*, can confer further resistance to rapamycin. On YPD medium at 37°C, neither *pph22*Δ nor *pph22*Δ *sup* strains grew, indicating *PPH22* is required for high-temperature growth, and the suppressor mutation does not rescue the defect. The *pph22*Δ and *pph22*Δ *sup* cells also failed to grow in the presence of the immunosuppressive drugs FK506 or cyclosporine A at 30°C, both of which inhibit the activity of the PP2B phosphatase, calcineurin, illustrating synthetic lethality due to the loss of two phosphatases.

A similar role in nutrient and stress response was investigated for *FAR8.* In serial dilution assays, *far8*Δ cells grew more slowly on YPD at 30°C and formed smaller colonies compared to the wild type (Fig 7B). This growth defect was partially rescued by the supplementation of sorbitol. Contrary to our observations of *pph22*Δ cells grown in the presence of rapamycin, *far8*Δ strains were severely growth impaired on this medium compared to wild type, suggesting that these strains have different responses to TORC1 inhibition. *far8*Δ*-1* and -2 strains exhibited severe growth defects on YPD at 37°C, while *far8*Δ*-3* and -*4* strains (constructed in the YL99**a** background) exhibited almost no growth. Similar to *pph22*Δ mutants, *far8*Δ strains did not grow in the presence of FK506 or cyclosporine A at 30°C, suggesting a synthetic lethal interaction between the deletion of *FAR8* and the inhibition of calcineurin by FK506 or cyclosporine A.

These findings suggest that PP2A is important for cell growth under various stress conditions and that *PPH22* in particular may have a role in response to osmotic stress or in maintaining cell wall integrity. The mutants were tested for sensitivity to cell wall-disrupting agents (calcofluor white and caffeine) (Fig 7C). *pph22*Δ mutants were unable to grow in the presence of calcofluor white, which was partially rescued in the *pph22*Δ *#1 sup1* and *pph22*Δ *#2 sup2* strains. In the presence of calcofluor white, *far8*Δ mutants exhibited a subtle growth defect compared to WT, similar to YPD alone. On media supplemented with caffeine, both wild-type and *pph22*Δ cells grew slightly poorer and formed smaller colonies than on YPD, while the *pph22*Δ *sup* strains exhibited strong caffeine sensitivity. *far8*Δ cells showed even higher caffeine sensitivity and exhibited almost no growth. Therefore, *pph22*Δ and *far8*Δ mutants display opposite phenotypes in response to calcofluor white and caffeine. DIC microscopy images of *pph22*Δ and *far8*Δ cells grown in nutrient-limiting synthetic complete (SC) medium also revealed changes in cell morphology (Fig 7D). Specifically, while cells of the wild-type strain were homogenously round and typical in size, both *pph22*Δ and *far8*Δ mutants produced heterogenous cell populations, which were mixtures of normal-looking cells with elongated cells (*pph22*Δ) or considerably enlarged cells (*far8*Δ), suggesting both *PPH22* and *FAR8* are important for faithful mitotic cytokinesis.

The above growth analyses of the mutant strains indicated severe growth impairment when exposed to conditions affecting cell wall integrity. We hypothesized that *pph22*Δ and *far8*Δ mutants might exhibit hypersensitivity to the clinically relevant antifungal drug fluconazole, which targets the ergosterol biosynthesis pathway and weakens the fungal cell membrane. We investigated this with Etest strips (Fig 7E). As predicted, the *far8*Δ strain showed a larger zone of inhibition and increased susceptibility to fluconazole compared to the wild-type control. Surprisingly, *pph22*Δ mutants were completely resistant to fluconazole at the full range of concentrations on the Etest strip, suggesting an MIC value greater than 256 μg/mL. A *pph22*Δ *sup* mutant was also highly resistant to fluconazole. Similar results were obtained for independent *pph22*Δ, *pph22*Δ *sup,* and *far8*Δ mutant strains (Fig S8).

### *PPH22* and *FAR8* are crucial for melaninization, capsule production, and virulence in *C. neoformans*

The role of PP2A-STRIPAK components, *PPH22* and *FAR8*, was next addressed in the production of two important virulence factors in *C. neoformans*: melanin and the polysaccharide capsule. On Niger seed agar, *pph22*Δ mutants showed severe growth impairment, complicating the assessment of melaninization (Fig 8A). The *pph22*Δ *sup* strains also exhibited growth defects but produced some melanin pigment. The *far8*Δ strains failed to produce melanin on this media and remained white, similar to the *lac1*Δ mutant control. Copper availability is known to regulate melaninization [54], prompting us to test whether the melanin defect in *far8*Δ strains is copper-dependent. Melanin production of the mutants was assessed on L-DOPA medium, supplemented with either copper sulfate or the copper chelator, bathocuproine disulfonate (BCS) (Fig 8B). For controls, we used strains H99α (produces melanin), H99α *cbi1*Δ *ctr4*Δ (produces melanin only when copper is present), and H99α *lac1*Δ (melanin production defective). Under the copper-deficient condition (BCS), *pph22*Δ produced minimal melanin, which was partially restored in the suppressor mutant, resembling observations on Niger seed medium. The *far8*Δ strains did not produce melanin in the presence of BCS. When copper sulfate was added, the *pph22*Δ strains did not show increased melaninization, but the *pph22*Δ *#1 sup1* strain did show a copper-dependent increase in pigment. The *far8*Δ strains were only able to produce melanin when copper was added to the media, suggesting that their inability to make melanin is partially linked to defects in maintaining copper homeostasis.

**Fig 8.**
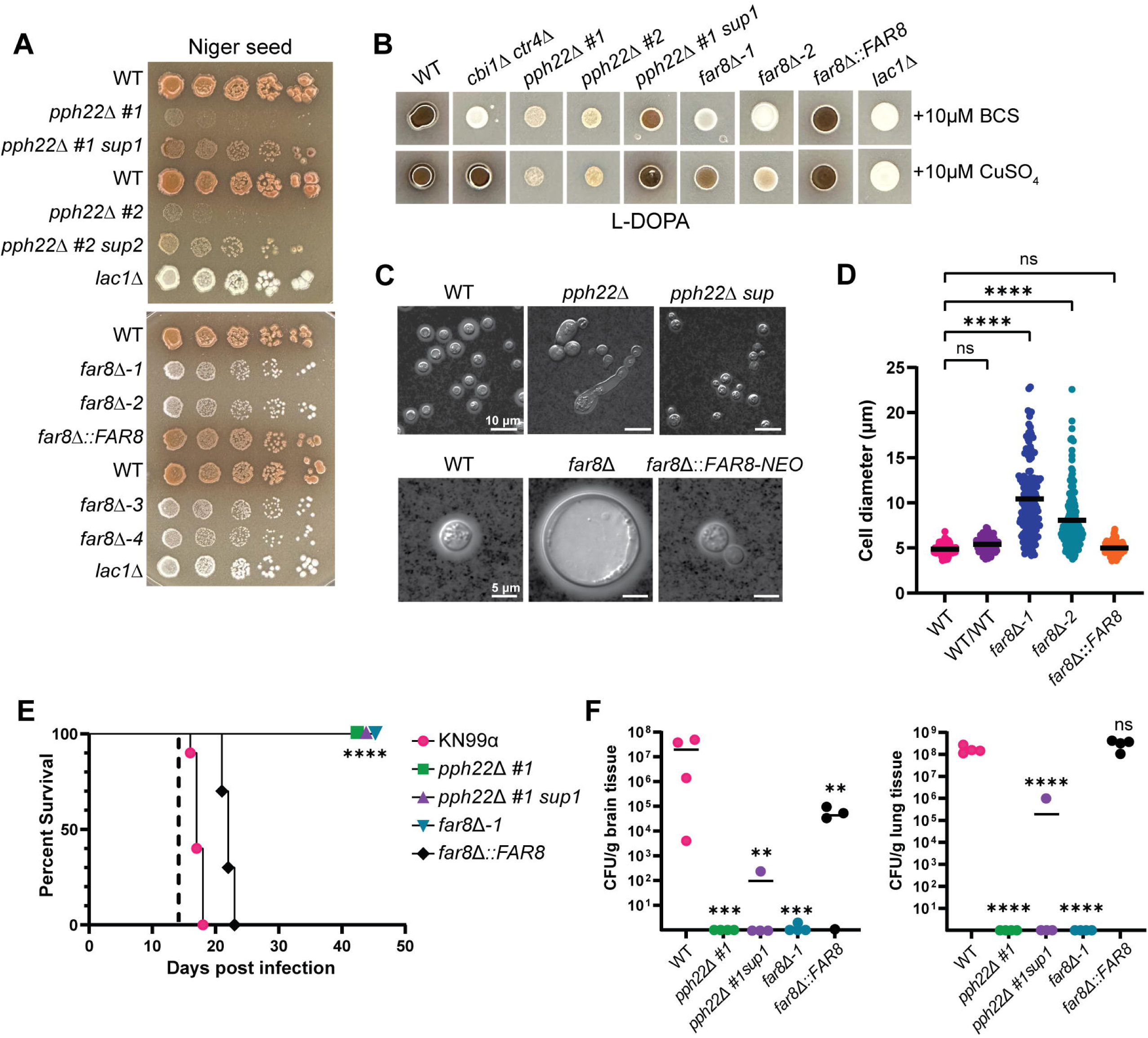
Role of *PPH22* and *FAR8* in production of virulence factors and pathogenicity. (A) WT (KN99**a**, H99α, or YL99**a**) and isogenic *pph22*Δ, *pph22*Δ *sup, far8*Δ, and *far8*Δ::*FAR8* strains were serially diluted and plated onto Niger seed agar medium to induce melanin production. The *lac1*Δ mutant was included as a negative control. Plates were incubated at 30°C for 7 days in the top panel and 4 days in the bottom panel. (B) Analysis of copper-dependent melanin production in *pph22*Δ and *far8*Δ mutants. The indicated strains were grown in a YPD overnight culture to saturation and spotted onto L-DOPA plates supplemented with either 10 μM of a copper chelator, bathocuproine disulfonate (BCS), or 10 μM copper(II) sulfate (Cu_2_SO_4_). The *cbi1*Δ *ctr4*Δ double mutant, which can only produce melanin in the copper-supplemented condition, served as a positive control. Plates were incubated at 30°C for 2 days. (C) Analysis of capsule formation by India ink staining of WT (KN99**a** or H99α), *pph22*Δ, *pph22*Δ *sup, far8*Δ, and *far8*Δ::*FAR8* cells. Strains were grown for 3 days at 30°C in RPMI media to induce capsule formation. Cells were harvested, resuspended in PBS, and stained with India ink. The scale bar for the top panel of DIC images is 10 µm, and the scale bar for the bottom panel of images is 5 µm. (D) Cell size analysis of WT (H99), WT/WT (CnLC6683) (KN99**a**/KN99α), *far8*Δ, and *far8*Δ::*FAR8* complemented strains. Cells analyzed were from the same experiment as (C). Images were analyzed with ImageJ/Fiji. Data are presented as a scatter dot plot with the indicated mean cell diameter. Statistical significance was calculated using one-way ANOVA with Dunnett’s multiple comparisons test (****, *P* <0.0001; ns, not significant) (WT, n=190; WT/WT, n=104; *far8*Δ*-1,* n=151; *far8*Δ*-2,* n=162; *far8*Δ::*FAR8,* n=140). (E, F) Virulence of WT (KN99**α**), *pph22*Δ, *pph22*Δ *sup, far8*Δ, and *far8*Δ::*FAR8* cells in a murine model of *C. neoformans* infection via the intranasal inhalation route. Equal numbers of male and female A/J mice (n=14 per group) were inoculated with 10^5^ cells and monitored for 45 days post-infection. (E) Survival rates of mice infected with the indicated strains. Ten mice per strain were analyzed. Dashed line indicates the time of fungal burden analysis (****, *P* <0.0001). (F) Brain and lungs were harvested from four randomly selected mice per group at 14 days post-infection to quantify CFU per gram of organ tissue (One-way ANOVA; ****, *P* <0.0001; ***, *P* <0.001; **, *P* <0.01; ns, not significant).

Capsule production was assessed in WT, *pph22*Δ, *pph22*Δ *sup, far8*Δ, and *far8*Δ::*FAR8* strains by growing cells in RPMI medium at 30°C, and then analyzing capsule size with India ink (Fig 8C). The *pph22*Δ mutants grew very slowly in this nutrient-poor medium, produced no visible capsule, and formed elongated cells. The *pph22*Δ *sup* strains produced small capsules with normal cell morphology (Fig 8C, top panel). The *far8*Δ strains produced capsules relatively similar in size to the wild type, but the cells were significantly larger, appearing as titan cells, which are enlarged cells greater than 15 μm in diameter [55, 56]. Cells larger than 10 μm accounted for ∼30% of the total cell population. This experiment was repeated to include a WT/WT diploid control, ruling out the possibility that the large cell size was due to the diploid nature of the *far8*Δ strains. Cells were grown in RPMI media for two days and their cell diameters were quantified (Fig 8D). Both *far8*Δ*-1* and *far8*Δ*-2* strains produced significantly larger cells than the WT haploid and diploid control cells, with diameters ranging from around 5 to 23 μm. Subsequent FACS analysis showed that *far8*Δ cells from RPMI were still primarily diploid (peaks at 2C and 4C) and did not become polyploid, which is typical of titan cells (Fig S9). Therefore, we can conclude that deletion of *FAR8* leads to an abundant production of diploid titan cells in nutrient-limiting conditions.

The virulence of *pph22*Δ and *far8*Δ mutants was tested in a murine infection model. As expected, given their temperature-sensitive growth defects, *pph22*Δ, *pph22*Δ *sup,* and *far8*Δ mutants could not cause disease upon intranasal instillation in mice before the experiment was terminated after 45 days (Fig 8E). Four mice from each cohort were randomly selected and sacrificed on day 14 for fungal burden analysis in the brain and lungs (Fig 8F). The *pph22*Δ and *far8*Δ mutants could not persist in the lungs or disseminate to the central nervous system. However, cells from the *pph22*Δ *#1 sup1* strain were able to persist in the lungs and disseminate to the brain in one out of four infected mice, although at a significantly reduced level. These results indicate that *PPH22* and *FAR8* are required for the virulence of *C. neoformans*.

### Deletion of STRIPAK complex subunit *MOB3* leads to hypervirulence

Phenotypic analyses demonstrated that the *C. neoformans* STRIPAK complex is crucial for sexual development and multiple aspects of vegetative growth. The *pph22*Δ and *far8*Δ mutants exhibited dramatic phenotypes under different stress conditions. It was hypothesized that *mob3*Δ mutant strains might present similar phenotypes. However, *mob3*Δ mutants did not exhibit any growth defects compared to the wild-type strain on YPD media at 30°C or 37°C, on YNB, or in the presence of rapamycin, CFW, caffeine, FK506, or CsA (Figs 9A and S10). At a higher temperature of 39°C, the *mob3*Δ mutants grew significantly better than the wild type. Additionally, *mob3*Δ mutant strains also grew slightly better than the wild type at 37°C in the presence of 5% CO_2_.

**Fig 9.**
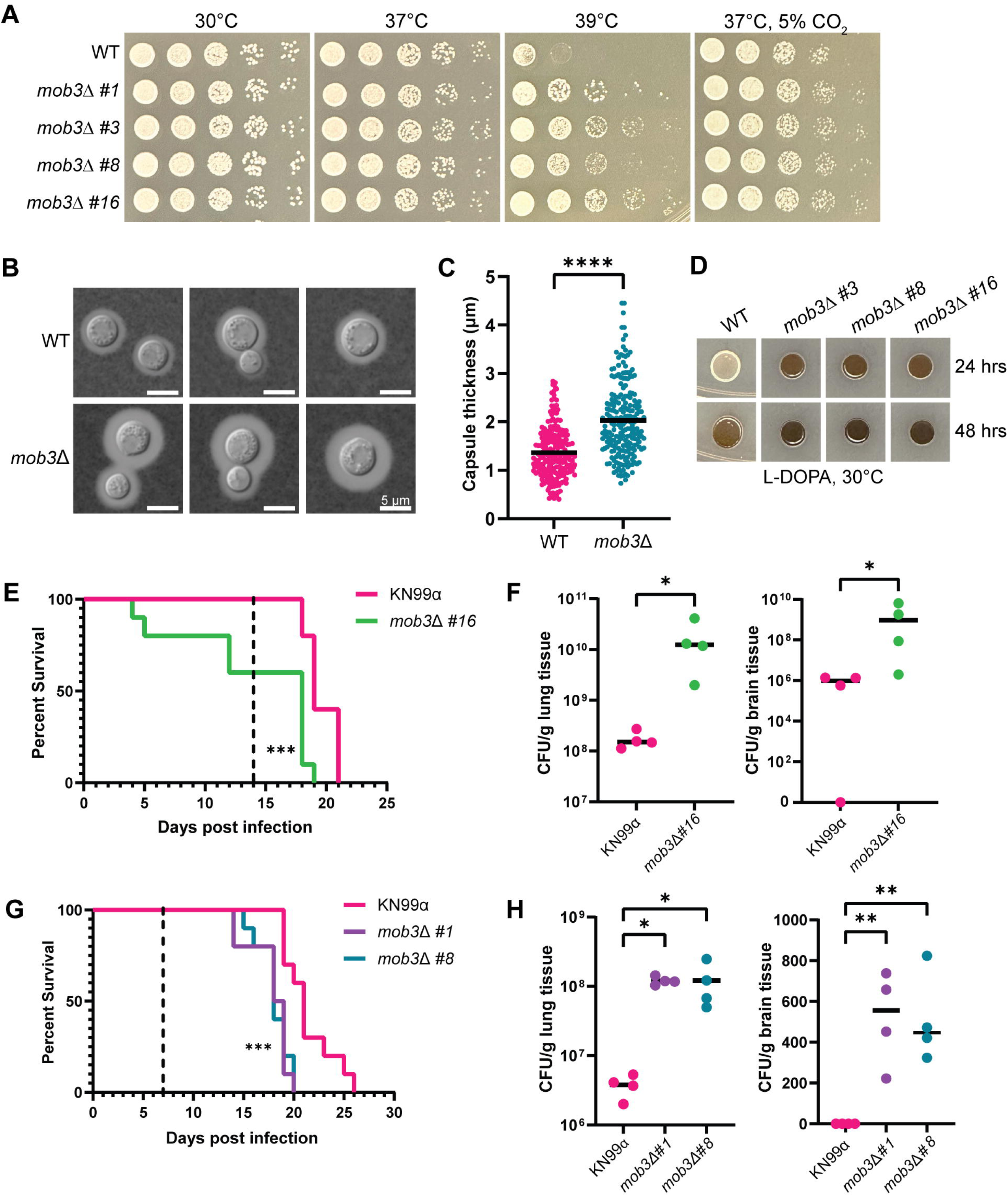
*mob3*Δ mutants are hypervirulent. (A) Thermotolerance of WT (KN99**a**) and isogenic *mob3*Δ strains. Cells were serially diluted, spotted onto YPD at 30°C, 37°C, 39°C, and 37°C with 5% CO_2_, and incubated for two days. (B) WT (KN99**a**) and *mob3*Δ cells were grown in RPMI at 37°C for three days, harvested, washed with PBS, and stained with India ink to analyze capsule formation. (C) Capsule thickness measurements were made by subtracting the cell body diameter from the capsule diameter and dividing by 2. Data are presented in a scatter dot plot with the mean capsule thickness indicated for each strain. Statistical analysis was performed using the Mann-Whitney U test (****, *P* <0.0001) (WT, n=214; *mob3*Δ, n=222). (D) Melanization of WT (KN99**a**) and *mob3*Δ strains on L-DOPA medium at 30°C after 24 and 48 hours of incubation. (E-H) Equal numbers of male and female A/J mice were infected intranasally with 10^5^ cells of the indicated WT and isogenic *mob3*Δ mutant strains and analyzed for survival rate (n=10) and fungal burden (n=4). (E) Survival analysis of WT (KN99α) and *mob3*Δ *#16* infected mice. Mice were 4 weeks old at the time of infection. The dashed line indicates the day at which fungal burden analysis was performed (***, *P* <0.001). (F) CFUs per gram of lung and brain tissue recovered from organs harvested at 14 days post-infection (Mann-Whitney U test; *, *P*=0.014). (G) Survival rates of 5-week-old mice infected with WT (KN99α), *mob3*Δ *#1,* and *mob3*Δ *#8* strains. The dashed line represents the day post-infection of fungal burden analysis (***, *P* <0.001). (H) CFUs per gram of lung and brain tissue recovered from organs harvested at 7 days post-infection (One-way ANOVA; **, *P* <0.01; *, *P* <0.05).

Next, we investigated virulence factor production in the *mob3*Δ mutants. When examining the capsule, we noted an increase in capsule size in *mob3*Δ compared to wild type (Fig 9B). Quantification of capsule thickness from wild-type and *mob3*Δ cells revealed that *mob3*Δ produced significantly larger capsules, with some cells having capsules reaching nearly 4.5 μm (Fig 9C). For melanin production, *mob3*Δ strains produced significantly more melanin than the wild-type strain, which was visible after 24 and 48 hours of incubation on L-DOPA medium (Fig 9D). Collectively, these results indicate that the deletion of *MOB3* leads to increased heat and CO_2_ tolerance, and enhanced production of capsule and melanin, suggesting possible alterations in virulence potential.

The virulence of *mob3*Δ mutants was tested in a murine inhalation model of *C. neoformans* infection. Four-week-old mice were infected intranasally with *mob3*Δ *#16* strain or an isogenic KN99α wild-type strain. Animals infected with the *mob3*Δ *#16* strain exhibited significantly decreased survival, with several succumbing to death prior to 7 days post-infection (Fig 9E). The fungal burden of lung and brain tissues was assessed at 14 days post-infection and the *mob3*Δ mutation led to increased proliferation in the lungs and dissemination into the brain (Fig 9F).

To confirm that the changes in virulence were indeed due to the *mob3*Δ mutation, the infection experiment was repeated with two other independent mutant strains, *mob3*Δ *#1* and *mob3*Δ *#8.* For this experiment, five- to six-week-old mice were used to rule out any influence of smaller mouse size on disease progression. Fungal burden assays were performed at day 7 post-infection instead of day 14 to better understand how rapidly *mob3*Δ cells disseminate from the lungs of infected animals to the brain. The survival curves indicated that animals infected with *mob3*Δ *#1* and *mob3*Δ *#8* succumbed to death significantly faster than those infected with the wild type (Fig 9G). Fungal burden was also measured at higher levels for *mob3*Δ mutants in the lungs and brain tissue than the control. Notably, CFUs were obtained from the brains of all four mice in both *mob3*Δ *#1* and *mob3*Δ *#8* groups, whereas no CFUs were found in the brains of animals infected with wild-type KN99α at 7 days post-infection. These findings demonstrate that the *mob3*Δ mutation significantly enhances the virulence of *C. neoformans*.

### Transcriptional analyses of *pph22*Δ*, far8*Δ, and *mob3*Δ mutants

Collectively, our results demonstrated that deletion of *PPH22, FAR8,* and *MOB3* in *C. neoformans* led to significantly altered vegetative growth, response to temperature and stress, production of virulence traits, and pathogenicity. To address whether changes in gene expression could explain some of the observed phenotypes in *pph22*Δ*, pph22*Δ *suppressor, far8*Δ, and *mob3*Δ strains, we performed transcriptional profiling to identify genes regulated by the STRIPAK complex. Total RNA was extracted from cells grown in YPD media at 30⁰C and RNA sequencing was performed with poly(A) enrichment for mRNAs. The expression data gathered for all differentially expressed genes was filtered for genes with significant changes (log_2_ fold change greater than 1.5 or less than -1.5 and adjusted *p* value less than 0.05) among four comparison groups: 1) *pph22*Δ vs. WT; 2) *pph22*Δ *suppressor* vs. WT; 3) *far8*Δ vs. WT; and 4) *mob3*Δ vs. WT (Table S3). *pph22*Δ, *pph22*Δ *sup,* and *far8*Δ mutants exhibited similar proportions of genes that were significantly differentially expressed out of the total number of DEGs, whereas the *mob3*Δ mutant had a much lower number of significantly expressed genes. Fig 10A shows the number of upregulated (left) and downregulated (right) genes shared and unique among the four deletion mutant groups compared to the wild-type control. Surprisingly, there was minimal overlap in the up- or down-regulated genes in *pph22*Δ and *far8*Δ mutants, despite both encoding subunits of the PP2A complex and sharing several phenotypic traits. Together, the four mutant groups (*pph22*Δ, *pph22*Δ *sup, far8*Δ, and *mob3*Δ) shared six genes with increased expression, all of which are annotated as hypothetical proteins, and zero genes with decreased expression. There were 111 total common genes significantly upregulated among *pph22*Δ, *pph22*Δ *sup,* and *far8*Δ mutant groups. Notable genes with potential biological significance include those involved in iron and metal homeostasis, such as *SIT2* and a *CDF* zinc transporter, the chitin synthase *CHS7*, the DNA replication regulator *CDC6*, and the DASH complex subunits *DAD1* and *DAD3*, which play roles in kinetochore-microtubule interactions. Additionally, three genes annotated as membrane transporters or drug efflux pumps were upregulated in these groups (Table S4). Conversely, there were 128 common genes that were downregulated among *pph22*Δ, *pph22*Δ *sup,* and *far8*Δ mutants, including many genes involved in core metabolic processes, such as aconitase *ACO1,* galactokinase *GAL1,* fatty acid synthases *FAS1* and *FAS2,* ATP citrate synthase subunit *ACL1,* and high-affinity glucose transporter *SNF3.* The decreased fitness observed in these mutant strains could be attributed to reduced carbohydrate and lipid metabolic processes. In the *mob3*Δ mutants, only 82 genes were found to be significantly differentially regulated, suggesting that the deletion of *MOB3* has a relatively modest impact on overall gene expression.

**Fig 10.**
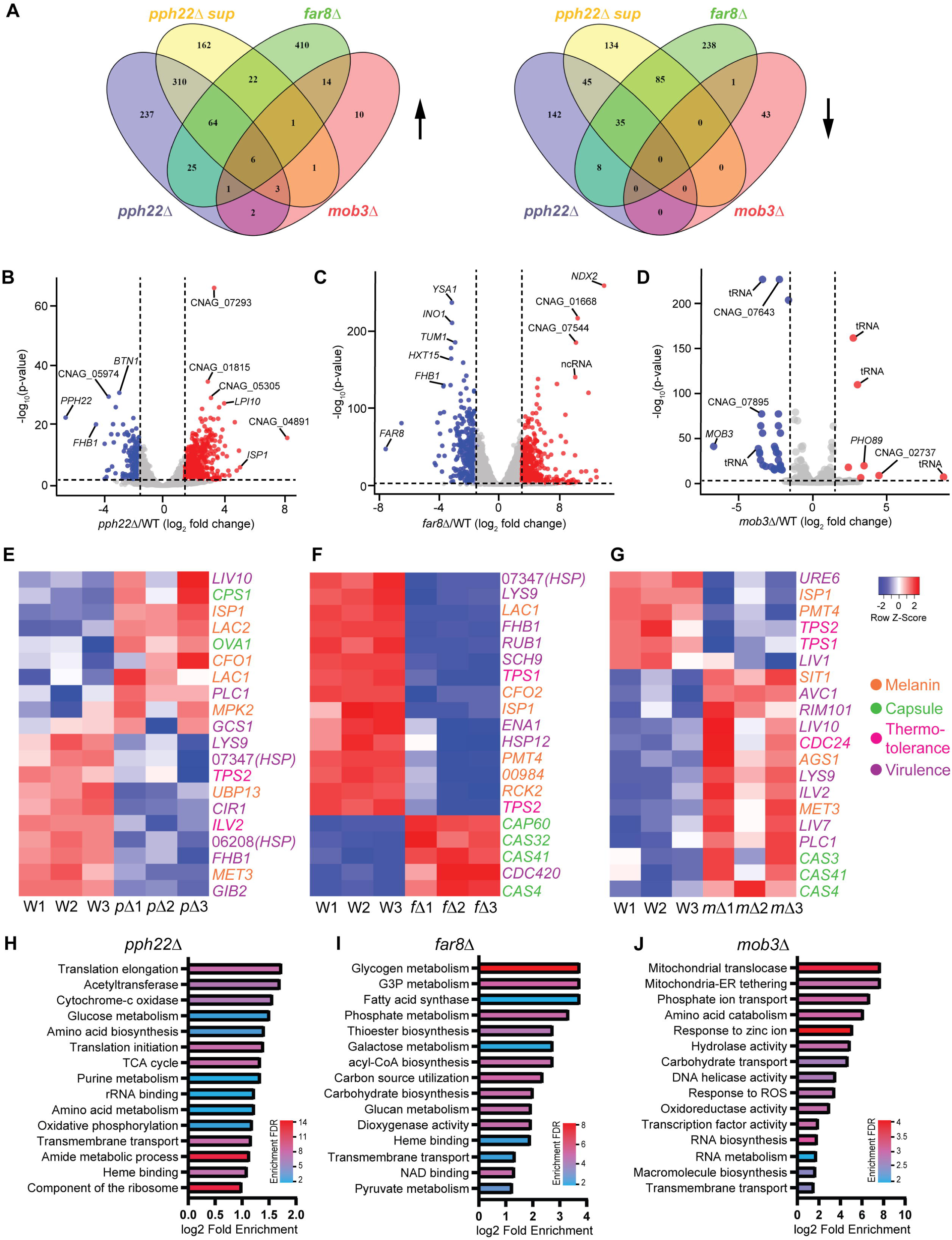
Transcriptome analysis of *pph22*Δ, *far8*Δ, and *mob3*Δ mutants. (A) Venn diagrams depicting shared differentially expressed genes (DEGs) among the indicated mutant strains for genes that were upregulated (left) or downregulated (right) compared to the isogenic wild-type control. Criteria for significant DEGs was log_2_ fold change >1.5 or <-1.5 and *P* adjusted <0.05. (B, C, D) Volcano plots of -log_10_ adjusted *P* value versus log_2_ fold change for all DEGs in *pph22*Δ (B), *far8*Δ (C), and *mob3*Δ (D) mutants. Significant genes are highlighted as downregulated (blue) or upregulated (red). The top 10 most significantly regulated genes are labeled in each panel. The vertical and horizontal lines denote the log_2_ fold change thresholds (-1.5 and 1.5) and -log_10_(adjusted *P* value) threshold (2.0), respectively. (E, F, G) Heatmaps of hierarchical clustering depicting normalized expression data for selected virulence genes in *pph22*Δ, *p*Δ (E), *far8*Δ, *f*Δ (F), *mob3*Δ, *m*Δ (G), and wild-type, W samples. Row Z scores are color coded according to the legend at the right. Transcriptomic data for each mutant was gathered in biological triplicate and plotted with an independent set of wild-type control samples. (H, I, J) Gene ontology (GO) enrichment analysis of GO terms overrepresented in the transcriptomes of *pph22*Δ (H), *far8*Δ (I), and *mob3*Δ (J) samples. The top 15 significant GO terms are ranked based on the log_2_ fold enrichment score compared to the background set (all genes in the genome) and color coded by enrichment false discovery rate (FDR) to show significance [-log_10_(adjusted *P* value)].

The transcriptomes of *pph22*Δ, *far8*Δ, and *mob3*Δ mutants also revealed genes that were highly dysregulated, with log_2_ fold changes >3 or <-3. In *pph22*Δ, a short-chain dehydrogenase/reductase *ISP1* and the multidrug transporter *LPI10* were highly upregulated, while *BTN1*, homologue of human *CLN3,* and *FHB1,* which encodes flavohemoglobin and is involved in virulence, were significantly downregulated (Fig 10E). Decreased expression of *FHB1* was also shared by *far8*Δ, along with hexose transporter *HXT15,* the inositol synthase *INO1,* and *YSA1,* which regulates oxidative stress response (Fig 10F)*. FHB1* and *YSA1* were also downregulated in *pph22*Δ *suppressors* (Fig S11), along with two genes encoding fructosamine kinase *IRK2* and another short-chain dehydrogenase/reductase *SPS19*. *PPG1* was the most significantly overexpressed gene in this group, supporting the hypothesis that increased *PPG1* expression may be the causative mutation driving suppression in *pph22*Δ mutants. The genes with the most significant log_2_ fold changes in *mob3*Δ mutants encoded either hypothetical proteins or tRNA molecules, with the exception of phosphate transporter *PHO89,* which was upregulated (Fig 10G). Interestingly, nearly half of the significant DEGs in this group encode tRNAs (Table S4), 17 of which are upregulated in *mob3*Δ but downregulated in *far8*Δ, suggesting *MOB3* and *FAR8* may have differential roles in tRNA synthesis or regulation.

To visualize more global changes in the transcriptomes of *pph22*Δ, *pph22*Δ *sup, far8*Δ, and *mob3*Δ mutants, normalized transcript counts of DEGs were filtered (log_2_ fold change >1.2 or <-1.2; *p* value <0.05), clustered using average linkage with Pearson distance measurement, and plotted in a heatmap along with isogenic wild-type controls (Fig S12). In *mob3*Δ samples, the criteria for filtering were less stringent due to the lower number of significant DEGs compared to wild type (log_2_ fold change >0.6 or <-0.6; *p* value <0.05). The heatmaps demonstrate a high degree of similarity among the three biological replicates within each group, indicating consistent gene expression patterns and supporting the robustness of the differential expression results between groups. Interestingly, although the *pph22*Δ suppressor strains, which overexpress *PPG1*, effectively mitigate many of the phenotypic defects seen in *pph22*Δ mutants, their gene expression profiles are strikingly similar to those of *pph22*Δ. This suggests that suppression of *pph22*Δ phenotypes may occur through mechanisms beyond broad transcriptional changes.

From the heatmaps, we selected a group of 106 genes that are known to be involved in virulence of *C. neoformans*, based on published literature on FungiDB, deletion of which have either reduced virulence factor production or an attenuated virulence phenotype (Table S5). 47 of the genes from this list were collected by Kim et. al. (2015), in which they were assigned a virulence category (capsule, melanin, or thermotolerance) [57]. Heatmaps were generated for *pph22*Δ, *far8*Δ, and *mob3*Δ mutants compared to wild type of the top 20 most significantly differentially expressed genes involved in capsule production, melanization, thermotolerance, or virulence (if the gene had reported functions in multiple virulence factors or an unknown role in affecting virulence) (Figs 10E, 10F, and 10G). *pph22*Δ and *far8*Δ samples shared several overlapping genes involved in production of virulence factors, including decreased expression of *LYS9,* heat shock protein CNAG_07347, and *TPS2*. Decreased expression of *TPS1* and *TPS2* was shared by *far8*Δ and *mob3*Δ, along with two genes involved in melanin production *PMT4* and *ISP1. mob3*Δ mutants displayed increased expression of the important virulence regulators *PLC1* and *RIM101.* Overall, *mob3*Δ exhibited a larger proportion of virulence-related genes that were overexpressed compared to *pph22*Δ and *far8*Δ groups. Interestingly, *pph22*Δ and *far8*Δ mutants showed both up- and down-regulation of the selected virulence-related genes. These results demonstrate baseline changes in gene expression outside of host-mimicking conditions, suggesting regulatory shifts that may influence virulence potential.

Finally, gene ontology enrichment analysis was performed for significant DEGs in *pph22*Δ, *far8*Δ, and *mob3*Δ mutants (Fig 10H, 10I, 10J and Table S6). In the downregulated genes in *pph22*Δ, there was enrichment of GO terms involved in protein synthesis, such as translation initiation and elongation, rRNA binding, and component of the ribosome, as well as in energy production, including glucose metabolism, cytochrome c oxidase, TCA cycle, and oxidative phosphorylation (Fig 10H). These results suggest that *pph22*Δ strains undergo metabolic remodeling and possibly reduce the activity of resource-intensive cellular processes. GO terms related to energy storage and utilization were also enriched among downregulated DEGs in *far8*Δ, including metabolism of glycogen, glycerol-3-phosphate, galactose, phosphate, and glucan (Fig 10I). Genes with GO terms related to heme binding were among the most significantly downregulated DEGs in *pph22*Δ and *far8*Δ, suggesting a reduction in pathways linked to electron transport and oxidative metabolism that depend on heme availability. Both groups also displayed enrichment for genes encoding transmembrane transporters (many of which are annotated as major facilitator superfamily transporters) among upregulated DEGs, and in *pph22*Δ, several of the most upregulated transporters encode multidrug efflux pumps. In *mob3*Δ, upregulated DEGs were enriched for GO terms related to mitochondrial function, such as mitochondrial translocase and mitochondrial-ER tethering, as well as terms related to cell growth and adaptive signaling, including RNA biosynthesis, RNA metabolism, response to zinc ion, and response to reactive oxygen species (ROS) (Fig 10J). This suggests that *mob3*Δ may activate pathways to respond and adapt to cell stresses. Genes with associated GO terms relating to transport (phosphate ion transport, carbohydrate transport, and transmembrane transport) were found to be both up- and down-regulated in *mob3*Δ mutants.

Our results show that the phenotypes of *pph22*Δ and *far8*Δ mutants largely differ from those of *mob3*Δ. Transcriptome analysis further suggests that the expression profiles of *pph22*Δ and *far8*Δ are more aligned with each other than with *mob3*Δ, supporting the hypothesis that Mob3 may act as an opposing regulator in the STRIPAK complex. Genes with expression changes opposite to those in *mob3*Δ may represent direct or indirect STRIPAK complex targets. We identified nine such genes by analyzing differentially expressed genes that were oppositely regulated in *mob3*Δ versus *pph22*Δ and *far8*Δ (Fig S13). Genes that are downregulated in *pph22*Δ and *far8*Δ but upregulated in *mob3*Δ are phosphate transporter *PHO84,* D-lactate dehydrogenase *DLD1,* glycine dehydrogenase *GCV2,* sulfate adenyltransferase and virulence-related gene *MET3,* and a sarcosine oxidase, suggesting STRIPAK may be directly involved in nutrient sensing and metabolism. Conversely, genes with the opposite expression pattern were a phosphoglycerate dehydrogenase, alcohol dehydrogenase *ADH1,* allantoate transporter *DAL5,* and a virulence-related gene *LIV1.* Taken together, these expression patterns suggest that STRIPAK may play a key role in balancing metabolic regulation with pathways involved in virulence, and potentially revealing a coordinated function within the complex.

## Discussion

The STRIPAK signaling complex is highly conserved across eukaryotic species and regulates numerous developmental processes. In this study, we characterized the organization of the *C. neoformans* STRIPAK complex and determined its role in genome stability, development, and virulence. The *C. neoformans* STRIPAK complex is composed of the PP2A subunits Tpd3, Far8, and Pph22, along with associated proteins Far9, Far11, and Mob3. Sequence analysis, structural modeling, and protein interaction analysis revealed that *C. neoformans* STRIPAK closely resembles the human ortholog. The predicted arrangement of STRIPAK components in *C. neoformans* is remarkably similar to that in mammalian systems, and our yeast two-hybrid assay detected similar interactions among STRIPAK components to what has been reported in other fungi. For example, a study with *S. macrospora* [9] also identified interactions between homologs of Far8 (Pro11) and Mob3, and Far11 (Pro22) and Tpd3. Similarly, AP-MS experiments in *N. crassa* showed that the Far9 homolog (HAM-4) interacts with Tpd3, and the Far11 homolog (HAM-2) associates with both Pph22 and Tpd3 [8], further supporting the conservation of core STRIPAK subunit interactions across diverse eukaryotes. The STRIPAK multimer structural prediction reveals a close interaction between the coiled-coil domains of Far8 and Far9, suggesting a unique assembly within *C. neoformans*. In humans, SLMAP (Far9) and SIKE form a regulatory STRIPAK subcomplex, a feature absent in the human STRIPAK core’s cryo-EM structure. [43]. The localization of fungal Far9-STRIPAK at the ER, nuclear envelope, or mitochondria supports its role in linking signal transduction pathways [58]. This structural variability suggests that membrane association of Far9 may govern the unique organization of STRIPAK in *C. neoformans*.

STRIPAK’s role in cell cycle control is widely conserved among eukaryotes, with both overlapping and distinct functions across species. Studies in filamentous and non-filamentous ascomycetes have demonstrated STRIPAK complex homologs play similar but unique roles in regulating cell proliferation during vegetative and sexual development [59]. In humans, STRIPAK dysfunction is correlated with genome instability and DNA damage and leads to enhanced proliferation and migration of cancer cells [60]. Here, we demonstrate a specific role for STRIPAK complex subunits Pph22, Far8, and Mob3 in maintaining genome stability in *Cryptococcus neoformans*. In a diploid, deletion of one copy of the *PPH22* or *MOB3* genes leads to both segmental and whole chromosome aneuploidy. The aneuploid chromosomes vary between mutant strains, based upon differences in the read coverage maps of *PPH22/pph22*Δ*, MOB3/mob3*Δ, *pph22*Δ, and *mob3*Δ isolates, with chromosome 13 being most frequently duplicated. The *far8*Δ mutants were generated in haploid backgrounds but underwent near whole-genome endoreplication to become primarily diploid, as supported by FACS analysis. Our data reinforce the conserved role of *C. neoformans* STRIPAK in maintaining genome stability. The aneuploidy observed in *pph22*Δ, *far8*Δ, and *mob3*Δ mutants complicates phenotypic analysis; however, analysis of multiple independent mutants with differing chromosomal aneuploidy strengthens the hypothesis that the phenotypes described may be attributable to direct effects of *pph22*Δ, *far8*Δ, and *mob3*Δ mutations, rather than indirect effects of aneuploidy.

STRIPAK’s involvement in sexual development regulation is well-documented in fungi, and we uncovered a similar role in *C. neoformans.* Deletion mutations in *PPH22*, *MOB3*, and *FAR8* led to defects in sexual development when mated with the wild type, affecting hyphal initiation and elongation, basidia formation, and sporulation. Deletion of *PPH22* and *FAR8* orthologs in the filamentous fungi *S. macrospora* and *N. crassa* affects cell fusion, fruiting body formation, and septation, leading to obstruction of the sexual life cycle [9, 11, 61]. Filamentous ascomycetes also contain homologs of Mob3, which is absent in the ascomycetous yeasts like *S. cerevisiae* or *S. pombe*, suggesting a more specialized function in the sexual life cycle. In *Cryptococcus,* a basidiomycetous yeast, Mob3 appears to play a similar role in sexual development. Further study of the STRIPAK components in *C. neoformans* will address the mechanisms underlying STRIPAK regulation of sexual development.

Mutations in the PP2A catalytic subunit lead to severe growth defects or lethality in other fungal species [27] and *PPH22* has been assumed to be essential in *C. neoformans* [41]. We concluded that *PPH22* is not essential but *pph22*Δ does lead to highly reduced fitness during vegetative growth and under various stress conditions, likely due to severely diminished PP2A activity. However, interestingly, the growth defects observed in *pph22*Δ mutants were not only abolished by the addition of rapamycin, both *pph22*Δ and *pph22*Δ *sup* strains exhibited greater rapamycin resistance than the wild type. These findings demonstrate that inhibition of TORC1 effectively rescues growth defects due to *pph22*Δ, suggesting PP2A and TORC1 counteract each other’s activities. Another notable phenotype of *pph22*Δ mutants is their inability to grow on YPD with 1 M sorbitol. It is possible that the presence of sorbitol in YPD medium causes hyperosmotic stress, and *pph22*Δ cells are unable to activate an adequate stress response to maintain osmotic homeostasis. The High-Osmolarity Glycerol (HOG) pathway, well-characterized in *S. cerevisiae* and *Cryptococcus*, is essential for cells to adapt to fluctuating osmotic conditions [62, 63]. *PPH22* may modulate the HOG pathway or coordinate a response with other stress signaling pathways to maintain cell growth during hyperosmotic stress. Our data highlight the importance of PP2A in regulating numerous growth processes in *C. neoformans*.

Another critical protein phosphatase in *C. neoformans* is protein phosphatase 2B, calcineurin, which is required for growth at high temperatures and virulence [64], and is the target of the immunosuppressive drugs FK506 and cyclosporine A (CsA) [65]. We found that deletion of PP2A catalytic and regulatory subunits, *PPH22* and *FAR8,* also led to an inability to grow at elevated temperatures or in the presence of FK506 or CsA at 30°C, suggesting a synthetic lethal relationship with calcineurin inhibition. In fungal pathogens, calcineurin also plays key roles in host temperature tolerance, sexual development, morphological transitions, cell wall integrity, drug tolerance, and ER stress response [66]. It is possible that PP2A and calcineurin share some overlapping functions by dephosphorylating common targets, with one phosphatase partially compensating for the other’s loss. However, the fact that calcineurin is required for growth at high temperatures and that FK506 or CsA only inhibit cell growth at 37°C suggests that PP2A has distinct functions that are independent of calcineurin.

For *mob3*Δ mutants, apart from their mating defects, we did not observe any other significant fitness costs due to the deletion mutation, as seen in *pph22*Δ and *far8*Δ strains. Strikingly, both our *in vitro* and *in vivo* analyses demonstrate that *mob3*Δ mutants are hypervirulent. Deletion of *MOB3* leads to increased thermotolerance, CO_2_ tolerance, melanin production, and capsule production, all of which are important for survival in the host environment. These combined effects likely caused decreased survival of mice, elevated proliferation in the lungs, and increased brain dissemination in murine infection models. A role for *MOB3* in virulence or pathogenicity has not been described in studies on the STRIPAK complex in pathogenic ascomycetes, suggesting it may uniquely negatively regulate virulence in *Cryptococcus.* Despite increased virulence and pathogenicity due to *mob3*Δ deletion, there are likely other fitness trade-offs beyond the host environment in nature that could have limited the occurrence or persistence of mutations in this key gene during *Cryptococcus* evolution. Further study is needed to explore host-pathogen interactions involving *mob3*Δ mutants.

While *C. neoformans* is primarily a haploid yeast species, relatively stable diploid isolates have been described and used in genetic studies to elucidate gene functions [67–70]. One limitation of previous studies analyzing heterozygous diploid gene deletion mutants by sporulation and dissection is the nature of the parental diploid isolates. In most cases, these diploids have been selected by fusing complementary auxotrophic parental isolates, resulting in a prototrophic diploid, but with auxotrophic mutations (such as *ade2* and *ura5*) segregating in the haploid F1 progeny. Additionally, in some cases, such as the *C. neoformans* diploid strain AI187, one of the parental isolates resulted from UV irradiation of strain H99α and contains numerous nucleotide variants throughout the genome, attributable to UV mutagenesis [71, 72]. Thus, analyzing gene function based on the segregation of F1 haploid progeny from a heterozygous deletion mutant in the AI187 background has limitations due to the segregation of *ade2*, *ura5*, and additional heterozygous genetic factors. In this study, we capitalized on the generation of a new diploid strain (CnLC6683), resulting from a spontaneous fusion of two congenic strains, KN99α and KN99**a**, which has no auxotrophic mutations or mutations introduced by mutagenesis. This approach provides a robust platform for analyzing essential genes, as well as genes that are critical for cell growth and challenging to delete in haploid genetic backgrounds.

In our analysis of RNA-seq data, we observed a limited overlap in DEGs among the STRIPAK complex mutants *pph22*Δ, *far8*Δ, and *mob3*Δ, highlighting the distinct regulatory roles of these proteins within the complex. It is likely that *PPH22* functions more generally outside of the STRIPAK complex, influencing a wider array of cellular processes, while *FAR8* plays a more specialized role in STRIPAK-specific signaling. The unique roles of *PPH22*, *FAR8*, and *MOB3* may account for the divergent transcriptional effects, as well as phenotypic differences *in vitro* and *in vivo* that were observed in the mutants. The distinct transcriptional profiles of the three STRIPAK complex mutants warrant further investigation to elucidate both its overall function and the specialized roles of individual subunits in regulating cellular processes. Notably, each mutant exhibited significant changes in the expression of genes associated with virulence, even under baseline conditions. In *pph22*Δ and *far8*Δ, genes important for thermotolerance and regulation of virulence were downregulated which may explain their increased susceptibility to thermal stress and attenuated virulence in an animal infection model. The transcriptomes in these mutants also displayed enrichment for functions in energy metabolism and protein synthesis among the downregulated gene sets, suggesting reduced activity in these pathways may cause the growth defects observed in *pph22*Δ and *far8*Δ. Additionally, *pph22*Δ mutants were characterized by increased expression of genes encoding transmembrane transporters and drug efflux pumps, which may contribute to the strain’s resistance to fluconazole and rapamycin. In contrast, *mob3*Δ mutants showed increased expression for the majority of the selected virulence-associated genes and enrichment for functions related to mitochondrial activity and stress response, aligning with their enhanced production of virulence factors and hypervirulence in a murine model of *C. neoformans* infection. Collectively, the transcriptomic findings offer insights into the underlying molecular mechanisms linked to the observed phenotypes in *pph22*Δ, *far8*Δ, and *mob3*Δ strains.

The data in this study demonstrate that *C. neoformans* STRIPAK is vital for genome stability, sexual development, and virulence. Functional characterizations also reveal both shared and unique roles for STRIPAK components Pph22, Far8, and Mob3 in regulating cell growth, stress responses, and virulence processes. These results are consistent with its established function as a signaling hub in other organisms. However, the regulators and downstream effectors of the STRIPAK complex remain unclear. Investigations into the mechanisms through which STRIPAK can modulate various cellular processes may broaden our understanding of its comprehensive functions across diverse eukaryotes. While the structural similarities between the *C. neoformans* STRIPAK complex and its human counterpart suggest potential challenges for selective targeting, our findings highlight specific roles of individual subunits in virulence. For example, the fact that deletion of *MOB3* uniquely enhances virulence could guide subunit-specific targeting strategies. This finding, combined with future functional studies, could help refine STRIPAK’s viability as a therapeutic target. Phospho-proteomic analyses and investigation into the subcellular localization of its components, will provide insights into how STRIPAK regulates key signaling pathways in this important human fungal pathogen, as well as in other fungal species and beyond.

## Materials and methods

### Ethics statement

All animal experiments in this manuscript were approved by the Duke University Institutional Animal Care and Use Committee (IACUC) (protocol #A098-22-05). Animal care and experiments were conducted according to IACUC ethical guidelines.

### Strains, media, and growth conditions

*C. neoformans* strains used in this study are listed in Table S1. Strains were stored as 20% glycerol stocks at -80°C. Fresh cultures were revived and maintained on YPD (1% yeast extract, 2% Bacto Peptone, 2% dextrose) agar medium at 30°C. *Cryptococcus* transformants were selected on YPD medium supplemented with 100 μg/mL nourseothricin (NAT) or 200 μg/mL neomycin (G418). Strains were grown in YPD, synthetic complete (SC) (0.67% yeast nitrogen base with ammonium sulfate and without amino acids, 0.2% amino acid drop-out mix, 2% dextrose), or RPMI 1640 (Sigma-Aldrich R1383, 2% dextrose) liquid medium at either 30°C or 37°C, as indicated. For plate growth assays, strains were cultivated on YPD, Murashige and Skoog (MS) (Sigma-Aldrich M5519), YNB (0.67% yeast nitrogen base with ammonium sulfate and without amino acids, 2% dextrose), Niger seed (7% Niger seed, 0.1% dextrose), or L-3,4-dihydroxyphenylalanine (L-DOPA) (7.6 mM L-asparagine monohydrate, 5.6 mM glucose, 22 mM KH_2_PO_4_, 1 mM MgSO_4_.7H_2_O, 0.5 mM L-DOPA, 0.3 μM thiamine-HCl, 20 nM biotin, pH 5.6). All plate media were prepared with 2% Bacto agar. To induce copper sufficiency or deficiency, L-DOPA plates were supplemented with 10 μM CuSO_4_ or 10 μM of the copper chelator bathocuproine disulfonate (BCS). To analyze cell wall-associated phenotypes, sorbitol (1 M), caffeine (0.5 mg/mL), and calcofluor white (3 mg/mL) were added to YPD medium. To analyze cell growth in response to immunosuppressive agents, rapamycin (100 ng/mL), FK506 (1 μg/mL), and cyclosporine A (100 μg/mL) were added to YPD medium. Fluconazole Etest was performed using 0.016-256 μg/mL MIC test strips (Liofilchem 921470). For capsule analysis, strains were incubated for 2 days in RPMI media at 30°C or 37°C, followed by negative staining with India ink. For serial dilution assays, fresh cells were diluted to a starting OD_600_ of 0.1, serially diluted 20-fold, and spotted onto plates for the indicated media and temperature conditions. Plates were incubated for 2 to 7 days and photographed daily.

### Generation of marker-free *Cryptococcus neoformans* diploid strain CnLC6683

Wild-type KN99α and KN99**a** cells were mixed in equal numbers and incubated on V8 medium for 24 hours at room temperature in the dark to allow cells to fuse. The cells were then harvested and plated on YPD to isolate single colonies, with 100-300 colonies per plate. Colonies from YPD were replica-plated onto filament agar medium (1X YNB without amino acids or ammonium sulfate, 0.5% glucose, 4% agar) and incubated at room temperature in the dark. Self-filamentous colonies were isolated, and single colonies were purified on filament agar. Clones with a stable self-filamentation phenotype on filament agar were recovered on YPD and maintained as yeast colonies. Colony PCR was performed to confirm the presence of both **a** and α mating types, and flow cytometry and whole genome sequencing were used to confirm the ploidy of the resulting diploid, marker-free strain CnLC6683 (Figs 3A and 3D).

### Construction of mutant strains

Deletion mutant strains were generated in the *C. neoformans* H99α, YL99**a,** or KN99**a**/KN99α backgrounds. To generate the deletion alleles, the nourseothricin acetyltransferase gene expression cassette (NAT) or the neomycin resistance gene expression cassette (NEO) were amplified from plasmids pAI3 and pJAF12, respectively. Approximately 1 kbp homologous 5′ and 3′ regions of the targeted genes were amplified from H99 genomic DNA. These homologous arms were assembled with the drug resistance marker with overlap PCR, as previously described [73, 74]. The *PPH22/pph22*Δ and *MOB3/mob3*Δ heterozygous mutant diploid strains were generated via CRISPR-Cas9-directed mutagenesis. A codon-optimized version of *CAS9* for *C. neoformans* was PCR-amplified from plasmid pBHM2403 with universal primers M13F and M13R [75]. The desired target sequences for the sgRNA constructs were designed using the Eukaryotic Pathogen CRISPR guide RNA/DNA Design Tool (EuPaGDT) with default parameters [76]. The selected 20-nt guide sequence was added to the primers. The *Cryptococcus* U6 promoter, 20-nt guide sequence, scaffold, and 6T terminator were assembled using single-joint PCR with plasmid pBHM2329 as a template [75]. Gene deletion, sgRNA, and Cas9 expression constructs were introduced into *Cryptococcus* cells via electroporation, as previously described [77]. To generate *far8*Δ mutants, we employed homologous recombination in the *C*. *neoformans* H99α and YL99**a** backgrounds, using gene disruption cassettes containing *NAT* or *NEO* markers. Mutants were constructed according to previously described methods [53, 78]. For the *far8*Δ::*FAR8-NEO* complemented strain, full-length *FAR8* was amplified from H99α genomic DNA and cloned into a pNEO plasmid via the Gibson assembly method. After confirming integration of the *FAR8* gene into the plasmid through sequencing analysis, the plasmid was linearized with enzymatic digestion (BglII) and targeted reintegration of *FAR8-NEO* at the native locus was performed via biolistic transformation. Stable transformants from the YPD+NAT or YPD+NEO selection plates were screened by diagnostic PCR to confirm cassette integration at the endogenous locus. Positive transformants from diagnostic PCR were further confirmed via Illumina whole genome sequencing, which demonstrated the absence of reads mapping to the open reading frame in the deletion mutants, reads mapping to both the open reading frame and the deletion allele in the heterozygous diploid mutants, and integration of the *FAR8-NEO* construct at the endogenous locus in the *far8*Δ::*FAR8-NEO* complemented strain. Primers used in this study are shown in Table S2.

### Yeast two-hybrid assay

DNA segments encoding Pph22, Tpd3, Far8, Far9, Far11, and Mob3 were amplified by PCR from *C. neoformans* H99 cDNA and cloned into the BamHI and EcoRI restriction sites of pGADT7, a Gal4 transcriptional activation domain vector, and pGBKT7, a Gal4 DNA-binding domain vector (Takara Bio). Yeast two-hybrid strains Y187 (Clontech Laboratories) and Y2HGold (Takara Bio) were transformed with plasmids expressing GAD and GBD constructs, respectively, via the high-efficiency method [79]. The transformed strains were crossed to generate diploid cells co-expressing GAD and GBD fusion constructs. For the analysis of *ADE2* and *HIS3* reporter gene expression under the control of the Gal4-dependent promoter, transformants were grown on synthetic dextrose (SD) dropout medium (0.67% yeast nitrogen base, 2% dextrose) minus leucine, tryptophan, or histidine. On media lacking histidine (to select for Gal4-dependent expression of both *HIS3* and *ADE2* reporter genes), adenine sulfate was included at a low concentration of 10 mg/L, compared to the 100–150 mg/L typically found in standard yeast media. This limited adenine concentration helps reduce background growth, allowing for clear detection of strong protein-protein interactions that activate *ADE2* expression. The competitive inhibitor 3-amino-1,2,4-triazole (3-AT) was added at a concentration of 3 mM to limit *HIS3* activity. Amino acids and uracil were added at standard concentrations to support auxotrophic growth requirements.

### Flow cytometry analysis

Fluorescence-activated cell sorting (FACS) analysis to determine *Cryptococcus* ploidy was performed as previously described [80], with some modifications. Wild-type and mutant strains were grown on YPD medium at 30°C overnight, harvested, and washed with PBS. The cells were then fixed in 70% ethanol for 16 hours at 4°C. Fixed cells were pelleted and washed with 1 mL NS buffer (10 mM Tris-HCl, 0.25 M sucrose, 1 mM EDTA, 1 mM MgCl_2_, 0.1 mM ZnCl_2_, 0.4 mM phenylmethylsulfonyl fluoride, and 7 mM β-mercaptoethanol). After centrifugation, the cells were treated with RNase (0.5 mg/mL) and stained with propidium iodide (10 μg/mL) in a 200 μL suspension of NS buffer for 2 hours in the dark. Then, 50 μL of the stained cells were diluted into 2 mL of 50 mM Tris-HCl, pH=8.0, and submitted to the Duke Cancer Institute Flow Cytometry Shared Resource for analysis. Fluorescence was measured using a BD FACSCanto flow cytometer and analyzed with BD FACSDiva software. Approximately 15,000 events were analyzed for each sample.

### Scanning electron microscopy (SEM) and microscopic quantification

For sample preparation for SEM from self-filamenting diploid strains, an agar slice of the plated cells was fixed in a solution of 4% formaldehyde and 4% glutaraldehyde for 16 hours at 4°C. The fixed cells were then gradually dehydrated in a graded ethanol series (30%, 50%, 70%, and 95%), with a one-hour incubation at 4°C for each concentration. This was followed by three washes with 100% ethanol, each for 1 hour at room temperature. The samples were further dehydrated using a Ladd CPD3 Critical Point Dryer and coated with a layer of gold using a Denton Desk V Sputter Coater (Denton Vacuum, USA). Hyphae, basidia, and basidiospores were observed with a scanning electron microscope with an EDS detector (Apreo S, ThermoFisher, USA).

Brightfield and differential interference contrast (DIC) microscopy images were visualized with an AxioScop 2 fluorescence microscope and captured with an AxioCam MRm digital camera (Zeiss, Germany). Consistent exposure times were used for all images analyzed. Cell body sizes were measured using the measurement tool in ImageJ/Fiji. The thickness of polysaccharide capsules were calculated using the Quantitative Capture Analysis program, which uses the exclusion zone generated by India ink to differentiate the capsule from the cell body of individual cells [81]. Statistical differences in cell body diameter between groups were determined by one-way ANOVA with Dunnett’s multiple comparisons test, and the statistical difference in capsule thickness was determined using an unpaired *t*-test.

### Self-filamentation and mating analysis

To monitor self-filamentation efficiency, the wild-type diploid strain CnLC6683 (KN99**a**/KN99α) and the indicated heterozygous mutant diploid strains were grown overnight in YPD liquid media. The cultures were diluted to an OD_600_ of 1.0, and 4 μL was spotted onto MS plates. The plates were incubated at room temperature in the dark and monitored for signs of filamentation and sporulation for at least 4 weeks. For mating analyses, strains were grown overnight in YPD liquid media. *MAT***a** cells were mixed with *MAT*α cells in equal amounts and spotted onto MS plates. For crosses involving *pph22*Δ and *far8*Δ strains, mutant and wild-type cells were spotted in a 10:1 ratio to compensate for the growth defects of *pph22*Δ and *far8*Δ on MS media. First, *pph22*Δ and *far8*Δ mutant cells grown overnight in YPD liquid media were spotted onto MS plates and allowed to pre-grow for two days without the presence of the wild-type partner. After this pre-incubation, wild-type cells of the opposite mating type were spotted on top of the mutant cells. A H99α x KN99**a** cross served as a wild-type control on each mating plate. Mating efficiencies between groups were compared based on data obtained from at least 4 biological replicates.

### Whole-genome sequencing, ploidy, and SNP analysis

Genomic DNA for whole-genome sequencing was extracted from saturated 4 mL YPD cultures with the MasterPure Yeast DNA Purification Kit (LGC Biosearch Technologies, MPY80200). The precipitated DNA was dissolved in 35 μL of 1x TE buffer (100 mM Tris-HCl, 10 mM EDTA, pH 8.0), and the concentration was estimated using Qubit. Illumina sequencing was performed at the Duke Sequencing and Genomic Technologies core facility (https://genome.duke.edu) with Novaseq 6000, providing 250 bp paired-end reads. The Illumina sequences were mapped to the H99α genome assembly using Geneious software. The resulting BAM files were converted to TDF format, and read coverage was visualized in IGV to estimate ploidy for each chromosome. To assess significant changes in read coverage for each chromosome within a sample, regions of the genome were colored based upon a *Z-*score >1.96 or <-1.96 and a *P-*value <0.05. *Z*-scores between -1.96 and 1.96 with a *P-*value >0.05 were considered statistically insignificant. For SNP calling, the Illumina sequences were mapped to the H99α genome assembly using the Geneious default mapper in five iterations. Variant calling was performed using mapped read files, with parameters set to a 0.9 variant frequency and a minimum of 100x coverage per variant. Illumina sequences from H99α and KN99**a**/KN99α (CnLC6683) served as controls for SNP calling analysis.

### Competition assay

KN99**a**/KN99α and *PPH22/pph22*Δ diploid strains were cultured overnight at 30°C in liquid YPD or YPD+NAT, respectively. Cells were adjusted to equal densities using OD_600_ measurements and mixed in equal numbers in a 4 mL YPD co-culture. The cell density at the onset of competition was confirmed by plating the mixed cell dilution on YPD+NAT to select for *PPH22/pph22*Δ mutants and on YPD to determine the total cell count. This plating process was repeated at 24 and 48 hours to calculate the cell density of each strain in the co-culture. The data presented are based on four biological replicates, each with three technical replicates.

### Murine infection model

*C. neoformans* inoculum was prepared by culturing cells in 5 mL YPD on a tissue culture roller drum at 30°C for approximately 16 hours. Cells were collected by centrifugation, washed twice with sterile phosphate-buffered saline (PBS), and the cell density was determined with a hemocytometer. The final cell concentration was adjusted to 4 x 10^6^ /mL in PBS. Four- to five-week-old A/J mice (Jackson Laboratory, USA) were utilized for the murine intranasal infection model (n=14 for each group, 7 male and 7 female). Mice were anesthetized with isoflurane and infected by intranasal instillation of 25 μL inoculum (10^5^ cells). Mice survival was monitored daily, and euthanasia was performed via CO_2_ exposure upon reaching humane endpoints, including greater than 20% weight loss, reduced grooming and mobility, or a hunched appearance. For fungal burden analysis, four mice (2 male, 2 female) from each group were randomly selected and euthanized via CO_2_ exposure at 7 or 14 days post-infection. The brain and lungs were dissected and homogenized in 1 mL sterile PBS using bead-beating. Organ homogenates were plated onto YPD agar containing antibiotics (100 μg/mL ampicillin, 30 μg/mL chloramphenicol) to isolate fungal colonies. Survival data were plotted using Kaplan-Meier curves and statistically analyzed through log-rank (Mantel-Cox) test. Statistical analyses of fungal burdens were performed using either Mann-Whitney U test or one-way ANOVA with Dunnett’s multiple comparisons test. Data plotting and analysis of mouse survival and fungal burden was performed with GraphPad Prism v 10.2.3.

### RNA sequencing and differential expression analysis

Cultures of H99α, KN99**a,** and three independent *pph22*Δ*, pph22*Δ *suppressor, far8*Δ, and *mob3*Δ mutant strains were grown in 30 mL YPD at 30°C to an OD_600_ of 0.8-1.0. Total RNA was isolated from harvested cells with a Qiagen RNeasy Plant Mini Kit (Cat. #74904) with on-column DNase digestion (Qiagen, Cat. #79254) according to the manufacturer instructions. Biological triplicates were harvested for each strain. The concentration and integrity of RNA samples was estimated using Qubit and Nanodrop. Stranded mRNA-sequencing with poly(A) enrichment was performed at the Duke Sequencing and Genomic Technologies core facility with Illumina NovaSeq X Plus, which provided 2 x 150 bp paired-end reads. Read files were mapped to the *C. neoformans* H99α reference genome and gene expression was quantified using RNA STAR and featureCounts [82, 83]. Differential expression fold change, *p* value, and adjustment for multiple testing with the Benjamini-Hochberg procedure to control false discovery rate was determined with DEseq2 [84]. Gene count tables were normalized using variance stabilizing transformation (VST). For visualizing changes in gene expression, heatmaps and volcano plots of significant differentially expressed genes were generated using Heatmapper and VolcanoseR [85, 86]. Each set of mutant samples was compared to an independent set of wild-type controls, with all comparisons conducted as separate experiments. Gene Ontology (GO) terms for each differentially expressed gene set were analyzed using the built-in Gene Ontology Enrichment tool on FungiDB. Significantly enriched GO terms were plotted with GraphPad Prism 10.2.3.

## Acknowledgements

PPP is supported by NIH/NIAID T32 grant AI052080-20 as a Tri-I MMPTP fellow. This work is also supported by NIH/NIAID R01 grants AI039115-27, AI050113-20, and AI172451-02 awarded to JH. LEC is supported by the Canadian Institutes of Health Research (CIHR) Foundation grant (FDN-154288) and is a Canada Research Chair (Tier 1) in Microbial Genomics & Infectious Disease. JH and LEC are co-directors of the CIFAR Fungal Kingdom: Threats & Opportunities program. This work is also supported by National Research Foundation funded by the Korean government (MSIT) (2021R1A2B5B03086596, 2021M3A9I4021434, and 2018R1A5A1025077 to YSB). We thank Dr. Vikas Yadav, Dr. Maria Isabel Navarro-Mendoza, and Dr. Zhengchang Liu for their critical reading of this manuscript. We also thank laboratory manager Anna Floyd Averette for constant support, Dr. Corinna Probst for providing the *cbi1*Δ *ctr4*Δ strain, and Dr. Erica Washington for guidance on protein structure 3D modeling. We are grateful to Dr. Ulrich Kück for his advice and expertise. We commend the Duke Sequencing and Genomic Technologies Core Facility and the core’s director Dr. Devi Swain Lenz for their assistance.

