## Supplemental figures for "The *Cryptococcus neoformans* STRIPAK complex controls genome stability, sexual development, and virulence": S1 Fig. Y2H assay of STRIPAK components.docx

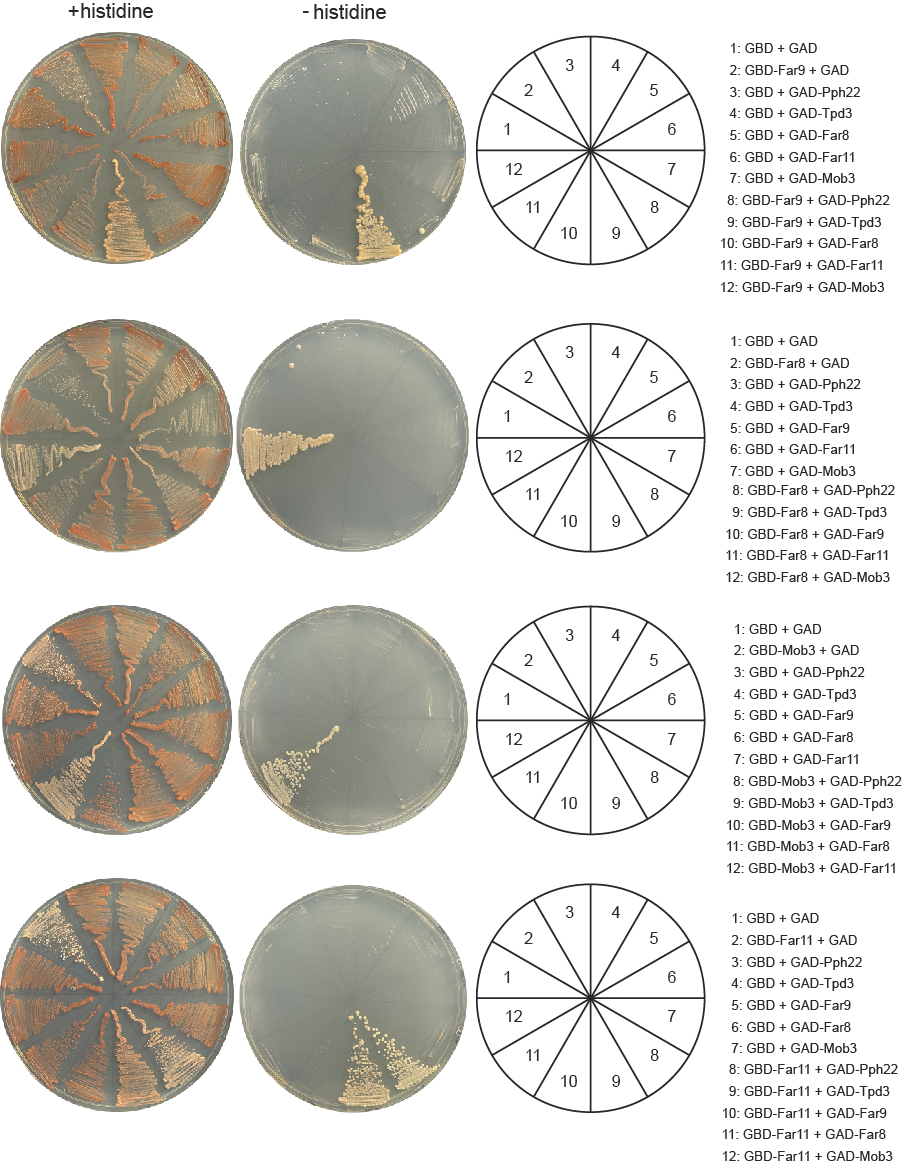
**A.**

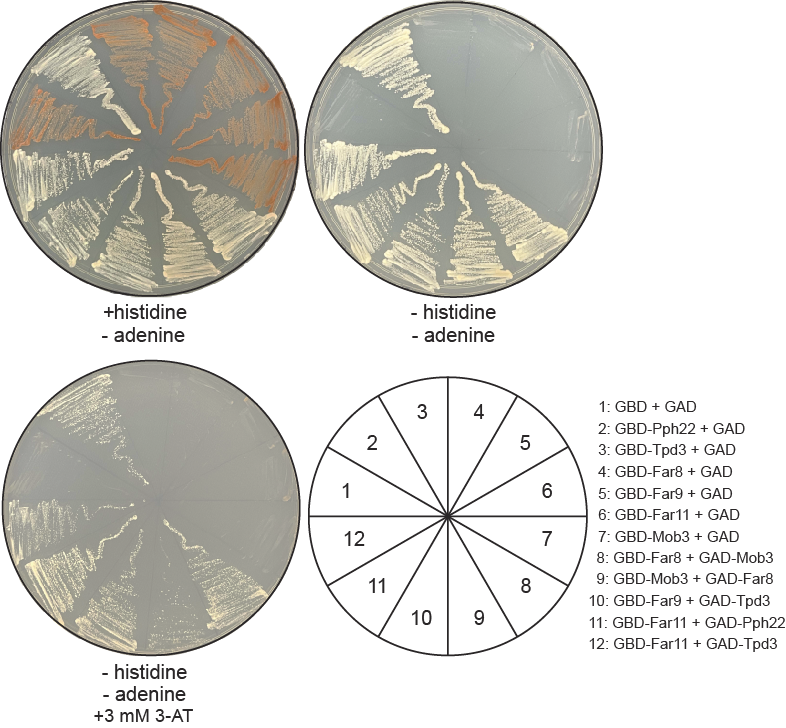
**B.**

**S1 Fig. Protein-protein interactions among STRIPAK components detected by yeast two-hybrid assay.**

(A) Yeast two-hybrid analyses of a 6x6 crossing of strains co-expressing plasmids encoding Gal4 DNA-binding and Gal4 transcription activation domains, fused to STRIPAK complex subunits. Expression of reporter genes *ADE2* and *HIS3*, indicating a positive interaction between GBD and GAD fusion proteins, makes cells appear less red on medium with histidine (Ade+) and able to grow on medium deficient in both histidine and adenine. (B) Confirmation of positive protein-protein interactions on media deficient in histidine and adenine, and on media with 3 mM 3-amino-1,2,4-triazole (3-AT). Autoactivation of the reporter genes in GBD-Pph22 allows for growth on media without histidine.
