## Supplemental figures for "The *Cryptococcus neoformans* STRIPAK complex controls genome stability, sexual development, and virulence": S1 Table. Strains used in this study.docx

| **Strain name** | **Description** | **Source/Reference** |
| --- | --- | --- |
| H99α | Wild-type *MAT*α | [1] |
| KN99**a** | Wild-type *MAT***a** | [2] |
| KN99α | Wild-type *MAT*α | [2] |
| YL99**a** | Wild-type *MAT***a** | [3] |
| CnLC6683 | Wild-type diploid (KN99**a**/KN99α) | Ci Fu and Leah Cowen |
| Y187 | *MAT*α yeast two-hybrid reporter strain | Takara Bio |
| Y2HGold | *MAT***a** yeast two-hybrid reporter strain | Takara Bio |
| PP68 | CnLC6683 *PPH22/pph22*Δ*::NAT-1* | This study |
| PP69 | CnLC6683 *PPH22/pph22*Δ*::NAT-2* | This study |
| PP70 | CnLC6683 *PPH22/pph22*Δ*::NAT-3* | This study |
| PP71 | CnLC6683 *PPH22/pph22*Δ*::NAT-4* | This study |
| PP72 | CnLC6683 *MOB3/mob3*Δ*::NAT-1* | This study |
| PP73 | CnLC6683 *MOB3/mob3*Δ*::NAT-2* | This study |
| PP74 | CnLC6683 *MOB3/mob3*Δ*::NAT-3* | This study |
| YSB9100 | H99α *far8*Δ*::NAT-1* | This study |
| YSB9102 | H99α *far8*Δ::*NAT-2* | This study |
| YSB9103 | H99α *far8*Δ::*NAT-3* | This study |
| YSB9104 | H99α *far8*Δ::*NAT-4* | This study |
| YSB11129 | YL99**a** *far8*Δ::*NEO-3* | This study |
| YSB11131 | YL99**a** *far8*Δ*::NEO-4* | This study |
| YSB11132 | YL99**a** *far8*Δ*::NEO-5* | This study |
| YSB11135 | YSB9100*::FAR8-NEO* | This study |
| PP119 | *MAT***a** *pph22*Δ*::NAT* (progeny from PP69) | This study |
| PP120 | *MAT***a** *pph22*Δ*::NAT* (progeny from PP69) | This study |
| PP121 | *MAT***a** *pph22*Δ*::NAT* (progeny from PP70) | This study |
| PP53 | *MAT*α *pph22*Δ*::NAT-1* (progeny from PP71) | This study |
| PP55 | *MAT*α *pph22*Δ*::NAT-3* (progeny from PP71) | This study |
| PP56 | *MAT***a** *pph22*Δ*::NAT-2* (progeny from PP71) | This study |
| PP58 | *MAT***a** *pph22*Δ*::NAT-4* (progeny from PP71) | This study |
| PP59 | *MAT*α *pph22*Δ*::NAT-5* (progeny from PP71) | This study |
| PP60 | *MAT*α *pph22*Δ*::NAT-6* (progeny from PP71) | This study |
| PP113 | *MAT*α *mob3*Δ*::NAT-1* (progeny from PP73) | This study |
| PP114 | *MAT*α *mob3*Δ*::NAT-3* (progeny from PP73) | This study |
| PP115 | *MAT***a** *mob3*Δ*::NAT-8* (progeny from PP73) | This study |
| PP117 | *MAT***a** *mob3*Δ*::NAT-16* (progeny from PP73) | This study |
| PP80 | *pph22*Δ*-1 suppressor 1* | This study |
| PP81 | *pph22*Δ*-2 suppressor 2* | This study |
| PP82 | *pph22*Δ*-5 suppressor 1* | This study |
| PP83 | *pph22*Δ*-8 suppressor 1* | This study |
| PP84 | *pph22*Δ*-9 suppressor 1* | This study |
| PP85 | *pph22*Δ*-9 suppressor 2* | This study |
| PP86 | *pph22*Δ*-10 suppressor 1* | This study |
| PP87 | *pph22*Δ*-11 suppressor 1* | This study |
| MCD16 | H99α *lac1*Δ*::URA5* | [4] |
| DTY1020 | H99α *cbi1*Δ*::HYG ctr4*Δ*::NAT* | [5] |
