## Supplemental figures for "The *Cryptococcus neoformans* STRIPAK complex controls genome stability, sexual development, and virulence": S2 Fig. Basidiospore dissection and genotyping of progeny.docx

**A.**

| Strain | # spores dissected | # spores germinated | % germination | # NAT resistant progeny | % NAT resistance |
| --- | --- | --- | --- | --- | --- |
| *PPH22/pph22*Δ*-1* | 140 | 44 | 31% | 18 | 41% |
| *PPH22/pph22*Δ*-2* | 80 | 30 | 38% | 6 | 20% |
| *PPH22/pph22*Δ*-3* | 80 | 49 | 61% | 21 | 43% |
| *PPH22/pph22*Δ*-4* | 140 | 58 | 41% | 30 | 52% |
| *MOB3/mob3*Δ*-1* | 77 | 24 | 31% | 0 | 31% |
| *MOB3/mob3*Δ*-2* | 140 | 123 | 88% | 62 | 88% |
| *MOB3/mob3*Δ*-3* | 126 | 45 | 36% | 0 | 36% |

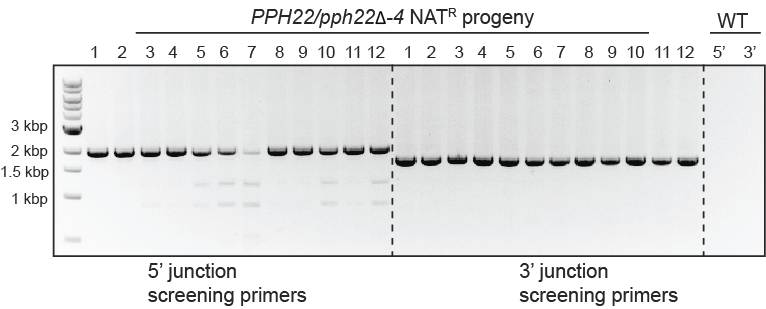

**B.**

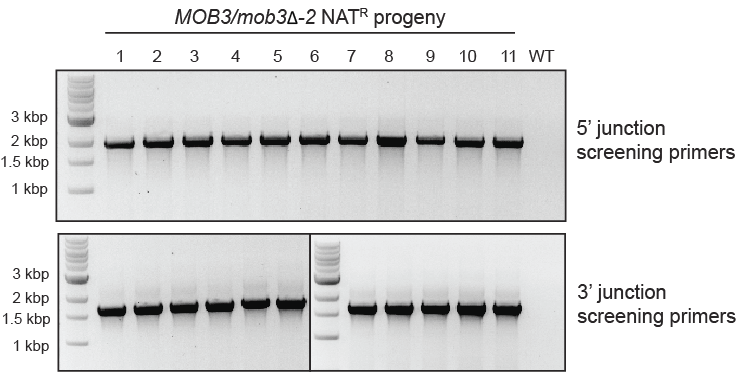

**C.**

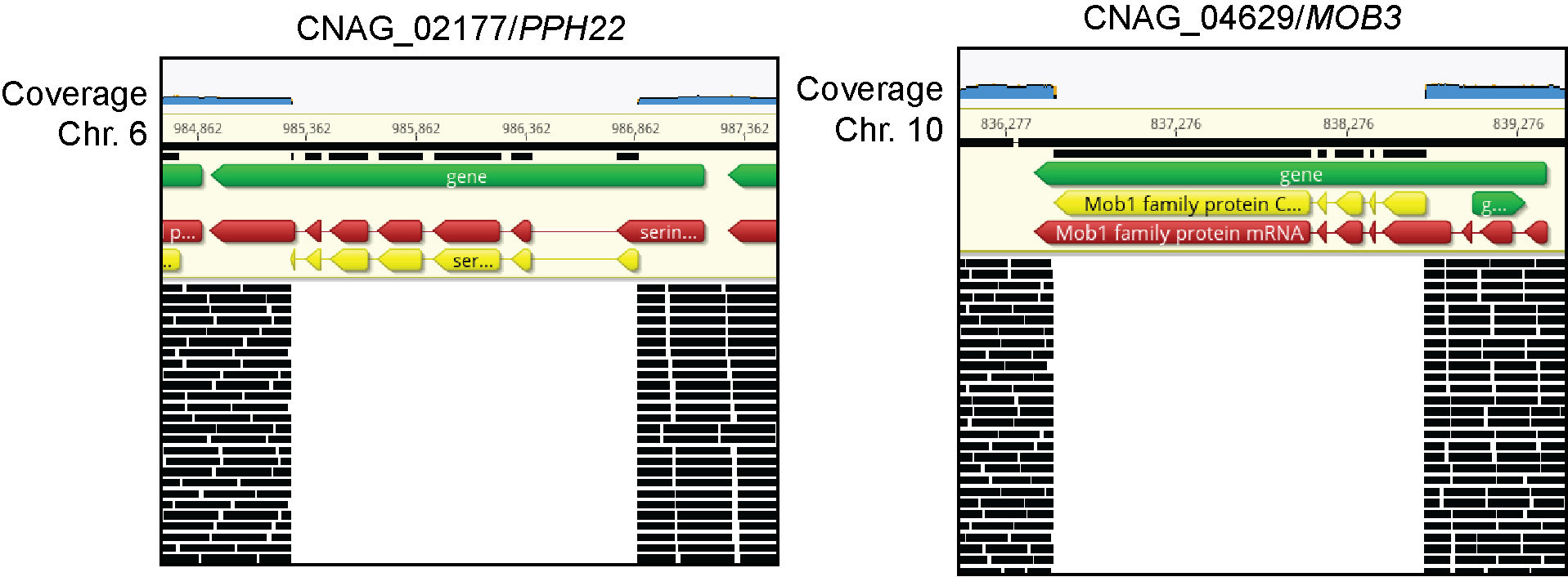
**D.**

**S2 Fig. Basidiospore dissection of *PPH22/pph22*Δ and *MOB3/mob3*Δ heterozygous diploid mutants and genotyping of progeny.**

(A) Table of basidiospore dissection results from *PPH22/pph22*Δ and *MOB3/mob3*Δ heterozygous diploid mutant strains in the CnLC6683 background. (B, C) PCR genotyping of 5’ and 3’ junctions of the integrated *NAT* deletion cassette in *pph22*Δ (B) and *mob3*Δ (C) progeny with the wild-type strain H99α used as a negative control. (D) Snapshots of Illumina whole-genome sequencing analysis in representative *pph22*Δ and *mob3*Δ mutants showing an absence of reads mapping to *PPH22* or *MOB3* open reading frames, respectively. The coding sequences (CDS) of each gene are depicted in yellow and individual DNA reads mapping to the H99α reference sequence are shown beneath in black. The read coverage for the region indicated by the chromosome coordinates is shown in blue.
