## Supplemental figures for "The *Cryptococcus neoformans* STRIPAK complex controls genome stability, sexual development, and virulence": S2 Table. Primers used in this study.docx

| **Primer #** | **Sequence** | **Purpose** |
| --- | --- | --- |
| M13F | GTAAAACGACGGCCAG | To amplify drug resistance cassettes |
| M13R | CAGGAAACAGCTATGAC | To amplify drug resistance cassettes |
| JOHE52463 | CTGGCGGAGGATAGAAGC | *ACT1* promoter reverse primer |
| JOHE52464 | GCGAATTCGAGACAGACATCG | *TRP1* terminator forward primer |
| JOHE52391 | CTGGCCGTCGTTTTACCTTGCTCTAGCTGTCTAAGACGCTGCTTTGAGGAA | *PPH22* deletion construct |
| JOHE52392 | GTCATAGCTGTTTCCTGTCGTTATCATCCCGAAACATTTTCTCAATCTCCCT |  |
| JOHE52401 | TTAACTTGCTTGGAGGTTTGC |  |
| JOHE52402 | TGCTCGGTCGACAGTGTAAG |  |
| JOHE52451 | AAAGTATAGCCAGCTCCTCGAACAGTATACCCTGCCGGTG | *PPH22* deletion guide RNA |
| JOHE52452 | CGAGGAGCTGGCTATACTTTGTTTTAGAGCTAGAAATAGCAAGTT |  |
| JOHE52393 | GCCTTAGAGCGACTGAATGG | *PPH22* deletion screening |
| JOHE52394 | ATCATTTCCGGGATTGTTGA |  |
| JOHE52407 | CTGGCCGTCGTTTTACTGTATGTTGGCTCTCGCTAAAGC | *MOB3* deletion construct |
| JOHE52408 | GTCATAGCTGTTTCCTGTGTGTTGTTGAATACGACATAGTG |  |
| JOHE52413 | CACGGTACCGGTGGAAGTAT |  |
| JOHE52414 | CAACAAAGCGACAACGAATG |  |
| JOHE52475 | GTATCTTCTCCCATGCCTACGTTTTAGAGCTAGAAATAGCAAGTT | *MOB3* deletion guide RNA |
| JOHE52476 | GTAGGCATGGGAGAAGATACAACAGTATACCCTGCCGGTG |  |
| JOHE52383 | TGTGTCTGTCAGCGGTCTTC | *MOB3* deletion screening |
| JOHE52384 | AGTGGGTCTTGTGTGGGAAG |  |
| B13314 | TCGTCCTTCGCTTGAAAACT | *FAR8* deletion construct |
| B13315 | TCACTGGCCGTCGTTTTACGTTGCTGACCTTGCTGTTGA |  |
| B13316 | CATGGTCATAGCTGTTTCCTGCCCAACGTGCATCCTTACTT |  |
| B13317 | TCTCGACCAGTTTGCTGATG |  |
| JOHE52379 | CTCTCCCGTCTTCTCGTTTG | *FAR8* deletion screening |
| JOHE52380 | GAGCCTGTCGCCTATGACTC |  |
| B20707 | GATGCATGCTCGAGCGGCCGCATTCCTGGGCACTCATTTTG | *FAR8* complementation construct |
| B20708 | TATCCATCACACTGGCGGCCGCTTTTGCCTCCAAAAAGGC |  |
| JOHE52477 | CATGGAGGCCGAATTCATGGCGGCCAATGACGATGTGGAT | *PPH22* yeast two-hybrid constructs |
| JOHE52478 | GCAGGTCGACGGATCCTTACAGGAAGTAGTCTGGGGGTCT |  |
| JOHE52489 | GGAGGCCAGTGAATTCATGGCGGCCAATGACGATGTGGAT |  |
| JOHE52490 | CGAGCTCGATGGATCCTTACAGGAAGTAGTCTGGGGGTCT |  |
| JOHE52479 | CATGGAGGCCGAATTCATGTCAGACCCGAACTCGGACCTC | *TPD3* yeast two-hybrid constructs |
| JOHE52480 | GCAGGTCGACGGATCCTTATGACAGGACTAACGGACAATA |  |
| JOHE52491 | GGAGGCCAGTGAATTCATGTCAGACCCGAACTCGGACCTC |  |
| JOHE52492 | CGAGCTCGATGGATCCTTATGACAGGACTAACGGACAATA |  |
| JOHE52483 | CATGGAGGCCGAATTCATGAACCGTGCCGTCCAGAGGAGT | *FAR8* yeast two-hybrid constructs |
| JOHE52484 | GCAGGTCGACGGATCCTCAAGCCAACCCCCAAAGTCTGAC |  |
| JOHE52495 | GGAGGCCAGTGAATTCATGAACCGTGCCGTCCAGAGGAGT |  |
| JOHE52496 | CGAGCTCGATGGATCCTCAAGCCAACCCCCAAAGTCTGAC |  |
| JOHE52481 | CATGGAGGCCGAATTCATGCAACATCAATACCTATCCCAT | *FAR9* yeast two-hybrid constructs |
| JOHE52482 | GCAGGTCGACGGATCCCTACTCCTTATGCTTATACCAGAA |  |
| JOHE52493 | GGAGGCCAGTGAATTCATGCAACATCAATACCTATCCCAT |  |
| JOHE52494 | CGAGCTCGATGGATCCCTACTCCTTATGCTTATACCAGAA |  |
| JOHE52485 | CATGGAGGCCGAATTCATGGCATCCTCAAAGCCAGTATTC | *FAR11* yeast two-hybrid constructs |
| JOHE52486 | GCAGGTCGACGGATCCCTACTCTCCGTAAGTATATTCTAC |  |
| JOHE52497 | GGAGGCCAGTGAATTCATGGCATCCTCAAAGCCAGTATTC |  |
| JOHE52498 | CGAGCTCGATGGATCCCTACTCTCCGTAAGTATATTCTAC |  |
| JOHE52487 | CATGGAGGCCGAATTCATGATAATACAGCCTCCCCAAGGG | *MOB3* yeast two-hybrid constructs |
| JOHE52488 | GCAGGTCGACGGATCCCTATGTCGTATTATACCTTTGCTT |  |
| JOHE52499 | GGAGGCCAGTGAATTCATGATAATACAGCCTCCCCAAGGG |  |
| JOHE52500 | CGAGCTCGATGGATCCCTATGTCGTATTATACCTTTGCTT |  |
