## Supplemental figures for "The *Cryptococcus neoformans* STRIPAK complex controls genome stability, sexual development, and virulence": S3 Fig. Confirmation of far8 deletion mutants.docx

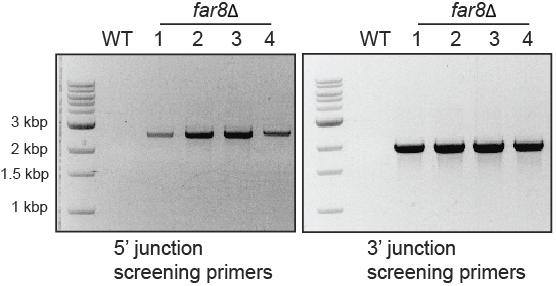
**A.**

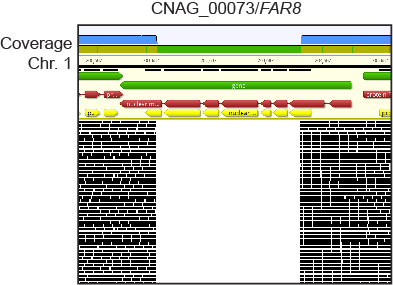
**B.**

**S3 Fig. Confirmation of *far8*Δ** **deletion mutants.**

(A) PCR genotyping of the 5’ and 3’ junctions for representative *far8*Δ mutants in the H99α background. WT (H99α) served as a negative control. (B) Snapshot of DNA read coverage at the *FAR8* locus from Illumina whole genome sequencing in a *far8*Δ mutant strain. The coding sequence is depicted in yellow and individual DNA reads are shown below in black. The *FAR8* gene was deleted from the middle of exon 1 to the middle of the last exon (exon 7), as verified by the absence of reads. The read coverage for the indicated chromosomal coordinates is shown in blue.
