## Supplemental figures for "The *Cryptococcus neoformans* STRIPAK complex controls genome stability, sexual development, and virulence": S3 Table. Comparison of DEGs from RNAseq.docx

**S3 Table. Differentially expressed genes in *pph22*Δ, *pph22*Δ *sup, far8*Δ, and *mob3*Δ**

**mutants from RNA sequencing analysis.**

| Comparison | Total DEGs | Up* | Down* |
| --- | --- | --- | --- |
| *pph22*Δ vs. WT | 6944 | 648 (9%) | 230 (3%) |
| *pph22*Δ *sup.* vs. WT | 6964 | 569 (8%) | 299 (4%) |
| *far8*Δ vs. WT | 8280 | 543 (7%) | 359 (4%) |
| *mob3*Δ vs. WT | 7984 | 38 (0.5%) | 44 (0.6%) |

* log_2_ fold change > 1.5 or < -1.5; *p* adjusted value < 0.05
