## Supplemental figures for "The *Cryptococcus neoformans* STRIPAK complex controls genome stability, sexual development, and virulence": S4 Fig. WGS and FACS analysis of mutants.docx

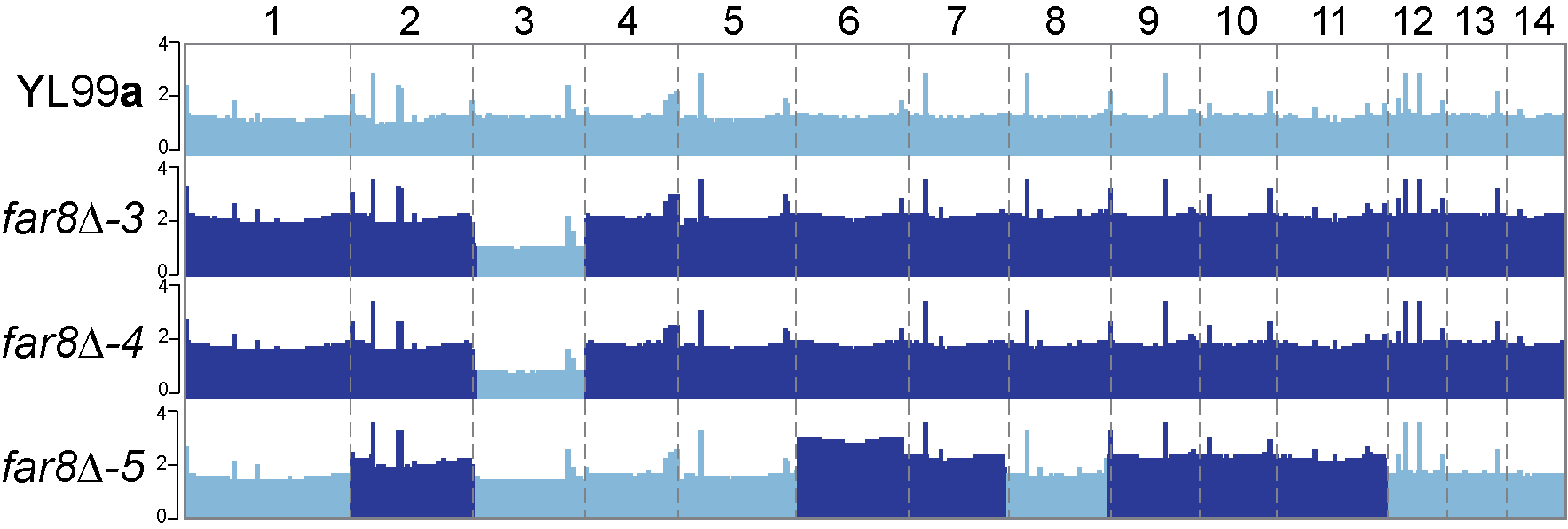
**A.**

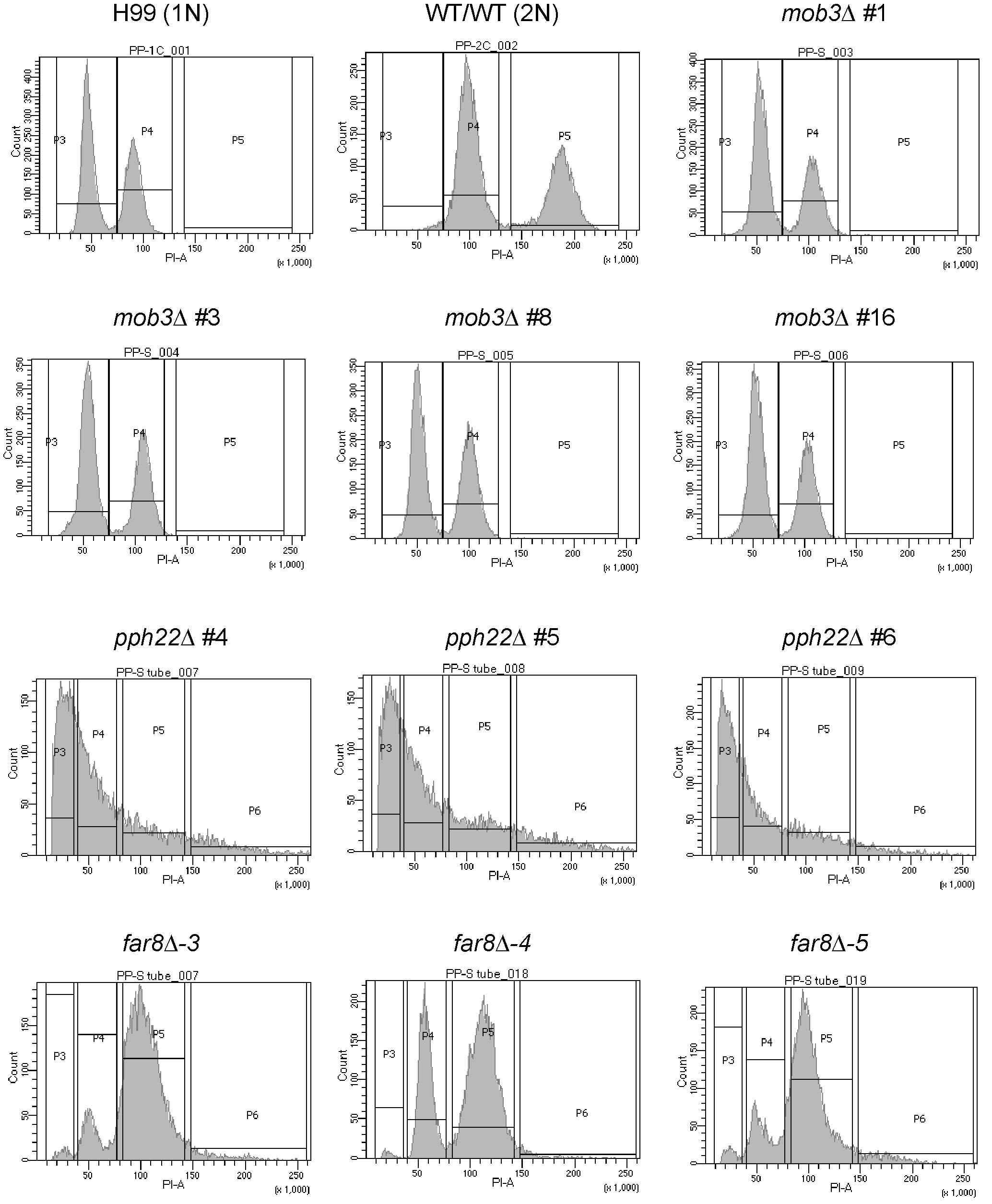
**B.**

**S4 Fig. Whole-genome sequencing and FACS analysis of *far8*Δ, *pph22*Δ, and *mob3*Δ mutants.**

(A) Read depth analyses of YL99**a** wild-type and *far8*Δ mutants showed duplications of the majority of chromosomes in the mutants, similar to what was observed in H99 *far8*Δ strains. *far8*Δ*-5* appeared to have an additional copy of chromosome 6. (B) FACS analyses of *mob3*Δ, *pph22*Δ, and *far8*Δ strains to determine ploidy, compared to wild-type 1N (H99) and 2N (CnLC6683) controls. Cells were stained with propidium iodide and DNA amount was quantitatively measured. *mob3*Δ mutants are haploid, with two peaks resembling H99 (1N). *pph22*Δ isolates only show one peak, suggesting cell cycle defects. YL99**a** *far8*Δ isolates had peaks resembling the wild-type 2N control, suggesting they are diploid.
