## Supplemental figures for "The *Cryptococcus neoformans* STRIPAK complex controls genome stability, sexual development, and virulence": S4 Table. Shared DEGs among mutant groups.docx

**S4 Table. Shared upregulated and downregulated genes among the four mutant groups**
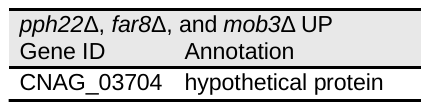

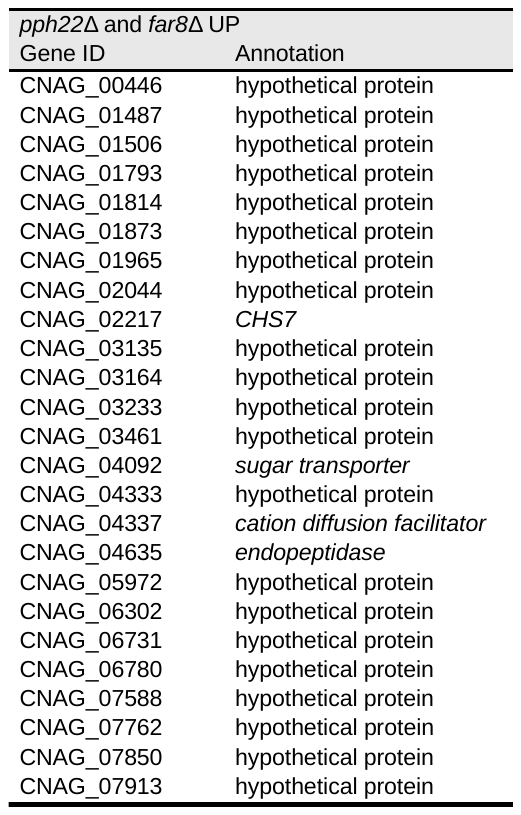

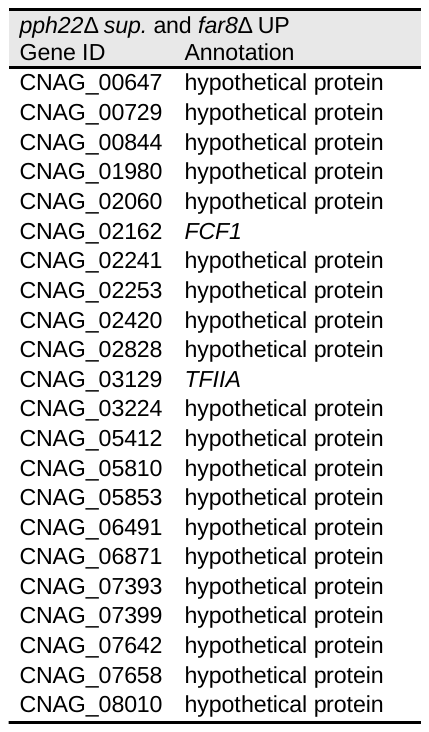
**.**

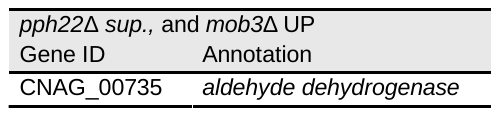

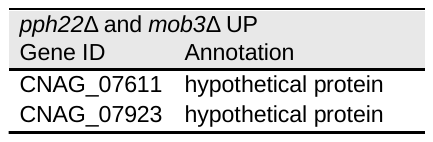

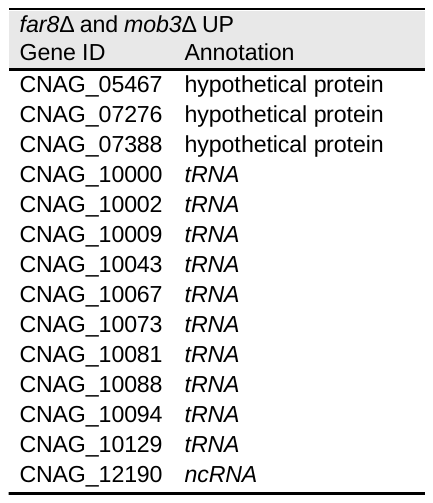

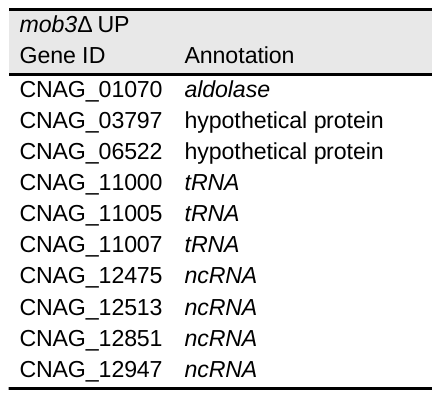

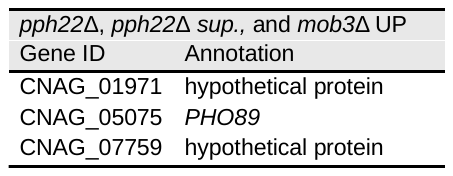

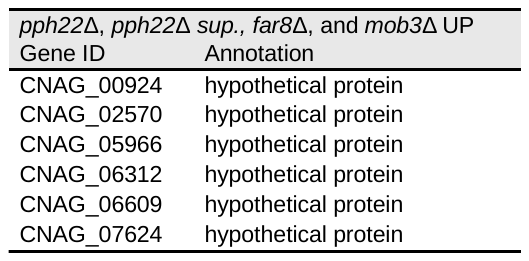

| *pph22*Δ, *pph22*Δ *sup.* and *far8*Δ UP | |  |  |
| --- | --- | --- | --- |
| Gene ID | Annotation |  |  |
| CNAG_00033 | hypothetical protein | CNAG_02122 | *NCS6* |
| CNAG_00177 | hypothetical protein | CNAG_02155 | hypothetical protein |
| CNAG_00359 | hypothetical protein | CNAG_02159 | hypothetical protein |
| CNAG_00430 | hypothetical protein | CNAG_02184 | hypothetical protein |
| CNAG_00643 | hypothetical protein | CNAG_02195 | *CDC6* |
| CNAG_00795 | hypothetical protein | CNAG_02220 | hypothetical protein |
| CNAG_00813 | hypothetical protein | CNAG_02339 | hypothetical protein |
| CNAG_00841 | hypothetical protein | CNAG_02441 | hypothetical protein |
| CNAG_01500 | *taurine dioxygenase* | CNAG_02582 | hypothetical protein |
| CNAG_01796 | hypothetical protein | CNAG_02586 | *sugar transporter* |
| CNAG_01806 | hypothetical protein | CNAG_02660 | hypothetical protein |
| CNAG_01901 | hypothetical protein | CNAG_02694 | hypothetical protein |
| CNAG_01953 | hypothetical protein | CNAG_03107 | hypothetical protein |
| CNAG_01977 | hypothetical protein | CNAG_03181 | hypothetical protein |
| CNAG_02066 | hypothetical protein | CNAG_03474 | *drug efflux protein* |
| CNAG_02082 | *DAD1* | CNAG_03519 | *FDC1* |
| CNAG_02083 | *SIT2* | CNAG_03717 | hypothetical protein |
| CNAG_03831 | hypothetical protein | CNAG_06259 | *MFS transporter* |
| CNAG_03835 | *DAD3* | CNAG_06324 | *zinc finger protein* |
| CNAG_03980 | hypothetical protein | CNAG_06341 | hypothetical protein |
| CNAG_04126 | hypothetical protein | CNAG_06538 | *MFS transporter* |
| CNAG_04385 | hypothetical protein | CNAG_06911 | hypothetical protein |
| CNAG_04919 | hypothetical protein | CNAG_07306 | hypothetical protein |
| CNAG_05184 | *glycosyl transferase* | CNAG_07391 | hypothetical protein |
| CNAG_05238 | hypothetical protein | CNAG_07471 | hypothetical protein |
| CNAG_05305 | hypothetical protein | CNAG_07644 | hypothetical protein |
| CNAG_05670 | hypothetical protein | CNAG_07648 | hypothetical protein |
| CNAG_05731 | *glyoxal oxidase* | CNAG_07843 | hypothetical protein |
| CNAG_05801 | hypothetical protein | CNAG_07868 | hypothetical protein |
| CNAG_05910 | hypothetical protein | CNAG_07998 | hypothetical protein |
| CNAG_05911 | hypothetical protein | CNAG_08000 | hypothetical protein |
| CNAG_06099 | hypothetical protein | CNAG_08006 | hypothetical protein |

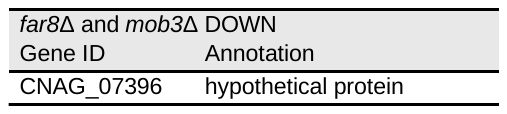

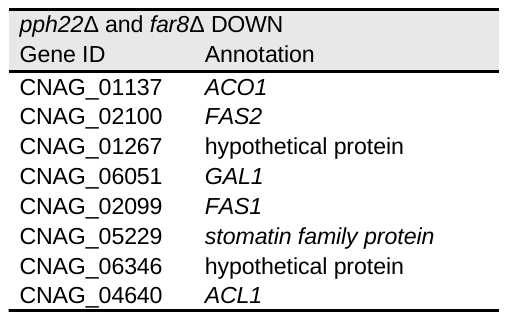

| *mob3*Δ DOWN | | |  |  |
| --- | --- | --- | --- | --- |
| Gene ID | | Annotation |  |  |
| CNAG_02489 | *ADH5* | | CNAG_07742 | hypothetical protein |
| CNAG_03460 | *phosphoglycerate dehydrogenase* | | CNAG_07895 | hypothetical protein |
| CNAG_04629 | *MOB3* | | CNAG_07910 | hypothetical protein |
| CNAG_05099 | hypothetical protein | | CNAG_07916 | hypothetical protein |
| CNAG_05331 | hypothetical protein | | CNAG_07918 | hypothetical protein |
| CNAG_05335 | hypothetical protein | | CNAG_10007 | *tRNA* |
| CNAG_06809 | *IKS1* | | CNAG_10012 | *tRNA* |
| CNAG_06877 | hypothetical protein | | CNAG_10014 | *tRNA* |
| CNAG_07306 | hypothetical protein | | CNAG_10019 | *tRNA* |
| CNAG_07454 | hypothetical protein | | CNAG_10022 | *tRNA* |
| CNAG_07496 | hypothetical protein | | CNAG_10031 | *tRNA* |
| CNAG_07643 | hypothetical protein | | CNAG_10039 | *tRNA* |
| CNAG_07705 | hypothetical protein | | CNAG_10040 | *tRNA* |
| CNAG_07713 | hypothetical protein | | CNAG_10056 | *tRNA* |
| CNAG_10066 | *tRNA* | | CNAG_10133 | *tRNA* |
| CNAG_10084 | *tRNA* | | CNAG_10140 | *tRNA* |
| CNAG_10086 | *tRNA* | | CNAG_10147 | *tRNA* |
| CNAG_10112 | *tRNA* | | CNAG_12106 | *tRNA* |
| CNAG_10117 | *tRNA* | | CNAG_12144 | *tRNA* |
| CNAG_10122 | *tRNA* | | CNAG_12366 | *tRNA* |
| CNAG_10126 | *tRNA* | | CNAG_12799 | *tRNA* |

| *pph22*Δ, *pph22*Δ *sup.,* and *far8*Δ DOWN | |  | |  |
| --- | --- | --- | --- | --- |
| Gene ID | Annotation |  |  | |
| CNAG_01464 | *FHB1* | CNAG_00727 | | *MMT1* |
| CNAG_01944 | hypothetical protein | CNAG_03333 | | *cytoplasmic protein* |
| CNAG_06388 | hypothetical protein | CNAG_04981 | | *CAT1* |
| CNAG_06298 | hypothetical protein | CNAG_02814 | | *GUT2* |
| CNAG_06207 | hypothetical protein | CNAG_07745 | | *ADH1* |
| CNAG_06297 | hypothetical protein | CNAG_07695 | | *gamma-aminobutyric acid transporter* |
| CNAG_00162 | *AOX1* | CNAG_06290 | | *SNF3* |
| CNAG_02542 | *IRK2* | CNAG_06868 | | *phosphopyruvate hydratase* |
| CNAG_04027 | hypothetical protein | CNAG_01155 | | *GUT1* |
| CNAG_00453 | *RCF2* | CNAG_03128 | | *gamma-glutamyltransferase* |
| CNAG_04757 | *MRX3* | CNAG_07347 | | *HSP104* |
| CNAG_04659 | *PDC1* | CNAG_04587 | | hypothetical protein |
| CNAG_02986 | *YSA1* | CNAG_01621 | | hypothetical protein |
| CNAG_06923 | *XFP2* | CNAG_04926 | | hypothetical protein |
| CNAG_06294 | hypothetical protein | CNAG_07877 | | hypothetical protein |
| CNAG_04217 | *PCK1* | CNAG_00686 | | hypothetical protein |
| CNAG_01558 | *chlorophyll synthesis pathway protein* | CNAG_02722 | | *aldose reductase* |
| CNAG_00115 | *chlorophyll synthesis pathway protein* |  | |  |

| *pph22*Δ and *pph22*Δ *sup.* DOWN | |  |  |
| --- | --- | --- | --- |
| Gene ID | Annotation |  |  |
| CNAG_00337 | hypothetical protein | CNAG_02041 | hypothetical protein |
| CNAG_00539 | *MDR membrane transporter* | CNAG_02177 | *PPH22* |
| CNAG_00559 | hypothetical protein | CNAG_02460 | *HEM13* |
| CNAG_00915 | hypothetical protein | CNAG_02527 | *MDR transporter* |
| CNAG_01013 | *SWD3* | CNAG_02577 | *oxidoreductase* |
| CNAG_01493 | hypothetical protein | CNAG_02857 | hypothetical protein |
| CNAG_01574 | hypothetical protein | CNAG_03259 | hypothetical protein |
| CNAG_01601 | *ATG15* | CNAG_03434 | *substrate transporter* |
| CNAG_01636 | *cyclophilin-like protein* | CNAG_03502 | *ALR2* |
| CNAG_01761 | hypothetical protein | CNAG_04085 | *oxidoreducatse* |
| CNAG_04394 | hypothetical protein | CNAG_06230 | hypothetical protein |
| CNAG_04551 | hypothetical protein | CNAG_06371 | *GUD1* |
| CNAG_04756 | hypothetical protein | CNAG_06525 | *fungal transcription factor* |
| CNAG_04773 | hypothetical protein | CNAG_06546 | hypothetical protein |
| CNAG_04816 | hypothetical protein | CNAG_06705 | hypothetical protein |
| CNAG_04920 | *HXT15* | CNAG_06816 | hypothetical protein |
| CNAG_05397 | hypothetical protein | CNAG_06936 | *beta glucosidase* |
| CNAG_05693 | hypothetical protein | CNAG_06956 | hypothetical protein |
| CNAG_05874 | hypothetical protein | CNAG_07317 | hypothetical protein |
| CNAG_05931 | *BTN1* | CNAG_07654 | hypothetical protein |
| CNAG_05938 | hypothetical protein | CNAG_07816 | *GOR1* |
| CNAG_06031 | *KRE63* | CNAG_07824 | hypothetical protein |
| CNAG_06188 | hypothetical protein |  |  |
