## Supplemental figures for "The *Cryptococcus neoformans* STRIPAK complex controls genome stability, sexual development, and virulence": S5 Fig. Serial diluton of pph22 diploid mutant.docx

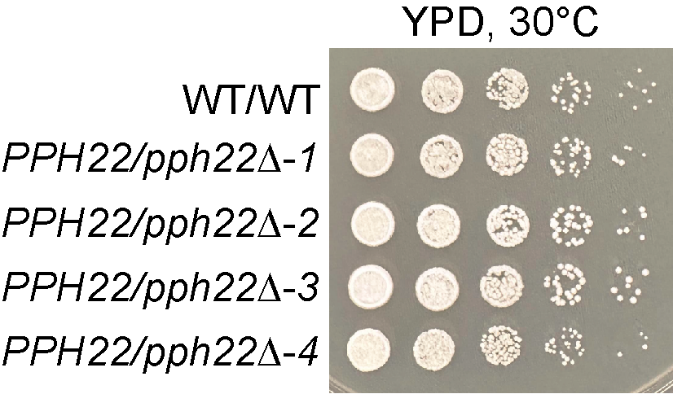

**S5 Fig. Serial dilution assay of *PPH22/pph22*Δ** **heterozygous mutant and wild-type diploid strains.**

Wild-type (CnLC6683) and *PPH22/pph22*Δ diploid strains were pre-grown in YPD liquid cultures overnight at 30°C. Cells were then serially diluted and spotted onto YPD medium and incubated at 30°C. The *PPH22/pph22*Δ strains did not exhibit any observable growth defects compared to the wild-type control strain.
