## Supplemental figures for "The *Cryptococcus neoformans* STRIPAK complex controls genome stability, sexual development, and virulence": S5 Table. List of virulence related genes.docx

**S5 Table. Genes known or predicted to be involved in capsule production, melanization, or important for virulence of *C. neoformans.***

| **Gene ID** | **Category** | **Gene name** | **Source/Reference** |
| --- | --- | --- | --- |
| CNAG_00124 | Capsule | *CAS32* | FungiDB phenotype search |
| CNAG_00130 | Melanin | *RCK2* | PLoS ONE 6(4): e18769 |
| CNAG_00247 | Virulence | *LYS9* | FungiDB phenotype search |
| CNAG_00268 | Thermotolerance | *ILV2* | FungiDB phenotype search |
| CNAG_00293 | Cell growth | *RAS1* | Eukaryot Cell. 2010 Mar;9(3):360-78. |
| CNAG_00375 | Thermotolerance | *GCN5* | Eukaryot Cell. 2010 Aug;9(8):1193-202. |
| CNAG_00396 | Virulence | *PKA1* | Mol Cell Biol. 2001 May;21(9):3179-91 |
| CNAG_00460 | Virulence | *LIV1* | FungiDB phenotype search |
| CNAG_00531 | Virulence | *ENA1* | FungiDB phenotype search |
| CNAG_00570 | Capsule | *PKR1* | Mol Cell Biol. 2001 May;21(9):3179-91. |
| CNAG_00600 | Capsule | *CAP60* | Infect Immun. 1998 May;66(5):2230-6 |
| CNAG_00701 | Capsule | *CAS31* | FungiDB phenotype search |
| CNAG_00721 | Capsule | *CAP59* | Mol Cell Biol. 1994 Jul;14(7):4912-9 |
| CNAG_00746 | Capsule | *CAS35* | FungiDB phenotype search |
| CNAG_00769 | Melanin | *PBS2* | Mol Biol Cell. 2005 May;16(5):2285-300. |
| CNAG_00797 | Virulence |  | FungiDB phenotype search |
| CNAG_00815 | Melanin | *SIT1* | Microbiology. 2007 Jan;153(Pt 1):29-41. |
| CNAG_00888 | Thermotolerance | *CNB1* | FungiDB phenotype search |
| CNAG_00984 | Melanin |  | FungiDB phenotype search |
| CNAG_00996 | Melanin | *PMT4* | PLoS One. 2009 Jul 27;4(7):e6321. |
| CNAG_01102 | Melanin |  | FungiDB phenotype search |
| CNAG_01106 | Capsule | *VPH1* | Mol Microbiol. 2001 Nov;42(4):1121-31. |
| CNAG_01377 | Melanin | *UBP13* | FungiDB phenotype search |
| CNAG_01431 | Virulence | *HOB1* | FungiDB phenotype search |
| CNAG_01446 | Thermotolerance | *HSP12* | FungiDB phenotype search |
| CNAG_01464 | Virulence | *FHB1* | FungiDB phenotype search |
| CNAG_01523 | Virulence | *HOG1* | Mol Biol Cell. 2005 May;16(5):2285-300. |
| CNAG_01530 | Thermotolerance | *MGA2* | Eukaryot Cell. 2004 Oct;3(5):1249-60. |
| CNAG_01563 | Virulence | *HDA1* | FungiDB phenotype search |
| CNAG_01626 | Virulence | *ADA2* | PLoS Pathog. 2011 Dec;7(12):e1002411 |
| CNAG_01654 | Capsule | *CAS34* | FungiDB phenotype search |
| CNAG_01845 | Virulence | *PKC1* | Eukaryot Cell. 2008 Oct;7(10):1685-98. |
| CNAG_01983 | Virulence | *RCV1* | FungiDB phenotype search |
| CNAG_02008 | Capsule | *OVA1* | PLoS Pathog. 2007 Mar;3(3):e42. |
| CNAG_02036 | Capsule | *CAS4* | FungiDB phenotype search |
| CNAG_02057 | Virulence | *URE6* | FungiDB phenotype search |
| CNAG_02357 | Virulence | *MKK2* | Mol Microbiol. 2005 Oct;58(2):393-408. |
| CNAG_02511 | Melanin | *CPK1* | FungiDB phenotype search |
| CNAG_02531 | Melanin | *CPK2* | FungiDB phenotype search |
| CNAG_02579 | Virulence | *AVC1* | FungiDB phenotype search |
| CNAG_02702 | Virulence | *CLC-A* | FungiDB phenotype search |
| CNAG_02827 | Virulence | *RUB1* | FungiDB phenotype search |
| CNAG_02867 | Virulence | *PLC1* | Mol Microbiol. 2008 Aug;69(4):809-26. |
| CNAG_02885 | Capsule | *CAP64* | Infect Immun. 1996 Jun;64(6):1977-83 |
| CNAG_02930 | Virulence | *LIV10* | FungiDB phenotype search |
| CNAG_02958 | Melanin | *CFO2* | FungiDB phenotype search |
| CNAG_03120 | Melanin | *AGS1* | FungiDB phenotype search |
| CNAG_03158 | Capsule | *CMT1* | FungiDB phenotype search |
| CNAG_03202 | Virulence | *CAC1* | Eukaryot Cell. 2002 Feb;1(1):75-84. |
| CNAG_03409 | Melanin | *SKN7* | Mol Biol Cell. 2006 Jul;17(7):3122-35. |
| CNAG_03464 | Melanin | *LAC2* | Eukaryot Cell 2005 4(1) 190-201 |
| CNAG_03465 | Melanin | *LAC1* | J. Exp. Med. 1996 184:377–386. |
| CNAG_03482 | Virulence | *TSA1* | FungiDB phenotype search |
| CNAG_03582 | Capsule | *RIM20* | FungiDB phenotype search |
| CNAG_03627 | Virulence | *CPA1* | FungiDB phenotype search |
| CNAG_03628 | Virulence |  | FungiDB phenotype search |
| CNAG_03644 | Capsule | *CAS3* | FungiDB phenotype search |
| CNAG_03670 | Virulence | *IRE1* | PLoS Pathog. 2011 Aug;7(8):e1002177 |
| CNAG_03695 | Capsule | *CAS41* | FungiDB phenotype search |
| CNAG_03754 | Melanin | *ISP1* | FungiDB phenotype search |
| CNAG_03765 | Thermotolerance | *TPS2* | FungiDB phenotype search |
| CNAG_03818 | Melanin | *SSK1* | Mol Biol Cell. 2006 Jul;17(7):3122-35. |
| CNAG_04090 | Virulence | *ATF1* | Genetics. 2010 Aug;185(4):1207-19. |
| CNAG_04215 | Melanin | *MET3* | Microbiology. 2002 Aug;148(Pt 8):2617-25. |
| CNAG_04243 | Thermotolerance | *CDC24* | FungiDB phenotype search |
| CNAG_04282 | Melanin | *MPK2* | FungiDB phenotype search |
| CNAG_04320 | Capsule | *CPS1* | FungiDB phenotype search |
| CNAG_04388 | Thermotolerance | *SOD2* | Eukaryot Cell. 2005 Jan;4(1):46-54. |
| CNAG_04505 | Virulence | *GPA1* | Eukaryot Cell. 2004 Dec;3(6):1476-91. |
| CNAG_04514 | Thermotolerance | *MPK1* | Mol Microbiol. 2003 Jun;48(5):1377-87 |
| CNAG_04755 | Thermotolerance | *BCK1* | Mol Microbiol. 2005 Oct;58(2):393-408. |
| CNAG_04796 | Thermotolerance | *CNA1* | FungiDB phenotype search |
| CNAG_04864 | Virulence | *CIR1* | PLoS Biol. 2006 Nov;4(12):e410. |
| CNAG_04969 | Virulence | *UGD1* | Eukaryot Cell. 2004 Dec;3(6):1601-8. |
| CNAG_05063 | Virulence | *SSK2* | Eukaryot Cell. 2007 Dec;6(12):2278-89. |
| CNAG_05081 | Virulence | *PDE1* | Eukaryot Cell, Dec. 2005, 4(12) 1971–1981 |
| CNAG_05218 | Capsule | *ACA1* | Eukaryot Cell. 2004 Dec;3(6):1476-91 |
| CNAG_05292 | Thermotolerance | *TPS1* | FungiDB phenotype search |
| CNAG_05293 | Capsule |  | FungiDB phenotype search |
| CNAG_05348 | Virulence | *CDC42* | Mol Microbiol. 2010 Feb;75(3):763-80 |
| CNAG_05422 | Virulence | *LIV11* | FungiDB phenotype search |
| CNAG_05431 | Virulence | *RIM101* | PLoS Pathog. 2010 Feb 19;6(2):e1000776. |
| CNAG_05465 | Virulence | *GIB2* | FungiDB phenotype search |
| CNAG_05540 | Virulence | *URE1* | FungiDB phenotype search |
| CNAG_05583 | Virulence | *GCS1* | FungiDB phenotype search |
| CNAG_05741 | Capsule |  | FungiDB phenotype search |
| CNAG_05817 | Virulence | *GMT1* | FungiDB phenotype search |
| CNAG_05968 | Virulence | *CDC420* | FungiDB phenotype search |
| CNAG_06016 | Capsule | *CAP6* | FungiDB phenotype search |
| CNAG_06085 | Virulence | *PLB1* | FungiDB phenotype search |
| CNAG_06134 | Thermotolerance | *BZP1* | PLoS Pathog. 2011 Aug;7(8):e1002177 |
| CNAG_06165 | Virulence |  | FungiDB phenotype search |
| CNAG_06208 | Virulence |  | FungiDB phenotype search |
| CNAG_06241 | Melanin | *CFO1* | FungiDB phenotype search |
| CNAG_06301 | Virulence | *SCH9* | Curr Genet. 2004 Nov;46(5):247-55. |
| CNAG_06464 | Virulence | *LIV7* | FungiDB phenotype search |
| CNAG_06465 | Virulence | *CDC50* | FungiDB phenotype search |
| CNAG_06469 | Virulence | *APT1* | FungiDB phenotype search |
| CNAG_06552 | Melanin | *SNF1* | Fungal Genet Biol. 2010 Dec; 47(12):994-1000. |
| CNAG_06762 | Capsule | *GAT204* | FungiDB phenotype search |
| CNAG_07347 | Virulence |  | FungiDB phenotype search |
| CNAG_07460 | Virulence |  | FungiDB phenotype search |
| CNAG_07470 | Virulence | *PDE2* | Eukaryot Cell, Dec. 2005, 4(12) 1971–1981 |
| CNAG_07554 | Capsule | *CAP10* | J Bacteriol. 1999 Sep;181(18):5636-43. |
| CNAG_07744 | Virulence | *PIK1* | FungiDB phenotype search |
| CNAG_07865 | Melanin |  | FungiDB phenotype search |
