## Supplemental figures for "The *Cryptococcus neoformans* STRIPAK complex controls genome stability, sexual development, and virulence": S6 Fig. Structure prediction of PP2A.docx

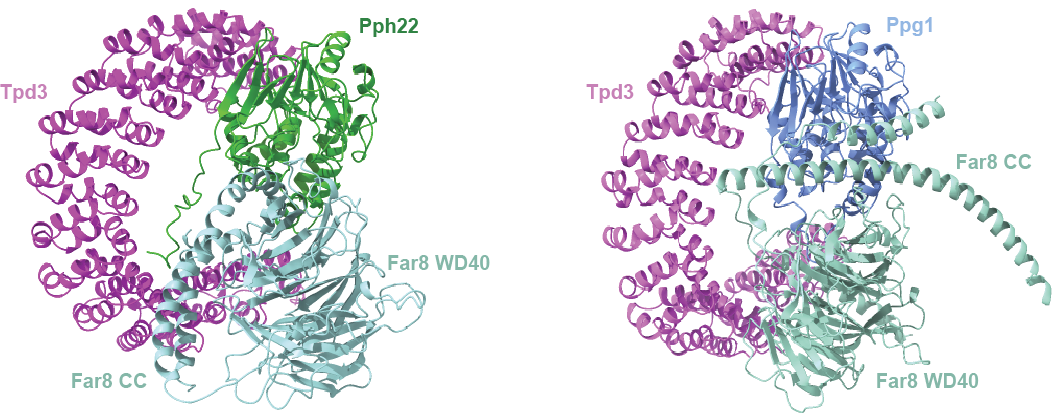

**S6 Fig. AlphaFold structure prediction of the PP2A heterotrimeric enzyme.** Shown with the scaffold subunit Tpd3, regulatory B’’’ subunit Far8, and either Pph22 (left) or Ppg1 (right) as the catalytic subunit. The WD40 and coiled-coil (CC) domains of Far8 are labeled.
