## Supplemental figures for "The *Cryptococcus neoformans* STRIPAK complex controls genome stability, sexual development, and virulence": S6 Table. GO tern enrichment.docx

**S6 Table. Gene ontology enrichment of molecular functions and biological processes represented in significant DEGs**

| GO term | Functional category | Fold enrichment | log2 FE | Enrichment FDR |
| --- | --- | --- | --- | --- |
| GO:0006414 | Translation elongation | 3.34 | 1.739848 | 3.284631 |
| GO:0016407 | Acetyltransferase | 3.27 | 1.709291 | 2.757205 |
| GO:0004129 | Cytochrome-c oxidase activity | 2.97 | 1.570463 | 2.743503 |
| GO:0006006 | Glucose metabolic process | 2.86 | 1.516015 | 2.264532 |
| GO:0008652 | Amino acid biosynthesis | 2.67 | 1.41684 | 2.332097 |
| GO:0006413 | Translational initiation | 2.64 | 1.400538 | 2.849638 |
| GO:0006099 | TCA cycle | 2.54 | 1.344828 | 3.437084 |
| GO:006163 | Purine nucleotide metabolism | 2.54 | 1.344828 | 2.085595 |
| GO:0019843 | rRNA binding | 2.35 | 1.232661 | 2.116831 |
| GO:006521 | Amino acid metabolic process | 2.35 | 1.232661 | 2.116831 |
| GO:006119 | Oxidative phosphorylation | 2.29 | 1.195348 | 2.233152 |
| GO:0022857 | Transmembrane transporter | 2.26 | 1.176323 | 3.548123 |
| GO:0043603 | Amide metabolic process | 2.21 | 1.144046 | 13.64604 |
| GO:0020037 | Heme binding | 2.14 | 1.097611 | 2.830607 |
| GO:0003735 | Structural component of the ribosome | 2.01 | 1.013911 | 9.778289 |

GO enrichment: *pph22*Δ

GO enrichment: *far8*Δ

| GO term | Functional category | Fold enrichment | log2 FE | Enrichment FDR |
| --- | --- | --- | --- | --- |
| GO:006090 | Pyruvate metabolism | 2.39 | 1.257010618 | 2.046103 |
| GO:0051287 | NAD binding | 2.52 | 1.333423734 | 2.918826 |
| GO:0022857 | Transmembrane transport | 2.55 | 1.350497247 | 2.508556 |
| GO:0020037 | Heme binding | 3.79 | 1.922197848 | 2.770937 |
| GO:0051213 | Dioxygenase activity | 3.87 | 1.952333566 | 3.164534 |
| GO:0044042 | Glucan metabolism | 3.87 | 1.952333566 | 2.791453 |
| GO:0016051 | carbohydrate biosynthetic process | 4.06 | 2.021479727 | 2.605937 |
| GO:0015976 | Carbon source utilization | 5.21 | 2.381283373 | 2.776075 |
| GO:0006637 | acyl-CoA biosynthesis | 6.77 | 2.759155834 | 2.170636 |
| GO:0006012 | Galactose metabolic process | 6.77 | 2.759155834 | 2.170636 |
| GO:0035384 | Thioester biosynthesis | 6.77 | 2.759155834 | 2.170636 |
| GO:0052646 | Alditol phosphate metabolism | 10.16 | 3.344828497 | 2.820776 |
| GO:0004312 | Fatty acid synthase | 13.54 | 3.759155834 | 2.264452 |
| GO:0006072 | Glycerol-3-phosphate metabolism | 13.54 | 3.759155834 | 2.264452 |
| GO:0005977 | Glycogen metabolism | 13.54 | 3.759155834 | 3.39828 |

GO enrichment: *mob3*Δ

| GO term | Functional category | Fold enrichment | log2 FE | Enrichment FDR |
| --- | --- | --- | --- | --- |
| GO:1990456 | Mitochondria-ER membrane tethering | 213.29 | 7.736673 | 3.067318 |
| GO:0070096 | Mitochondrial outer membrane translocase | 213.29 | 7.736673 | 4.185642 |
| GO:0006817 | Phosphate ion transport | 106.65 | 6.73674 | 3.640723 |
| GO:0009063 | Amino acid catabolism | 71.10 | 6.151778 | 3.493899 |
| GO:0010043 | Response to zinc ion | 35.55 | 5.151778 | 4.702312 |
| GO:0016787 | Hydrolase activity | 26.66 | 4.736605 | 3.48253 |
| GO:0008643 | Carbohydrate transport | 21.33 | 4.414812 | 2.627976 |
| GO:0003678 | DNA helicase activity | 12.00 | 3.584963 | 2.724238 |
| GO:0000302 | Response to reactive oxygen species | 11.23 | 3.489286 | 3.155433 |
| GO:0016491 | Oxidoreductase activity | 8.10 | 3.017922 | 3.412518 |
| GO:0003700 | DNA-binding transcription factor activity | 3.95 | 1.981853 | 2.755433 |
| GO:2001141 | Regulation of RNA biosynthesis | 3.68 | 1.879706 | 3.561817 |
| GO:0051252 | Regulation of RNA metabolism | 3.51 | 1.811471 | 2.014748 |
| GO:0010556 | Regulation of macromolecule biosynthesis | 3.32 | 1.731183 | 2.474042 |
| GO:0055085 | Transmembrane transport | 2.97 | 1.570463 | 2.412518 |
