## Supplemental figures for "The *Cryptococcus neoformans* STRIPAK complex controls genome stability, sexual development, and virulence": S7 Fig. Bilateral crosses of pph22 far8 and mob3 mutants.docx

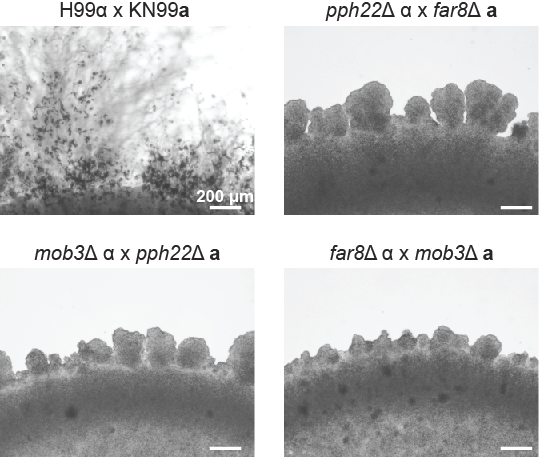

**S7 Fig. Bilateral crosses of** ***pph22*Δ*, far8*Δ*,* and *mob3*Δ mutants.**

Strains were grown in YPD overnight cultures before mixing cells in a 1:1 ratio for spotting onto MS medium. Each cross was spotted 10 times onto an MS plate, along with one H99α x KN99**a** wild-type control. Bilateral (mutant x mutant) crosses were incubated at room temperature (24°C) in the dark for 8 weeks. No signs of mating between *pph22*Δ*, far8*Δ*,* and *mob3*Δ strains, in both **a** and α mating types, were observed.
