## Supplemental figures for "The *Cryptococcus neoformans* STRIPAK complex controls genome stability, sexual development, and virulence": S8 Fig. Fluconazole Etest.docx

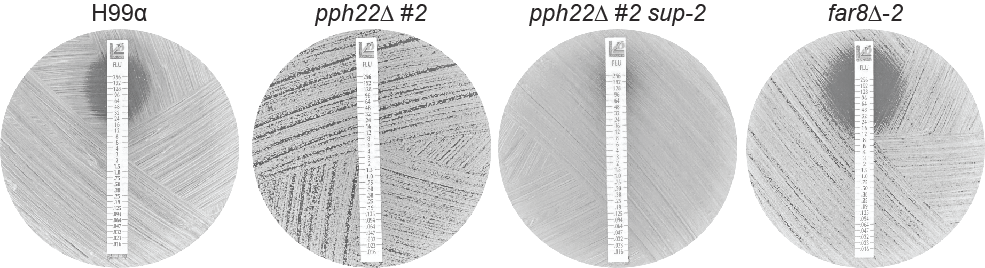

**S8 Fig. Fluconazole Etest of *pph22*Δ, *pph22*Δ *sup*, and *far8*Δ mutants**

The indicated strains were grown overnight in YPD media to saturation. Cells were spread onto YPD plates and allowed to dry before adding fluconazole Etest strips. Plates were incubated at 30°C and images were taken after 24 hours.
