## Supplemental figures for "The *Cryptococcus neoformans* STRIPAK complex controls genome stability, sexual development, and virulence": S9 Fig. FACS analysis of far8 mutants on RPMI.docx

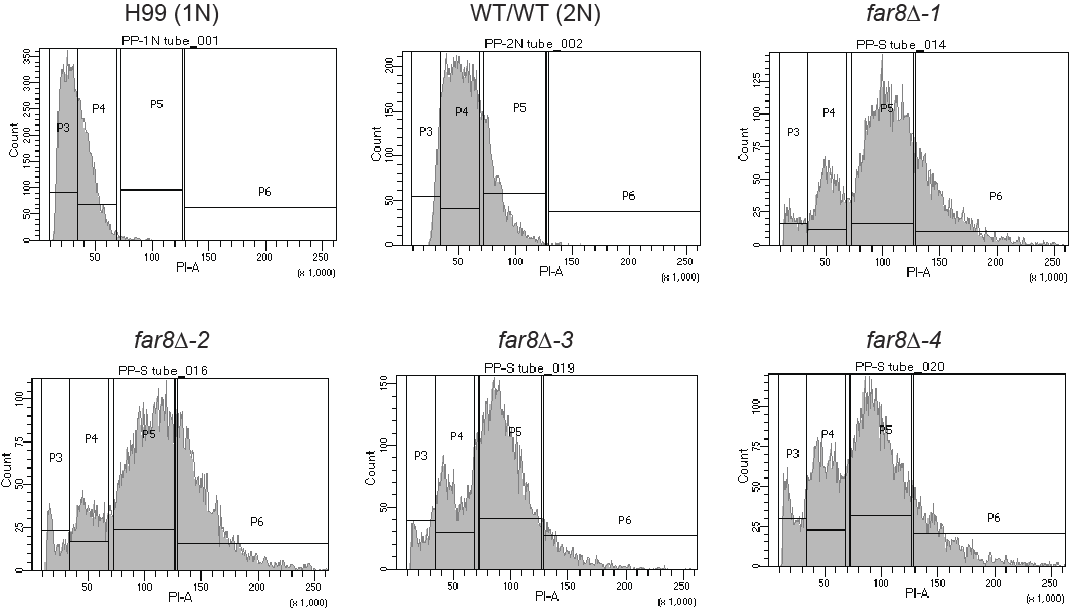

**S9 Fig. FACS analysis of *far8*Δ mutant strains on RPMI medium.**

*far8*Δ mutants and wild-type 1N (H99) and 2N (CnLC6683) control cells were grown on RPMI medium and stained with propidium iodide to quantify DNA amount. FACS profiles of *far8*Δ cells from RPMI are similar to those from YPD with two major peaks occurring at 2C and 4C, suggesting they are diploid.
