## Supplemental figures for "The *Cryptococcus neoformans* STRIPAK complex controls genome stability, sexual development, and virulence": S10 Fig. Phenotyping mob3 mutants.docx

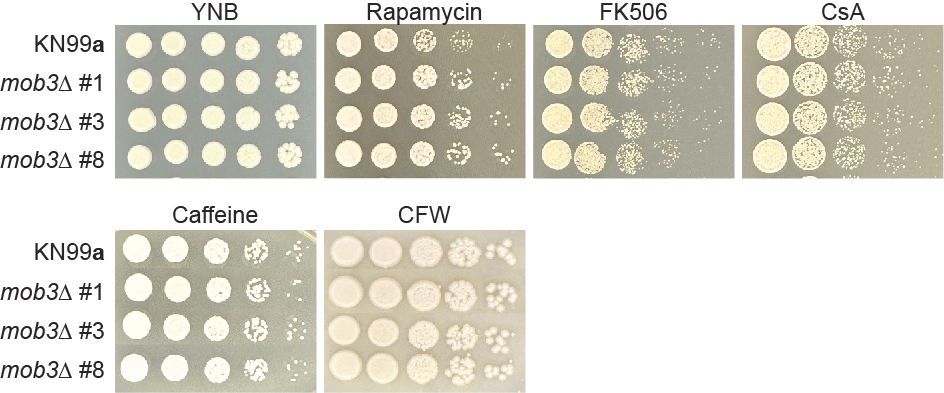

**S10 Fig. Phenotypic analyses of *mob3*Δ mutant strains.**

Serially diluted wild-type KN99**a** and isogenic *mob3*Δ mutants were spotted onto YNB, as well as YPD supplemented with rapamycin, FK506, cyclosporine A (CsA), caffeine, or calcofluor white (CFW). Plates were incubated at 30°C and images were taken after 3 days.
