## Supplemental figures for "The *Cryptococcus neoformans* STRIPAK complex controls genome stability, sexual development, and virulence": S11 Fig. Volcano plot of DEGs in pph22 suppressors.docx

**S11 Fig. Volcano plot of DEGs in *pph22*Δ *suppressor* mutants**

Log_2_ fold change is plotted against the -log_10_ *P* value for all DEGs showing significantly upregulated (red) and downregulated (blue) genes. The top 10 most significant genes are labeled.

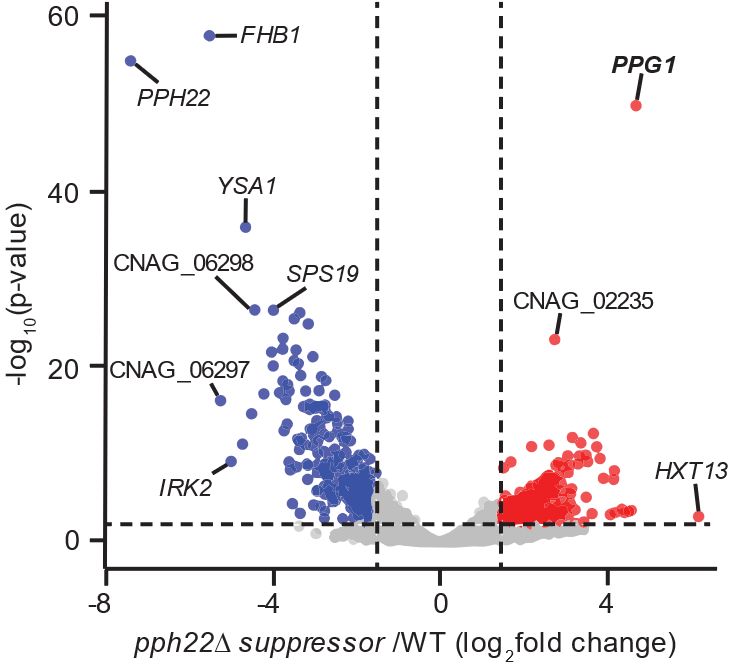
