## Supplemental figures for "The *Cryptococcus neoformans* STRIPAK complex controls genome stability, sexual development, and virulence": S12 Fig. Heatmaps of DEGs from RNAseq.docx

**S12 Fig. Heatmaps depicting hierarchical clustering of differential gene expression in *pph22*Δ (*p*Δ),** ***pph22*Δ *suppressor* (*p*Δs), *far8*Δ (*f*Δ), and *mob3*Δ (*m*Δ) strains compared to wild type (WT).**

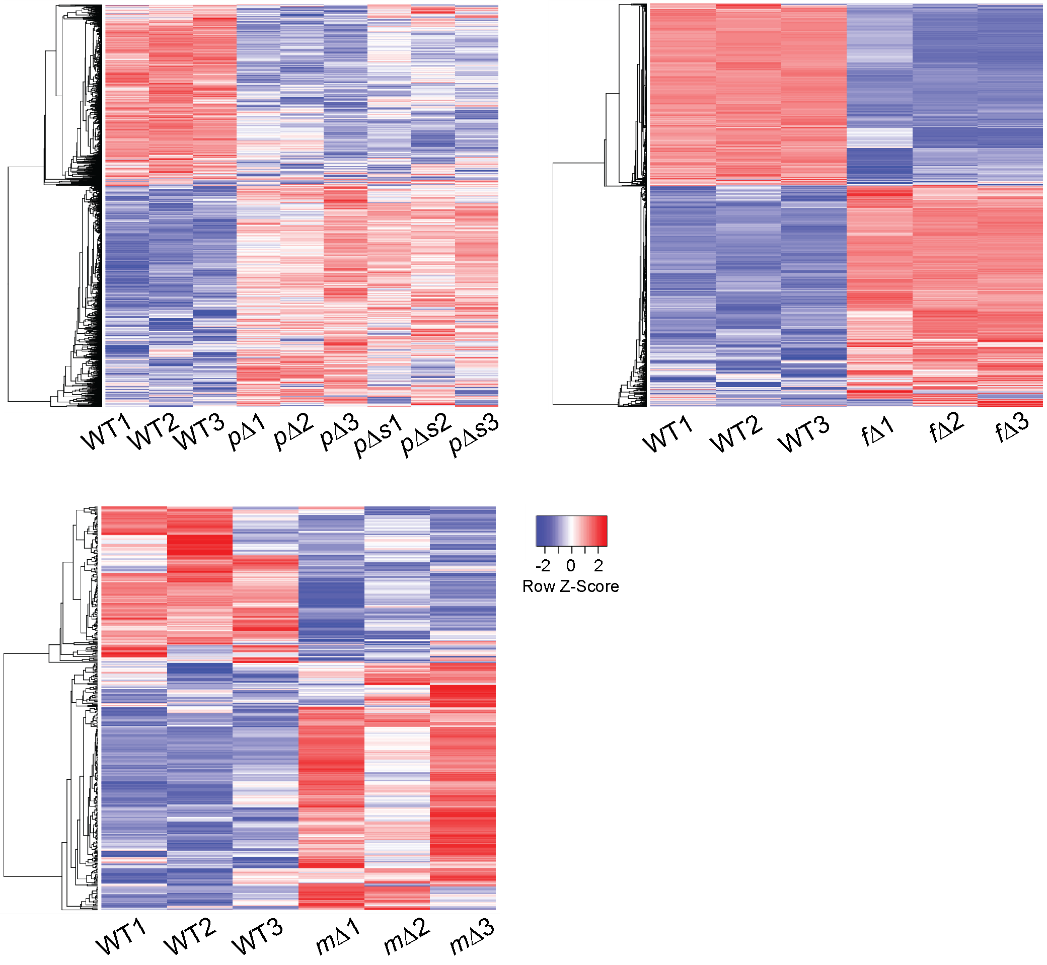
