## Supplemental figures for "The *Cryptococcus neoformans* STRIPAK complex controls genome stability, sexual development, and virulence": S13 Fig.Potential targets of STRIPAK from RNAseq.docx

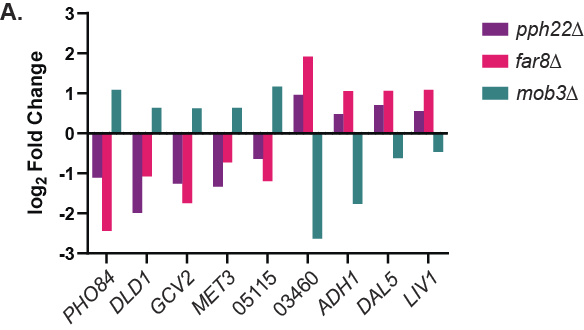

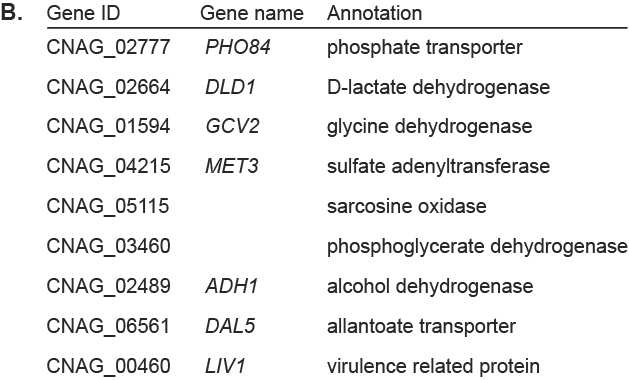

**S13 Fig. Potential direct or indirect targets of the STRIPAK complex.** (A) Expression (log_2_ fold change) of nine genes that are oppositely regulated in *mob3*Δ mutants compared to *pph22*Δ and *far8*Δ*.* (B) Table listing gene ID numbers of the identified genes and their annotations from FungiDB.
